# A unique *Toxoplasma gondii* haplotype under strong selection has accompanied domestic cats in their global expansion

**DOI:** 10.1101/2021.08.31.457674

**Authors:** Lokman Galal, Frédéric Ariey, Meriadeg Ar Gouilh, Marie-Laure Dardé, Azra Hamidović, Franck Letourneur, Franck Prugnolle, Aurélien Mercier

## Abstract

*Toxoplasma gondii* is a cyst-forming apicomplexan parasite of virtually all warm-blooded species, with all true cats (Felidae) as definitive hosts. It is the etiologic agent of toxoplasmosis, a disease causing substantial public health burden worldwide. Its wide range of host species and its global occurrence probably complicate the study of its evolutionary history, and conflicting scenarios have been proposed to explain its current global distribution. In this study, we analyse a global set of 156 genomes (including 105 new genomes) and we provide the first direct estimate of *T. gondii* mutation rate and the first estimate of its generation time. We elucidate how the evolution of *T. gondii* populations is intimately linked to the major events that have punctuated the recent history of cats. We show that a unique haplotype —whose length represents only 0.16% of the whole *T. gondii* genome— is common to all domestic *T. gondii* strains worldwide and has accompanied wild cats (*Felis silvestris*) during their emergence from the wild to domestic settlements, their dispersal in the Old World and their recent expansion to the Americas in the last six centuries. By combining environmental and functional data to selection inference tools, we show that selection of this domestic haplotype is most parsimoniously explained by its role in initiation of sexual reproduction of *T. gondii* in domestic cats.

## INTRODUCTION

*Toxoplasma gondii* is a zoonotic protozoan that has spread globally. This apicomplexan parasite infects all warm-blooded species including humans, and its wide range of host species suggests multiple routes for short and long-distance parasite migrations (Galal et al., 2019). *T. gondii* is found in approximately 30% of the human population and is the etiologic agent of toxoplasmosis, a disease causing a substantial public health burden worldwide (Montoya and Liesenfeld, 2004). Infection with *T. gondii* has been long considered as benign or even asymptomatic except for certain risk groups like the developing foetus in case of congenital infection —with 200,000 new cases of congenital toxoplasmosis each year (Torgerson and Mastroiacovo, 2013)— and immunocompromised patients, for whom toxoplasmosis can have severe health consequences either during primo-infection or reactivation. However, certain *T. gondii* populations have been associated with severe toxoplasmosis in immunocompetent individuals (Carme et al., 2009; Pomares et al., 2018; Schumacher et al., 2020). More importantly, an increasing number of epidemiological studies suggest that chronic infection with *T. gondii* is associated with a wide variety of neuropsychiatric disorders, substantially raising the public health importance of this global and highly prevalent parasite (Milne et al., 2020). Given gaps in both the current preventive (no vaccine available for humans) and therapeutic strategies (Innes et al., 2019; Konstantinovic et al., 2019), active research to discover new ways to target this clinically important protozoan are still needed.

*T. gondii* hosts get infected after ingestion of oocysts shed into the environment by contaminated faeces of felids and develop persistent tissue-cysts. Another source of infection for human and other meat-consuming species is the ingestion of raw or undercooked meat from animals harbouring infective tissue-cysts. In the domestic environment cats and rodents are considered as the most significant reservoirs for human infection, since life cycle completion relies mainly on transmission between these two categories of animal hosts, the rodents being the main prey of cats (Müller and Howard, 2016; Galal et al., 2020). Sexual recombination is possible when two different strains are found simultaneously in the cat’s gut. For this to occur a cat has to ingest within a few hours two prey infected with different strains. There is therefore a time barrier for recombination to occur, or alternatively the cat has to ingest a single prey infected with two different strains (mixed infection), a rare event in nature given that intermediate hosts develop immunity to new infections following their first infection.

From a genetic point of view, the population structure of *T. gondii* is characterized by contrasting patterns of strain diversity mainly varying according to geographical origin and ecotype. This diversity has mainly been identified based on the analysis of microsatellite markers (MS), or Restriction Fragment Length Polymorphism (RFLP) (refer to Supplementary Table 2 for correspondence MS and RFLP designations of lineages or genotypes). In the Old World (Africa, Asia and Europe), most *T. gondii* isolates from humans, domestic animals and wild fauna belong to few intercontinental clonal lineages: type I, type II, type III, Africa 1 (also designated as BrI) and Africa 4 (Shwab et al., 2014; Galal et al., 2019). Few other clonal lineages have been described in certain countries such as Chinese 1 in China (Chaichan et al., 2017) and Africa 3 in Gabon (Mercier et al., 2010), and strains not belonging to these major lineages were rarely isolated (Galal et al., 2018). This genetic evidence argues that sexual recombination between different strains is not frequent in these regions. In most regions of the Old World, populations of wild felids have undergone massive decline (Goodrich et al., 2015; Bauer et al., 2016), leaving domestic cats as virtually the only shedders of oocysts capable of infecting domestic animals and humans, before spreading over long distances via waterways to reach wildlife (Gotteland et al., 2014; VanWormer et al., 2013). In the New World (North and South America), the most common Old World clonal lineages (type I, II, III and Africa 1) are also found, in sympatry with a substantial diversity of local clonal lineages strains and non-clonal strains specific to South or North America (Shwab et al., 2014; Jiang et al., 2018). In contrast to the pattern observed in the Old World, the genotypic composition of strains from wildlife differs importantly from strains commonly isolated in the domestic environment (Mercier et al., 2011; Jiang et al., 2018). Globally, wild *T. gondii* populations — isolated from wild animals or from humans in contact with wildlife and genetically distinct from domestic *T. gondii* populations— are associated to biotopes where the presence of wild felids is well-established (Carme et al., 2009; Khan et al., 2011; Mercier et al., 2011; L Galal et al., 2019). This observation supports the notion that specific co-adaptations have occurred between *T. gondii* strains and different feline species (Jewell et al., 1972; Miller et al., 1972; Khan et al., 2014a).

To date, population genetic studies have only partially deciphered the phylogenetic relationship between strains from different geographical areas. In particular, the phylogenetic positioning of the most common clonal lineages relative to other *T. gondii* lineages and populations remains unclear (Su et al., 2012; Lorenzi et al., 2016). In addition, conflicting scenarios have been proposed to explain the global spread of these clonal lineages (Minot et al., 2012; Bertranpetit et al., 2017; Shwab et al., 2018). Given the crucial importance of domestic cats and rodents in the transmission of *T. gondii*, a presumed role of these host species in the global spread of the major clonal lineages has been repeatedly evoked in the literature (Lehmann et al., 2006; Shwab et al., 2018; Galal et al., 2019; Hamidović et al., 2021). However, this hypothesis could not be formally tested previously (Shwab et al., 2018; Hamidović et al., 2021) owing to the paucity of *T. gondii* samples in many regions (Su et al., 2003; Lehmann et al., 2006; Khan et al., 2007; Minot et al., 2012; Lorenzi et al., 2016). Moreover, lack of good estimates of parasite mutation rate and generation time hampered attempts to date dispersal time in relation to expansion history of principal hosts.

To address these questions, we generated the largest dataset of *T. gondii* genomes produced to date (n = 156). We decipher the recent evolutionary history of *T. gondii*, using both global and local ancestry analysis approaches. We provide the first direct estimate of both *T. gondii* mutation rate and generation time that allowed dating major events shaping parasite genome evolution. We uncover candidate genes whose geographic distributions, genomic patterns and stage-specific patterns of expression are most parsimoniously explained by local adaptation to the domestic ecotype and to transmission by domestic cats.

## RESULTS

Paired-end sequence reads from 59 publicly available haploid genomes and from 105 new haploid genomes were aligned to the new PacBio reference assembly RH-88. These samples included isolates from domestic (n=107) and wild animals (n=17), in addition to a number of human isolates (n=42) (Supplementary Table 1). These samples originated from 22 countries (in addition to three French Overseas Departments), and covered most of the global distribution of this species (Fig. 1). New identifiers were assigned to strains included in this study, indicating the ecotype of the strain (Dc for domestic and Wd for wild) and the country of origin. Note that most Old World samples included in this study belonged to clonal lineages previously identified and defined from MS markers (types I,II, III, Africa 1 and Africa 4; Supplementary Table 2), reflecting the very limited genetic diversity of *T. gondii* in the Old World. Many of these samples originated from port regions in Africa (Goree island, Saint-Louis, Dakar, Cotonou, Ouidah, Libreville) and Europe (Bordeaux, Le Havre), the most likely source populations for recent human-mediated global expansion of *T. gondii* strains. Isolates from the Americas originated from both coastal areas (e.g. Sao Paulo, Rio de Janeiro, Cayenne) and inland areas, including wild environments in South and North Americas. In addition, several isolates came from the Caribbean islands (Martinique, Guadeloupe), which are well-known for their importance in maritime history linking Old and New Worlds.

**Fig. 1.**
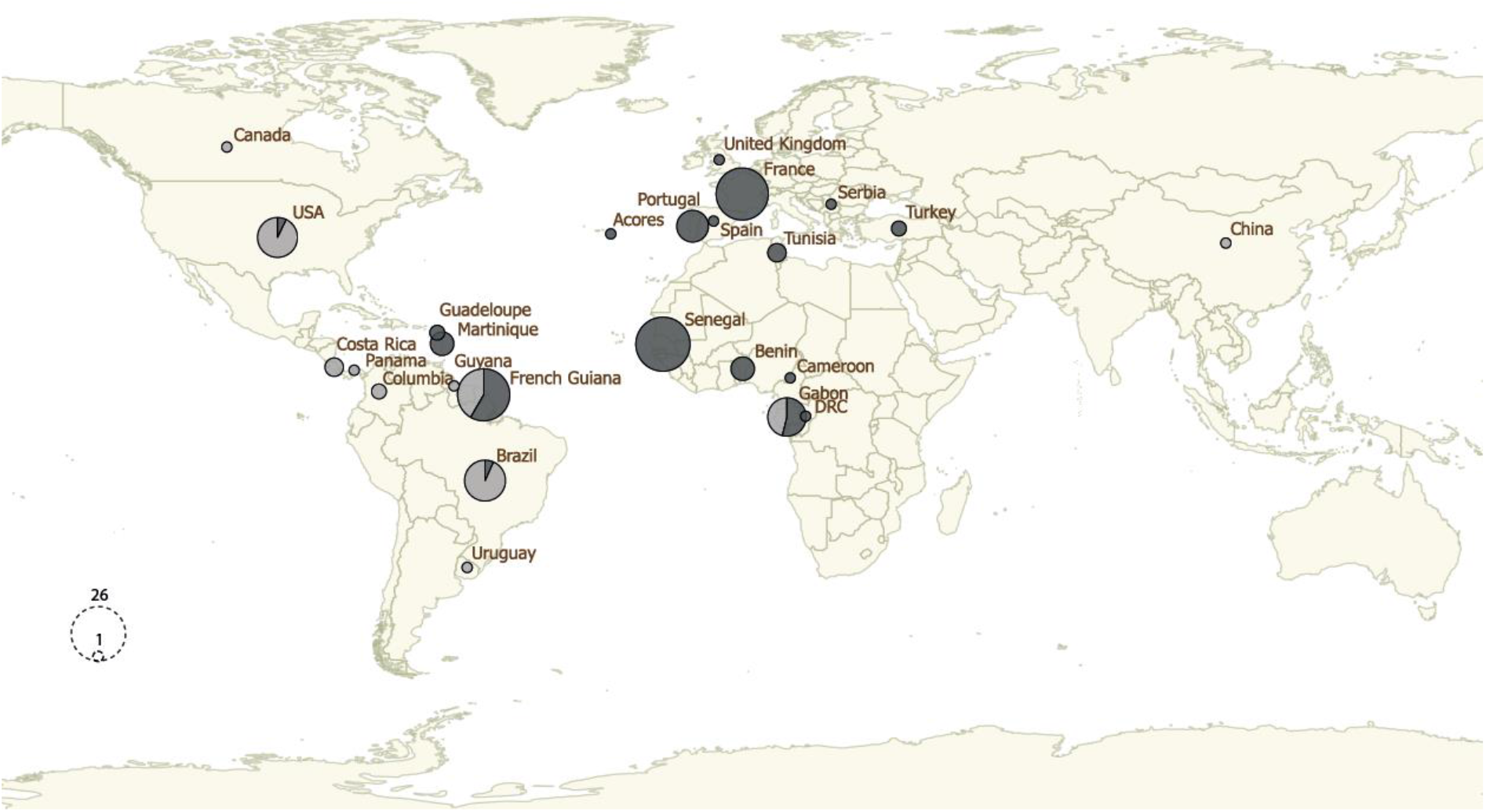
The geographical distribution of *Toxoplasma gondii* strains analysed in this study. Sizes of pie charts correlate with total number of mono-strains isolates for each country. Isolates sequenced specifically for this study are represented in dark grey and whole-genome sequence data from previous studies —publicly available on the European Nucleotide Archive (https://www.ebi.ac.uk/ena/browser/home)— are represented in light grey.

The 105 new genomes from this study were sequenced at a mean depth of 21X, ranging between 8 and 57X (Supplementary Table 1). In total, 156 genomes and 1,790,555 single-nucleotide polymorphisms (SNPs) passed all filtration criteria (see Methods). The 1,790,555 SNP-dataset was used for dating purposes. A second 1,262,582 SNP-dataset was generated after removing singletons SNP from the first one and was used for population genetics analyses.

### Clonal lineages

We first wondered whether the clonal lineages identified in *T. gondii* based on the analyses of multilocus markers was evident at the genome scale. Our analyses (see supplementary note for details) revealed the presence of four intercontinental clonal lineages, as well as a number of regional clonal lineages (restricted to one continent) (Fig. 2a). By matching these results to previous findings relying on the analysis of MS markers, we found that these four intercontinental clonal lineages correspond to type I (found in Asia, Europe, North and South America), type II (cosmopolitan), type III (cosmopolitan) and Africa 1 (found in Africa, South America and Western Asia) (Shwab et al., 2014; Chaichan et al., 2017; Galal et al., 2018). Africa 4 is found in both Africa and Asia, although samples of this lineage from Asia were not available for our study.

**Fig. 2.**
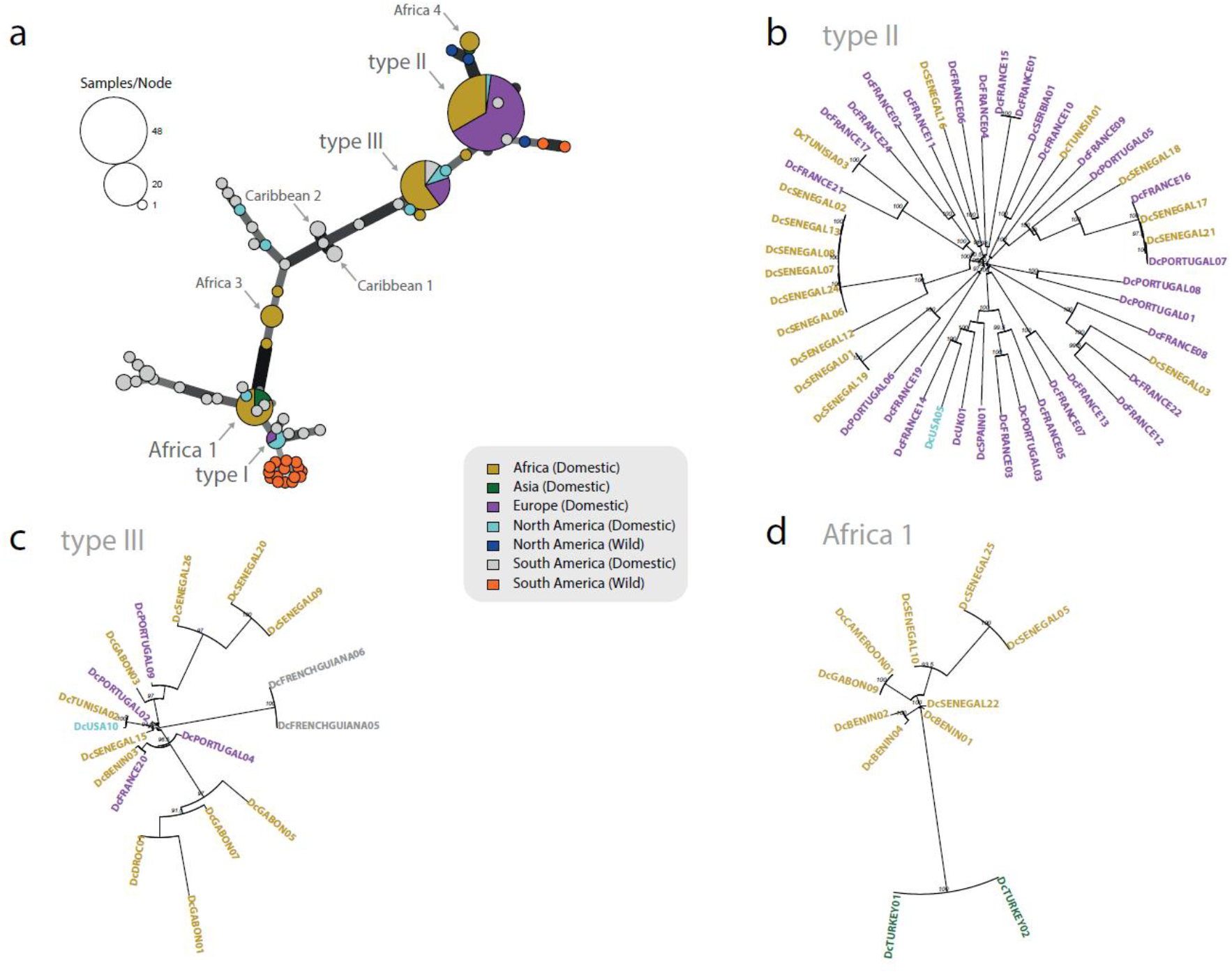
Toxoplasma gondii clonal lineages description. (a) Minimum spanning network of *T. gondii* genomes. Genomes separated by a genetic distance less than or equal to 0.01 are collapsed in a single circle and are considered belonging to the same clonal lineage. The size of each circle corresponds to the number of individuals, and the colours indicate the continent of origin and the ecotype of each individual. Thick and dark lines show MLGs that are more closely related to each other whereas edge length is arbitrary. Neighbour-joining trees of *T. gondii* major lineages (b) type II, (c) type III and (d) Africa 1. Individual genomes are colour-coded according to their continent of origin. Support values greater than 90% using 1,000 bootstrap samples are shown.

Within each clonal lineage, we did not find marked patterns of geographical structure separating strains from the Old and New Worlds (Fig. 2 b-d). Conversly, strong clustering between strains from different continents was repeatedly observed, suggesting recent waves of intercontinental dissemination of these lineages. Four South American clonal lineages could be identified although these lineages were undersampled. Two corresponded to the previously described Caribbean 1 and Caribbean 2 lineages. Most domestic and wild strains from South America were non-clonal.

### Global and local ancestry analyses

To decipher the genetic relationships between the *T. gondii* genomes from different geographical origins, we first performed ancestry analysis using unsupervised clustering with ADMIXTURE. This analysis was carried out after a step of clone-censoring of the dataset (keeping only one representative strain of each clonal lineage / type), resulting in a dataset of 71 strains and 588,777 SNPs. The optimal number of ancestral groups was determined to be five (lowest cross-validation error), but we also examined different K values. Old World strains were mostly a mixture of different intercontinental clonal lineages (types I, II, III, Africa 1 and Africa 4) (Supplementary Fig. 1). In the New World, the Wild Amazonian group (in orange) constituted a well-defined non-admixed ancestral group, clearly divergent from the different intercontinental clonal lineages at different K solutions. In addition, a wild group of strains from the Amazonian forest in South America and of a wild strain from North America could be distinguished at K=9, and was designated as Pan-American. Other New World lineages and strains were composed of New World-specific ancestries (in burgundy and in orange) and intercontinental lineages ancestries, with many of these strains exhibiting a mixed pattern of these two categories of ancestry in the same genome. This latter pattern was mainly noticed among domestic strains, and could be suggestive of hybridization events between intercontinental lineages and New World specific clades.

To better understand the patterns of admixture, we generated a co-ancestry matrix with whole nuclear genome data and independent co-ancestry matrices for each of the 13 nuclear chromosomes using ChromoPainter. This revealed that domestic strains from the New World all shared chunks of chromosomes with at least one of four intercontinental domestic clonal lineages type I, type II, type III and Africa 1 (Supplementary Fig. 2 a-n) and for many of them the wild American strains (Pan-American or Amazonian). By contrast, the four intercontinental domestic clonal lineages did not share any chromosome regions with the two above mentioned wild American populations. Accordingly, we performed local ancestry analyses by defining the intercontinental lineages (type I, type II, type III, Africa 1 and Africa 4) and the unadmixed wild populations (Amazonian and Pan-American) as putative ancestral populations. Putative hybrids (all the other genomes) were made up of few large blocks of different ancestries (up to five ancestors for the same strain), and whole chromosomes of single ancestry are also often observed (Fig. 3). At least one large segment (> 1Mb) of ancestry corresponding to an intercontinental lineage was found in all domestic strains from both Old and New Worlds. One exception was DcURUGUAY01 (CASTELLS) that had almost only an Amazonian ancestry. Amazonian ancestry was identified in nearly all domestic strains from South and Central America, and to a lesser extent in North America, but was absent from almost all Old World domestic strains. Pan-American ancestry was much rarer among putative hybrids, although it was found in both South and North America, including the wild putative hybrid strains from North America. Analyses of the apicoplast genome (a maternally inherited organelle that does not undergo recombination) confirmed the hybrid origin of the American domestic strains, as each apicoplast genome harboured a single ancestry, related either to an intercontinental lineage or to a wild New World population (mainly Amazonian) (Supplementary Fig. 4). In the Old Word, most admixed strains were the result of an admixture between the major lineages, but these hybrids remain rare compared to the American continent where almost all domestic strains have a hybrid origin.

**Fig. 3.**
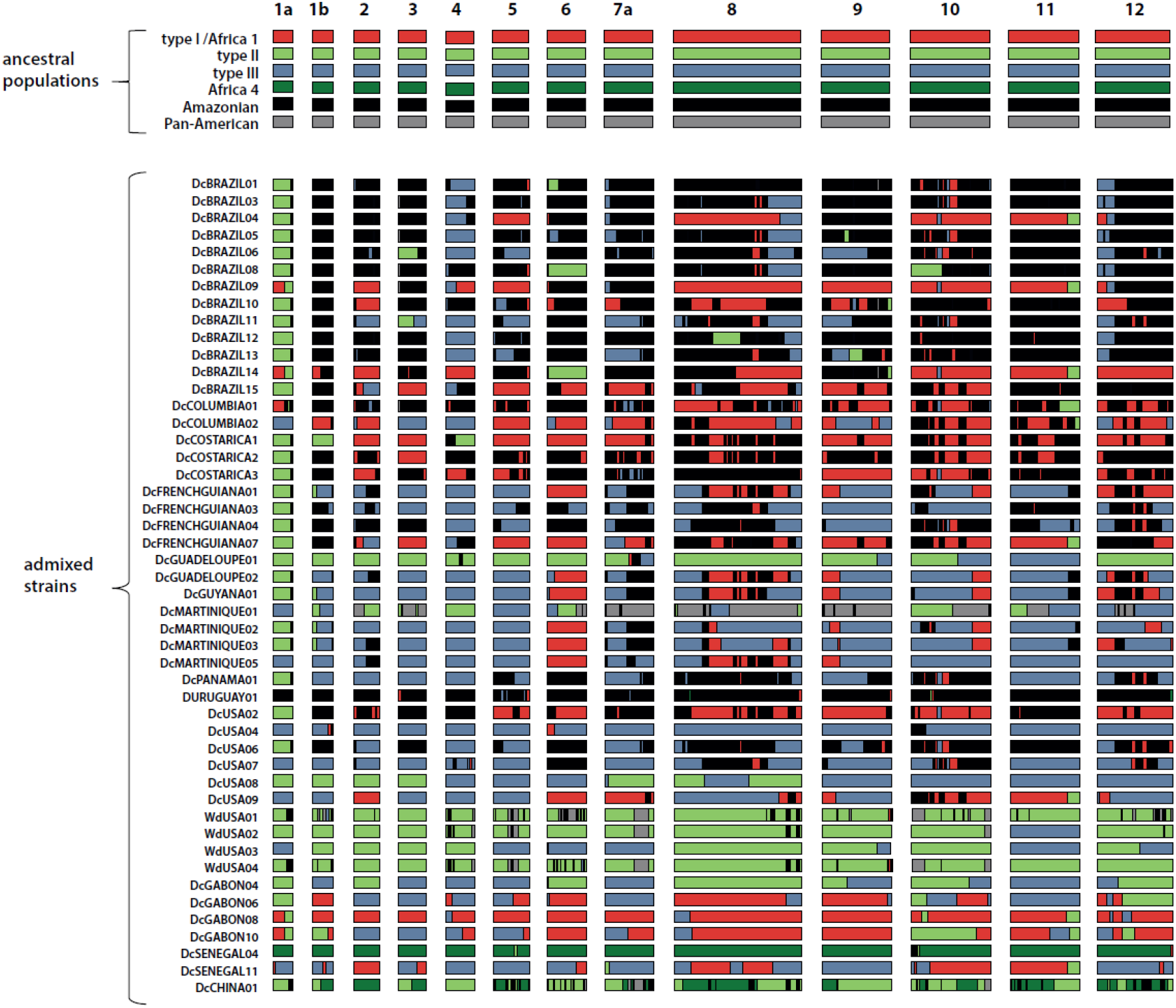
Genome-wide distribution of ancestry in all putative hybrid *Toxoplasma gondii* genomes. Plots are graphically displayed using karyoploteR (Gel and Serra, 2017) and show ancestry estimates at each genomic position for the 13 nuclear chromosomes. Colours reflect putative ancestral populations.

Overall, our results demonstrate that Old and New World *T. gondii* parasites present radically different patterns of genetic diversity. In the Old World (Europe, Africa and Asia), most strains belong to one of the previously defined intercontinental clonal lineages with rare hybrids observed between the different lineages. On the other hand, most New world parasites isolated from wild animals form well-defined non-clonal genetic clusters and those isolated from the domestic environment are the result of hybridizations between intercontinental lineages and a number of New World-specific wild populations (Supplementary Fig. 3).

### Timing of emergence, spread and introgressions of intercontinental *T. gondii* clonal lineages

Using stored aliquots of two long-term *in-vivo* cultured *T. gondii*, we estimated the mutation rate of the parasite (see Supplementary Methods). This was complemented with data from the literature to estimate the parasite generation time. For each clonal lineage using these estimates (ranging from 3.1×10^−9^ to 11.7×10^−9^ mutations per site per year) we first determined the time to the most recent common ancestor (TMRCA) between the two most divergent genomes (Fig. 4a). Regarding the four major intercontinental clonal lineages, our estimates showed that type II emerged 12,980-48,988 years ago, whereas type I, type III and Africa 1 emerged much later: 651-2457, 683-2,578 and 629-2,376 years ago, respectively. Note that inclusion of additional isolates from the same or different geographic areas could alter these estimates by revealing more ancient divergence times between strains of each respective lineage. This is particularly true for type I for which only three samples were available. Regarding the four South American-specific clonal lineages, the divergence times estimated between the most distant isolates ranged between 197 -742 and 225 - 850 years ago, for Caribbean 1 and Caribbean 2, respectively.

**Fig. 4.**
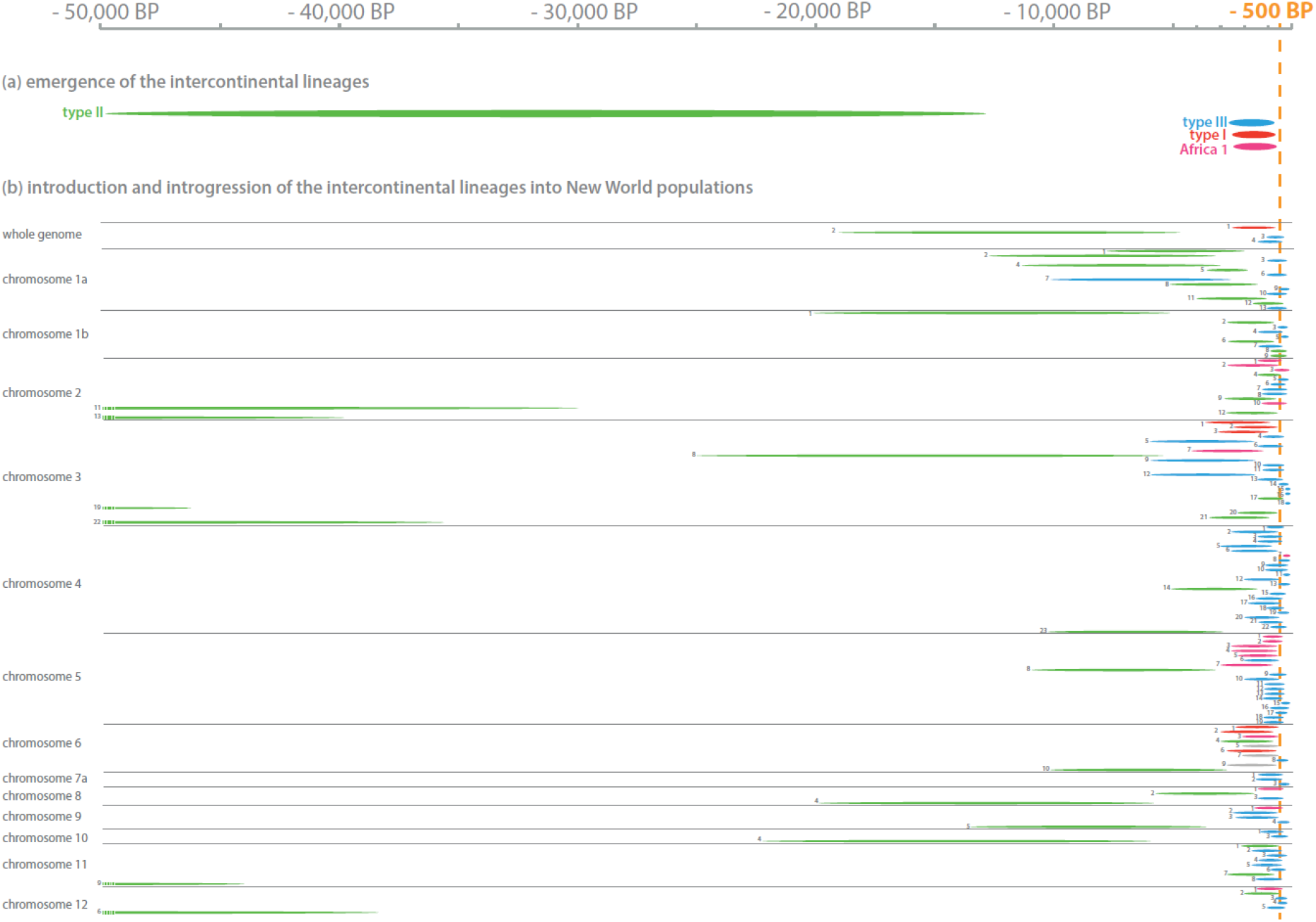
Dating estimates for emergence of intercontinental lineages and their expansion to the New World. (a) The length of coloured ellipses represents the estimated interval for the time to the most recent common ancestor (TMRCA) between the two most divergent genomes of each intercontinental lineage: type I, type II, type III and Africa 1. (b) The length of coloured ellipses represents the estimated interval for the TMRCA between full chromosomes of New World strains and their closest relative from the Old World having the same respective chromosomal ancestry: type I in red, type II in green, type III in blue, Africa 1 in purple and Africa 3 in grey. Precise TMRCA estimates are presented in Supplementary Table 3 and 4. A numeral is attributed to each TMRCA estimate on the figure to facilitate the correspondence with the Supplementary Table 3 and 4.

We next explored the divergence time between New and Old World strains sharing the same ancestry. We calculated TMRCA between New World strains and their closest relative from the Old World belonging to the same intercontinental clonal lineage (Fig. 4; Supplementary Table 3). In addition, we calculated TMRCA between full chromosomes of New World hybrid strains inherited from one of the four intercontinental lineages and their closest relative among Old World strains having the same respective chromosomal ancestry (Fig. 4; Supplementary Table 4). Note that TMRCA estimates are also very sensitive to sampling and obtaining accurate estimates depends on robust sampling of source populations (populations that expanded to the New World). Among isolates of each respective clonal lineage from Africa and Europe, isolates not belonging to source populations are expected to exhibit some degree of divergence from these source populations, and consequently from individuals that have expanded to the New World from these source populations. This divergence bias is expected to be more substantial in more diversified lineages (type II *versus* other intercontinental lineages) owing to greater divergences between strains of the same lineage, even within a single region (also refer to Fig. 2b). In this sense optimal sampling would have included isolates from all port areas historically involved in transatlantic and colonial trade in Europe and Africa, which is only partially the case in this study.

Most estimates of TMRCA indicate that massive introgressions of types I, II, III and Africa 1 into New World populations have occurred in the last few centuries, and provide evidence for very recent migrations of strains between Old and New Worlds (Fig. 4b). A number of TMRCA estimates (in particular among strains having type II ancestry) indicated older divergence times, which could be explained by older migratory waves, or by a divergence bias related to sampling.

Conversely, the wild strains of type 12 (WdUSA01 and WdUSA04) and type II, although having chromosomes of the same ancestry, had TMRCA estimates consistent with a divergence prior to the emergence of type II lineage (Fig. 4; Supplementary Table 4). It confirms that type 12 is a true wild population, sharing no recent ancestry with the domestic type II lineage. WdUSA02 (B41) showed an admixed pattern of divergence from the type II lineage, both recent and more ancient, consistent with a recombination between a domestic type II and a wild type 12 strain. WdUSA03 (B73), although isolated in the wild (from a bear) had a chromosomal ancestry consistent with recent recombination having occurred between type II and type III lineages. This latter observation is consistent with evidence from multilocus markers of a dissemination of domestic strains into the wild ecotype in North America (Jiang et al., 2018).

### Candidate genes contributing to adaptation to the domestic environment

We asked whether hybrid domestic strains from the New World are the result of random recombinations between intercontinental domestic lineages and New World-specific —wild— clades, or whether certain domestic alleles inherited from intercontinental lineages had been selected during this process. This could explain the greater success of hybrid strains compared to wild strains in occupying domestic niches in New World countries. To do so, the 71 genomes of the clone-censored dataset were split into three groups based on results from global and local ancestry analyses: (1) domestic populations (intercontinental lineages and hybrid strains having a recent shared ancestry with these lineages), (2) wild populations (Amazonian, Pan-American and type 12), and (3) DcURUGUAY01 (CASTELLS). This latter strain was used as outgroup given its New World-specific ancestry (Fig. 3), and its divergence from other populations (Lorenzi et al., 2016). The wild strains WdUSA02 and WdUSA03 were included in the domestic group given their recently inherited large genomic segments from domestic lineages of types II and III (Fig. 3 and 4; Supplementary Table 4). We computed the population branch statistic (PBS) using PBScan in order to detect genomic regions of unexpectedly high divergence between the three groups. A unique outlier region of marked divergence between input groups was identified (Fig. 5a); it occurred on chromosome 1a over a region ~600 kb long (Fig. 5b). In parallel, we scanned the nuclear genome for “domestic” variants (alleles common to domestic strains and not found in wild strains). We chose to include DcURUGUAY01 in the wild group given its nearly exclusive Amazonian ancestry (Fig. 3).

**Fig. 5.**
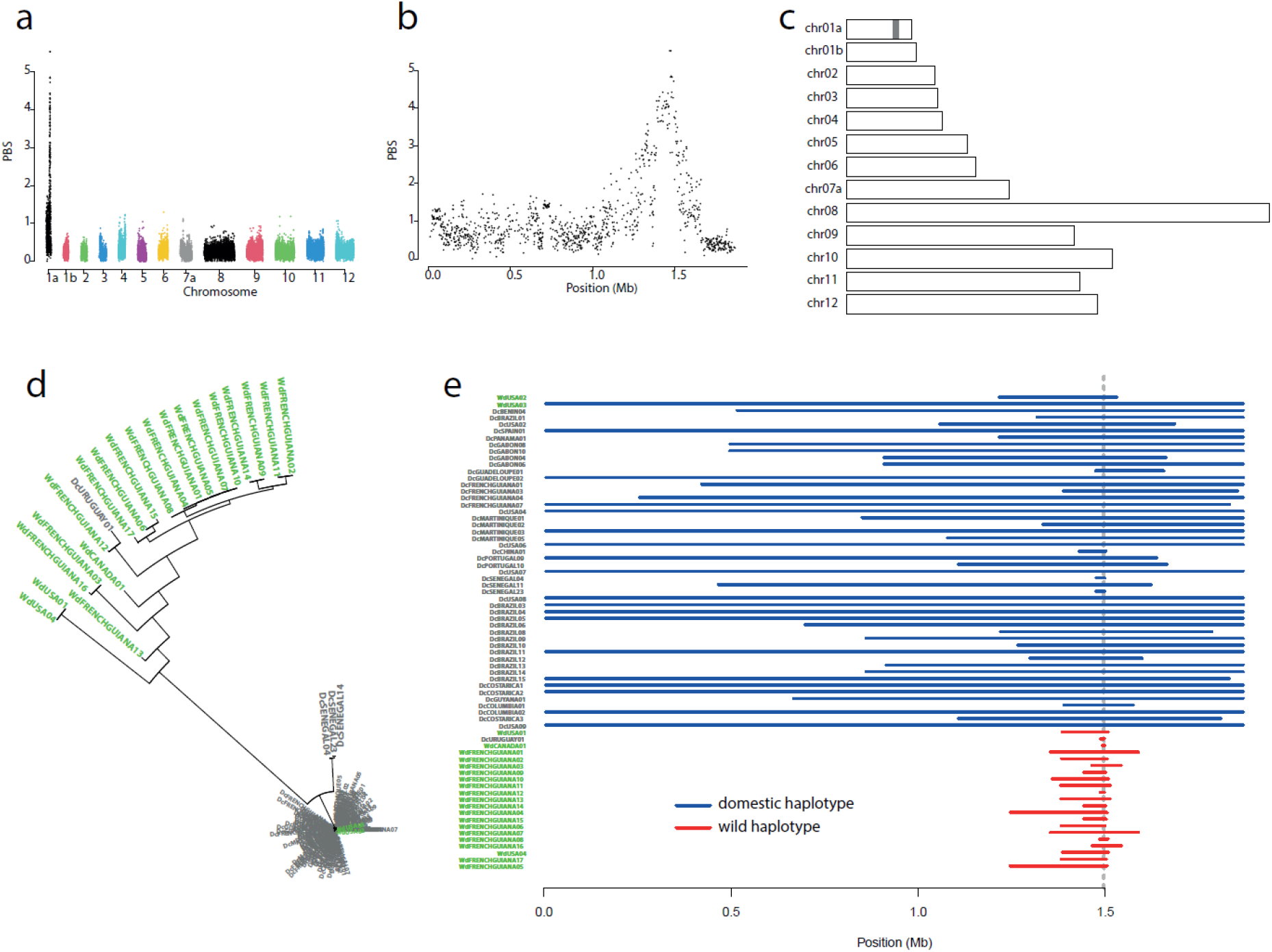
Outlier selection region for adaptation of *Toxoplasma gondii* to the domestic ecotype. (a) The Manhattan plot shows the genome-wide distribution of population branch statistic (PBS) calculated between wild and domestic strains for each 50-SNP sliding window using the clone-censored dataset (n=71). (b) This is a zoom on PBS obtained for chromosome 1a. (c) Genomic positions of domestic variants (n=310) from the clone-censored dataset are shown in grey. (d) Neighbour-joining tree of the outlier selection region (positions 1,390,579 to 1,502,589 on chromosome 1a). It was produced using ape R package by computing genetic distances based on the dissimilarity matrix produced by poppr R package. It includes all the strains of the dataset (n=156). Individual strains are colour-coded according to ecotype of origin (domestic strains in grey and wild strains in green). (e) Haplotype length around the position 1495970 on chromosome 1a in domestic strains relative to wild strains of the clone-censored dataset. The plot shows the boundaries of the longest shared haplotype (the range over which it is identical to at least one other haplotype) around the domestic allele of the focal marker (chr01a_1495970) relative to the wild allele.

In total, 310 variants were identified, of which 58 were missense variants (Supplementary Table 5). Strikingly, all 310 variants occurred within the outlier region identified using PBS statistic, over a region ~150 kb long containing 25 genes (Fig. 5c), between positions 1,352,873 and 1,501,765 (Supplementary Table 6). When we consider the ancestry pattern of this region (Fig. 3), we notice that nearly all hybrids strains have a type II/type III ancestry between positions 1,390,579 and 1,502,589. This latter genomic region of ~100 kb corresponds to a remarkably conserved haplotype shared by all domestic strains and clearly divergent from wild haplotypes (Fig. 5d). Interestingly, only Africa 4 strains showed some degree of divergence from other domestic strains, although they carried all 310 domestic variants identified within the outlier region of selection. Extended linkage disequilibrium was observed around this outlier region in domestic strains relative to wild ones, a signal of positive selection acting specifically on domestic strains (Fig. 5e).

In order to determine which gene(s) is/are most likely associated to adaptation to the domestic ecotype, we analysed the function of the 25 candidate genes present in the ~150 kb genomic region (carrying the 310 domestic variants) and their stage-specific pattern of expression. Although 14 genes were annotated for protein function, data was lacking in the literature about possible associations between allelic heterogeneity and differential adaptations to specific hosts or ecotypes. By focusing on stage-specific patterns of expression, we found that six genes displayed stage-specific expression as they were expressed at only one stage (Ramakrishnan et al., 2019; Farhat et al., 2020). Among them two genes were found to be only expressed during enteroepithelial stages (EES) of development (early cellular forms characteristic of the onset of the sexual stage in cat enterocytes): TGRH88_020260 (TGME49_295995) and TGRH88_020330 (TGME49_295920). Missense variants segregating domestic from wild strains were only found for TGRH88_020330 (TGME49_295920). This gene is annotated as encoding a hypothetical protein. Five missense variants segregating domestic strains from wild strains were found for this gene (Supplementary Tables 5 & 7).

## DISCUSSION

### Intercontinental lineages spread from the Old to the New World

The global dataset of whole genome sequences studied herein confirm previous findings based on multilocus genotyping (mainly MS and RFLP) of a strong clonal structure for most *T. gondii* populations (Su et al., 2012; Galal et al., 2019). We provide strong evidence of intercontinental dissemination of the most prevalent lineages (types I, II, III and Africa 1) using a strictly clonal mode of propagation. All these four lineages are well adapted to transmission by the domestic cat as they are frequently isolated in the domestic environment (Chaichan et al., 2017; Galal et al., 2018; Shwab et al., 2018).

We estimated that *T. gondii* lineage type II emerged 12,980-48,988 years ago (Fig. 6). At this period, cat domestication was still an ongoing process, and it is therefore likely that type II lineage was circulating in wild cats (*Felis silvestris*). The domestication of cats first occurred in the Near East, coincident with agricultural village development in the Fertile Crescent about 10,000 years ago (Driscoll et al., 2007; Ottoni et al., 2017). This *T. gondii* lineage could have accompanied cats of subspecies *F. s. lybica* (one of the five known clades of wild cats of species *Felis silvestris*, found in different regions of the Old World) that established themselves in human settlements in the Near East, or emerged later in the domestic environment from other wild cat populations in Africa, Asia or Europe following the expansion of the Neolithic revolution to those regions. Type I, type III and Africa 1 lineages emerged much later, 651-2457 years, 683-2,578 years and 629-2,376 years ago, respectively. Importantly, these domestic lineages emerged before dissemination of domestic cats to the New World 500 years ago; it is hence likely that they emerged in the Old World. From the beginning of the Iron age, about 3,000 years ago, maritime transportation substantially increased in the Old World (Jones et al., 2013). Maritime activities of Romans in Antiquity (Peters, 1998), of Phoenician during medieval times (Bonhomme et al., 2011) and of Vikings between the 7^th^ and the 11^th^ centuries (Jones et al., 2012), contributed to the expansion of domestic cats, mice and rats throughout the Mediterranean basin and to Continental Europe. During these periods, Egyptian domestic cats gradually took over other populations of domestic cats such as in Anatolia before spreading to most areas of the Old World (Ottoni et al., 2017). These movements of *T. gondii* hosts could have promoted encounters with previously allopatric populations of *T. gondii* (e.g. between Europe and Africa), fostering the emergence of new lineages by hybridization. This hypothesis is supported by the shared ancestry that is observed on certain genomic regions between the most common lineages. Boyle et al. (2006) showed that types I and III are respectively second- and first-generation offspring of a cross between a type II strain and one of two unknown ancestral strains. The most parsimonious scenario is that these lineages emerged following the expansion of the geographical range of type II strains in the Old World during this period.

**Fig. 6.**
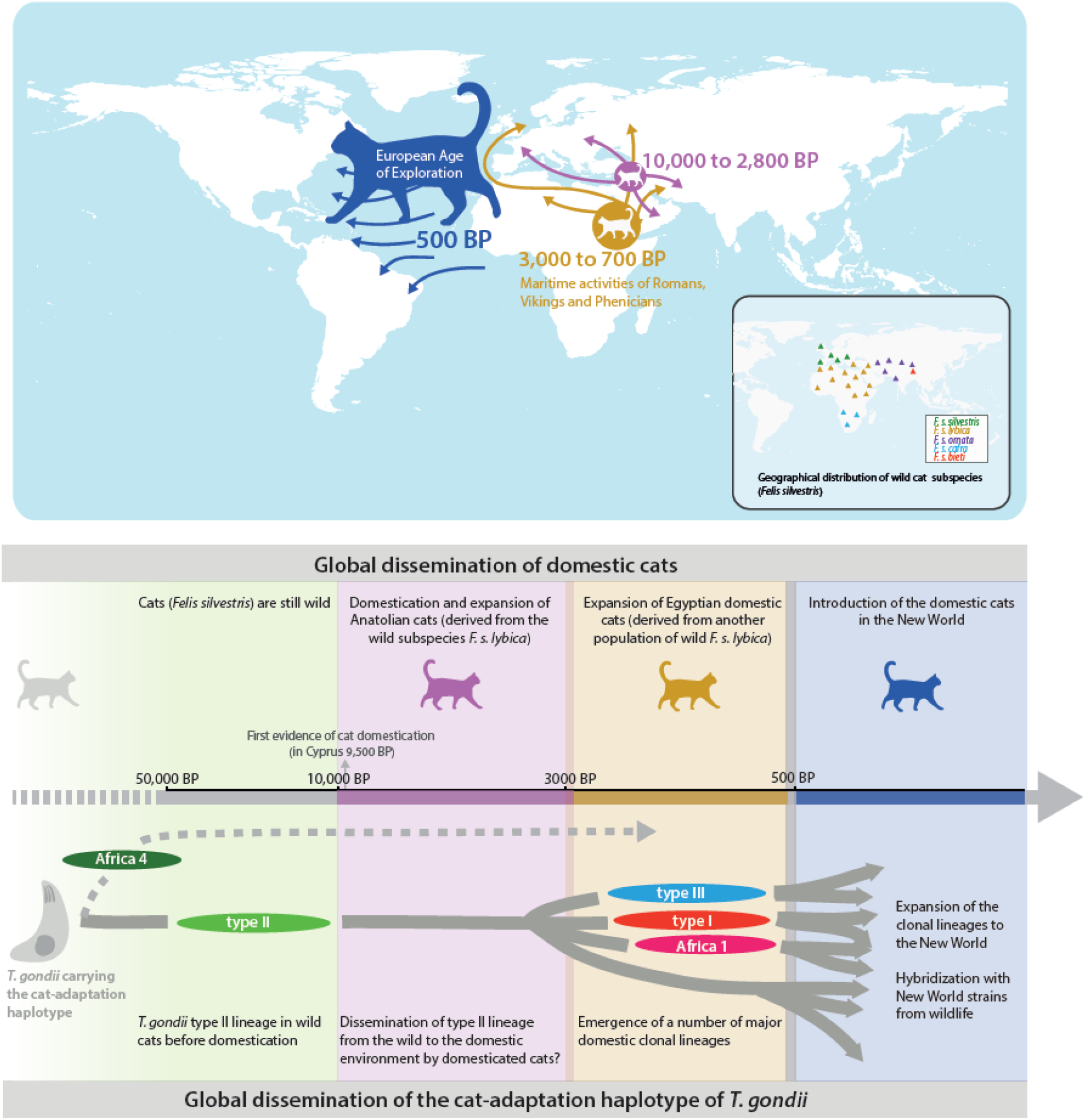
Graphical summary of major events that have punctuated the recent evolutionary history of *Toxoplasma gondii* in relation to history of cats’ dispersal. Geographical distribution of wild cat subspecies and historical data about cats’ dispersal are derived from Ottoni et al. (2017). The grey dotted arrow in the bottom part of the figure indicates that emergence of Africa 4 lineage could not be dated in this study due to a lack of samples belonging to this lineage.

### Introgression of the Old World domestic lineages into New World populations

A previous study showed that common inheritance of large haploblocks is the major factor in determining the phylogenetic grouping of *T. gondii* strains (Lorenzi et al. 2016).

Here, we demonstrate that nearly all New World domestic strains are hybrid strains harbouring large chromosomic regions of either types I, II, III or Africa 1 ancestry. Moreover, many TMRCA estimates suggest that hybridization events that gave rise to these strains occurred after the introduction of these lineages in the Americas in the last six centuries. These estimates coincide to the onset of the European “age of exploration” (Subrahmanyam and Alam, 2007). During this period, human activities enabled domestic cats, but also mice (*Mus musculus)* and rats (*Rattus rattus* and *Rattus norvegicus*) to reach the Americas for the first time (Lipinski et al., 2008; Macholán et al., 2012; Puckett et al., 2016), allowing for the first time the emergence of a domestic cycle of *T. gondii* in those areas.

It is noteworthy that hybrid strains are much more common in the New World (especially in South America) compared to the Old World indicating —currently or in the past— more frequent recombination events. Most hybrid strains appear to be the results of only one or a few rounds of meiotic reproduction when considering the chromosomal pattern of ancestry of experimental hybrids (Khan et al., 2014b). Indeed, we did not observe a fine mosaic of different ancestries alternating across genomes, as is usually observed when sexual recombination often occurs in a population (Henn et al., 2012; Fitak et al., 2018; Kim et al., 2020). Sexual recombination in *T. gondii* is favoured by mixed infections in cat prey, which is limited by the immunity developed by an intermediate host following its primary infection (referred to in the Introduction). This immunity often protects the intermediate host from new infections with different strains, but not from highly divergent strains as found in South America (Elbez-Rubinstein et al., 2009). We propose that following their introduction in the Americas, rodents infected with Old World lineages were exposed to highly divergent strains from the wild environment near human settlements. Their immunity being unable to contain these new infections, the rodents could have become superinfected with highly divergent strains giving rise to tissue-cysts of New World strains alongside tissue-cysts emanating from their primary infection. This unique situation would provide favourable conditions for the emergence of hybrid populations due to cats feeding on these superinfected intermediate hosts. The emergence of big cities, the decline of wildlife and the great proliferation of domestic cats would have gradually limited the exposure of domestic intermediate hosts to wild strains in South and North America.

We show that for certain chromosomes type II has sister clades (close but distinct) among wild strains from North America. The time to the most recent common ancestor (TMRCA) estimated for domestic type II and wild type 12 (WdUSA01 and WdUSA04) on several chromosomes is clearly anterior to the onset of domestication and to the emergence of type II. Evidence from apicoplast sequences also shows that Asian Chinese 1 shares a common ancestor with wild type 12. Note that all these strains belong to the same clade (refer to Supplementary Fig. 1). These data support the occurrence of *T. gondii* migrations between Asia and North America, probably anterior to the Neolithic revolution and the domestication era. Migrations were probably mediated by movements of animal herds through the land bridge formed by the Bering Strait during the late Pleistocene period until about 13,000 years ago. During this period a corridor was created by falling sea levels that provided an opportunity for Asian species including mammoths, bison, muskoxen, caribou, lions, brown bears, and wolves to move into North America (Guthrie, 2004; Lowe and Walker, 2014; Froese et al., 2017; Phillips et al., 2018). Assuming a role of late Pleistocene animal species in disseminating *T. gondii*, an Asian origin of this clade appears more likely given the direction of migrations inferred for these animals. It is consistent with the hypothesis of an Old World origin of type II lineage, as previously suggested by Shwab et al. (2018) using multilocus markers and not a North American origin as proposed by other studies (Khan et al., 2007; Minot et al., 2012).

### A candidate gene for adaptation of *T. gondii* to the domestic cat

We sought to identify genes under positive selection for adaptation of *T. gondii* strains to the domestic ecotype. Our genome-wide scan for selection identified a unique genomic region of ~100 kb (0.16% of the whole *T. gondii* genome) on chromosome 1a exhibiting a nearly perfect dichotomy between wild strains and domestic strains from all over the world. The few experimental infections carried out on domestic cats showed that cats infected with domestic lineages (carrying this domestic haplotype) produce oocysts more efficiently compared to when they are infected with wild strains (Khan et al., 2014b). The signal we found of a unique global haplotype common to domestic strains is therefore probably the results of an adaptation to the domestic cat (*Felis catus*), not forgetting that the latter is the only indispensable host species for the transmission of *T. gondii* in the domestic environment.

Note that a number of strains shared the same haplotype for the whole length of chromosome 1a, a pattern previously noticed in past studies (Khan et al., 2007; Lorenzi et al., 2016). We provide strong evidence that this pattern is specific to domestic strains. It can be explained by the much stronger linkage disequilibrium observed in domestic strains relative to wild ones around the outlier region identified in this study due to selection, often reaching the whole length of chromosome 1a. Given the rarity of sexual recombination in *T. gondii* populations, it is likely that a number of domestic lineages and strains inherited the entire chromosome 1a from an ancestor carrying the advantageous allele. Other domestic strains inherited a more or less important portion of chromosome 1a from this ancestor, leading to the tightening of the inherited portion around the domestic haplotype under selection following successive rounds of sexual recombination.

This domestic haplotype exhibited some degree of genetic divergence between Africa 4 strains and all other domestic strains. This divergence probably occurred in the Old World given the exclusive occurrence of Africa 4 lineage in Africa and Asia. This observation provides additional evidence for an Old World origin of the “cat adaptation haplotype”, not forgetting that it is also where domestic cats first emerged. It is therefore likely that this haplotype spread to the domestic environment from two different sources: the Africa 4 lineage that expanded in Africa and Asia, and the type II lineage (the presumably oldest domestic lineage) at the origin of the global expansion of the haplotype.

Within the outlier region of selection, TGRH88_020330 (TGME49_295920) was selected as the top candidate gene for adaptation to domesticity, as it was the only gene specifically expressed by *T. gondii* during its early stages of sexual reproduction in cat enterocytes that also carried functionally relevant (missense) variants (n = 5) segregating domestic *T. gondii* strains from wild ones. However, it was not possible to determine which of these missense variants has functional relevance. The genomic region carrying this gene exhibits strong linkage disequilibrium among domestic strains implying that certain variants have been fixed by hitch-hiking. The expression of this gene of unknown function begins at the earliest stages of sexual multiplication (Ramakrishnan et al., 2019; Farhat et al., 2020), suggesting that it could have a role in specific host recognition, to enable the initiation of sexual reproduction when a given *T. gondii* strain infects the proper host species within the felidae family. The highly successful haplotype carried by this genomic region and shared by almost all domestic strains probably enables efficient parasite transmission by present-day domestic cats, not forgetting that according to our dating estimates all variants constituting this haplotype were already fixed before the onset of domestication. It suggests that wild strains harbouring this haplotype before domestication must have been efficiently transmitted by wild ancestors of the present-day domestic cats.

Di Genova et al. (2019) have recently produced cat intestinal organoids for experimental purposes, an important breakthrough to study sexual reproduction of *T. gondii* knowing the important ethical concerns associated with the use of live cats. Developing similar experimental models from intestinal cells of wild felids could enable a more accurate understanding of the function of the top candidate gene and other candidate genes occurring within the outlier region of selection.

In summary, we have produced a large dataset of high-quality *T. gondii* genomes and estimated the parasite’s mutation rate and generation time. We dated the emergence of the most common *T. gondii* clonal lineages, their recent dispersal and introgressions into New World populations of *T. gondii.* We show that the substantial diversity of domestic strains found in the New World is the result of hybridizations between four recently introduced Old World domestic lineages (adapted to domestic cats) and New World strains from wildlife. A unique cat-adaptation *T. gondii* haplotype —today carried by almost all domestic strains worldwide— has been largely conserved since its initial emergence from wilderness to domestic settlements, and during its dissemination in the Old World and its recent expansion to the New World. The selection of this now global domestic *T. gondii* haplotype is most parsimoniously explained by its role in the initiation of sexual reproduction of *T. gondii* in domestic cats. Importantly, parasite gene(s) involved in the initiation of sexual reproduction could be promising targets for the development of a cat vaccine. In the context of a One Health integrated vaccine programme (Innes et al., 2019), controlling oocysts excretion by domestic cats is considered crucial, since it could be the most efficient way to break the cycle of transmission, limit environmental contamination by oocysts and prevent infection of other T. gondii hosts including humans.

## METHODS

We studied 106 *T. gondii* isolates provided by the French Biological Resource Centre (BRC) for *Toxoplasma* (http://www.toxocrb.com/). This certified structure (NF S96-900 standard) manages the storage of T. *gondii* strains from human or animal toxoplasmosis to make them available to the scientific community. Our analyses were complemented with whole-genome sequence data of 59 strains (100 bp paired-end reads) from a previous study (Lorenzi et al., 2016) made publicly available on the European Nucleotide Archive (https://www.ebi.ac.uk/ena/browser/home).

Following parasite culture (Supplementary Methods), total genomic DNA was extracted from 200μl of tachyzoites suspension, using the QIAamp DNA MiniKit (Qiagen, Courtaboeuf, France). *Toxoplasma gondii* DNA extracts were genotyped using 15 MS markers in a single multiplex PCR-assay, as described previously (Ajzenberg et al., 2010).This step was necessary to check for cross-contaminations between samples during culture and to identify mixed infections. A mixed infection was identified in one isolate (FR-Mac fas-002; *Macaca fascicularis* ; Mauritius) by the presence of two alleles at 12 loci; only one strain was genotyped at the time of isolation, and it is therefore likely that the second strain initially had a lower tissue load in the infected host, and took longer to grow to levels sufficient for library construction. Only the mono-strain isolates (n=105) were sequenced. DNA was sheared into 400–600-base pair fragments by focused ultrasonication (Covaris Adaptive Focused Acoustics technology, AFA Inc, Woburn, USA). Standard indexed Illumina libraries were prepared using the NEBNext DNA Library Prep kit (New England BioLabs), followed by amplification using KAPA HiFI DNA polymerase (KAPA Biosystems). 150 bp paired-end reads were generated on the Illumina NextSeq 500 according to the manufacturer’s standard sequencing protocol. The 105 samples sequenced for this study are deposited in ENA under the study accession number XXXXXXXXX.

### Mapping and variant calling

Recent advances in variant calling enabled the use of publicly available panels of validated single-nucleotide polymorphisms (SNPs) and indels to estimate the accuracy of each base call and minimize the generation of false positive SNPs. Unfortunately, building these panels is labour-intensive and such data is lacking for under-studied organisms such as *T. gondii*, which still falls under the category of “non-model” organisms. Ribeiro et al. (2015) explored the relationship between the choice of tools and parameters in non-model organisms, their impact on false positive variants, and formulated recommendations for variant calling. Here, we followed their recommendations in order to minimize the call of false positive variants, which was a critical point regarding our objective of estimating the occurrence times of recent events in *T. gondii* evolution. In this sense, reads were submitted to a stringent mapping configuration (not more than 2% of mismatches), by using BWA 0.7 against the newly available PacBio reference genome RH-88 (13 nuclear chromosomes that cover 63.97 Mb; available on https://toxodb.org; release date 2020-05-15). Mapped reads were sorted with Samtools 1.11 and duplicate reads were marked with Picard ‘MarkDuplicates’ 2.25. Individual BAMs were subsequently merged, and variant calling was performed with FreeBayes 1.3.5 that is considered better than the routinely used Genome Analysis Toolkit (GATK) in non-model organisms (Ribeiro et al., 2015; Calarco et al., 2018). At the individual level, alignments having a mapping quality (--min-mapping-quality) less than 20, a coverage (--min-coverage) less than 3, and alleles having a supporting base quality (--min-base-quality) less than 20 were excluded from the analysis. Genotype calls having fraction of conflicting base calls of more than 10% were also excluded. Finally, only individuals having missing genotype data less than 5% were kept for subsequent analyses to minimize false negative calls. At the population-level, SNPs having a high missing genotype rate (>10%) were filtered out. With the above filters in place, two individuals were excluded (missing genotype data >5%), before excluding an additional 6 samples due to unreliable information about the country of origin and/or the ecotype. The filtered set of SNPs was annotated with snpEff 5.0 6 using the RH-88 annotation file.

For the 35kb *T. gondii* apicoplast genome, mapping of reads was performed against the ME49 reference genome assembly (release date 2013-04-23), as the PacBio reference genome RH-88 had a high proportion of low complexity sequence (~70% in RH-88 genome *versus* ~30% in ME49 reference genome; data not shown). The mapping quality of reads in these low complexity regions was too low and therefore these regions were not exploitable for sequence analysis. A mapping configuration and variant calling parameters identical to those used for the nuclear genome were used for the apicoplast genome. In addition to the six samples having unreliable information about the country of origin and/or the ecotype, six additional samples were excluded due to a high frequency of missing genotype data (>5%).

### Clonality

Most computational tools for population genetics are based on concepts developed for sexual model organisms. Microbial pathogens are often clonal or partially clonal, and hence require different tools to address their population dynamics and evolutionary history. The R package *poppr 2.0* (Kamvar et al., 2015) specifically addresses issues with analysis of clonal and partially clonal populations. We first used this package to collapse individuals into clonal groups, by defining a genetic distance threshold based on 3 different clustering algorithms using the function `mlg.filter` (Kamvar et al., 2015). This initial step, besides enabling definition of clonal lineage boundaries is a necessary partial correction for a bias that affects metrics of most computational tools that often rely on allele frequencies assuming panmixia. A dissimilarity matrix was produced by *poppr* to compute genetic distances between genomes and a minimum spanning network was drawn based on these calculations, by collapsing individuals based on the previously defined genetic distance threshold. Neighbour-joining trees were generated for each intercontinental lineage with *ape* R package (Paradis and Schliep, 2019) using the dissimilarity matrix produced by *poppr* R package.

### Global ancestry inference

In order to identify ancestral populations and to characterize the admixture patterns in our dataset, we used ADMIXTURE 1.3 (Alexander and Lange, 2011). ADMIXTURE is useful as an exploratory tool in analyses of genetic structure, but should be interpreted with caution, since such model-based algorithms often provide only a caricature of a complex reality. The dataset was clone-censored (including only one randomly chosen strain from each clonal lineage) and pruned for linkage disequilibrium in PLINK10 (v. 1.07) using parameters --indep-pairwise 50 5 0.2 (it removes each SNP that has an R^2^ value greater than 0.2 with any other SNP within a 50-SNP sliding window, advanced by 5 SNPs each time). ADMIXTURE was run using the unsupervised mode with cross-validation (--cv). The number of ancestral populations (K) varied between 2 and 10.

We complemented our global ancestry analyses with ChromoPainter (Lawson et al., 2012), known to be particularly useful to discern signatures of recent admixture. ChromoPainter estimates the number of “chunks” of ancestry inherited by an individual or a population from a “donor” individual or population, and builds a co-ancestry matrix that summarizes the degree of sharing of ancestry among all pairs of individuals. Unlike ADMIXTURE, ChromoPainter takes into account patterns of linkage disequilibrium allowing one to combine information across successive markers to increase the ability to capture fine-scale population structure. Hence, linkage disequilibrium pruning was not required for this analysis and we used the unpruned clone-censored dataset.

### Local ancestry inference

Local ancestry inference (also designated as ancestry deconvolution) is the task of identifying the regional ancestral origin of chromosomal segments in admixed individuals. It requires specifying a set of candidate non-admixed populations as putative ancestors of the admixed individuals. After defining ancestral and admixed individuals using approaches of global ancestry inference we carried out local ancestry analysis using the recently released Ancestry_HMM software (Corbett-Detig and Nielsen, 2017). Ancestry_HMM is based on a novel hidden Markov model that does not require genotypes from reference panels and that is generalized to arbitrary ploidy, and is hence suitable for non-model haploid organisms.

In order to estimate the divergence between ancestral populations and their putative progeny as defined by Ancestry_HMM, we generated neighbour-joining trees for each of the 13 chromosomes using *ape* R package that computed genetic distances based on the dissimilarity matrix produced by *poppr* R package. In addition, a TCS network was produced from the apicoplast sequences using PopART.

### Dating emergence, global spread, and introgressions of intercontinental *T. gondii* clonal lineages

We estimated the mutation rate of *T. gondii* based on the *in vivo* mutation rate of the RH strain, through successive passage in outbred mice during 30 years (Supplementary Methods). We estimated the mutation rate of *T. gondii* to range between 3.1×10^−9^ to 11.7×10^−9^ mutations per site per year. We used Tang’s equation (Tang et al., 2002) to calculate the time to the most recent common ancestor (TMRCA), which we adapted to haploid genomes : 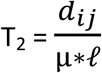, where *d_ij_* is the number of nucleotide differences between any two sequences i and j, *μ* is the mutation rate per site and ℓ is the length of the studied genomic region.

### Candidate genes for adaptation to the domestic environment

To identify candidate genes involved in the process of adaptation to the domestic environment a multistage process was used. We first carried out a divergence-based selection scan using the population branch statistic (PBS) (Yi et al., 2010). This method enables to detect genomic regions of unexpectedly high differentiation between different pre-defined groups (estimated with FST measure by Hudson et al. (1992)), a pattern indicative of directional selection (Lewontin and Krakauer, 1973). In parallel, in order to ascertain candidate SNPs, we used the bcftools 1.1 -- *private* function to identify SNPs that differentiate domestic strains from wild strains (alleles common to domestic strains and not found in wild strains). SNPs identified with these two approaches were annotated with SnpEff in order to define a primary list of candidate genes. To determine which candidates genes are most likely to be under positive selection, we focused on functional evidence. We carefully searched the literature for the function of candidate genes and their stage-related patterns of expression. Selection can result in patterns of extended linkage disequilibrium (LD) and extended haplotype homozygosity (EHH) around the selected site, especially relative to the alternative allele. EHH is defined as the probability that two randomly chosen chromosomes carrying the same allele at a focal SNP are identical by descent over a given distance surrounding it. Hence, for outlier SNPs validated by all types of evidence — environmental, statistical and functional—, we computed EHH using the using the *REHH 2.0* R package (Gautier et al., 2017) to examine the extent of linkage disequilibrium around them.

## Acknowledgements

We thank the French Agence Nationale de la Recherche (ANR project IntroTox 17-CE35-0004) and the Nouvelle-Aquitaine region (Directorate of Research, Higher Education and Technology Transfer), for funding this research. We thank Nicolas Plault, Lionel Forestier and Eden Lebrault for technical assistance. We want to acknowledge the Biological Resource Center for *Toxoplasma* for providing strains and the French National Reference Center for toxoplasmosis for data regarding human strains. We thank Aurélien Dumètre and Emmanuelle Gilot-Fromont for their advice on estimating the generation time of *Toxoplasma gondii.* We would also like to thank Gordon Langsley for revision of the English language. Genome sequencing was performed by the genomic platform GENOM’IC of Cochin Institute in Paris. Animal work was conducted in the animal facility managed by the technical platform BISCEm (US042 Inserm - UMS 2015 CNRS) of the University and CHU of Limoges. The computations presented in this article were carried out on the CALI calculator of the University of Limoges (CAlcul en LImousin), funded by the Limousin region, the European Union, the XLIM, IPAM, GEIST institutes, and the University of Limoges.

