## SUPPLEMENTARY INFORMATION for "A unique *Toxoplasma gondii* haplotype under strong selection has accompanied domestic cats in their global expansion"

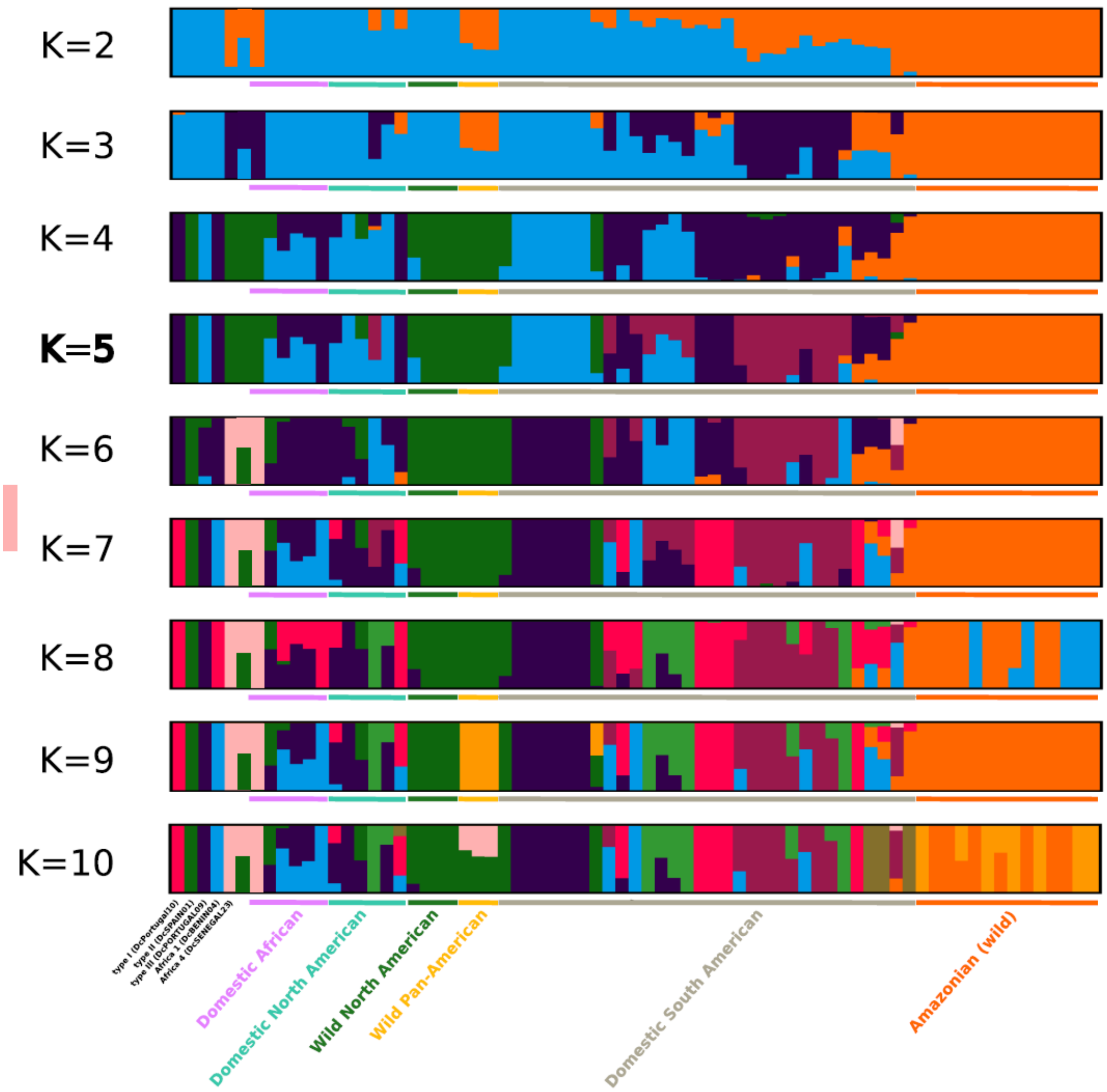

**Supplementary Fig. 1. Ancestry plots showing proportions of ancestral populations for each *Toxoplasma gondii* genome for K = 2 to 10.** Ancestry plots are graphically displayed using CLUMPAK (Kopelman et al., 2015). Apart from samples representing the major clonal lineages (in the left), samples are ordered according to ecotype and continent of origin.

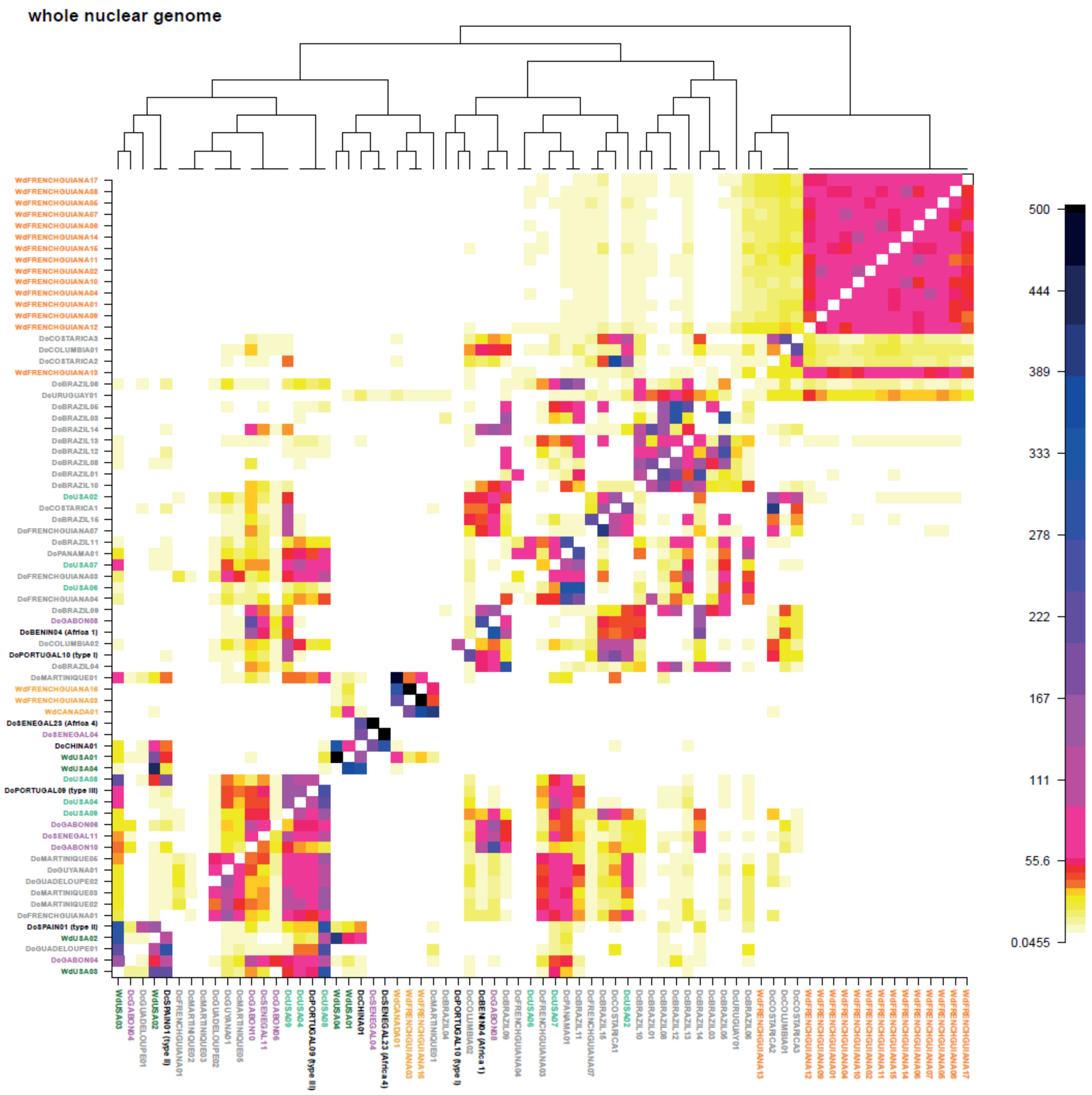

*Supplementary Fig. 2a.*

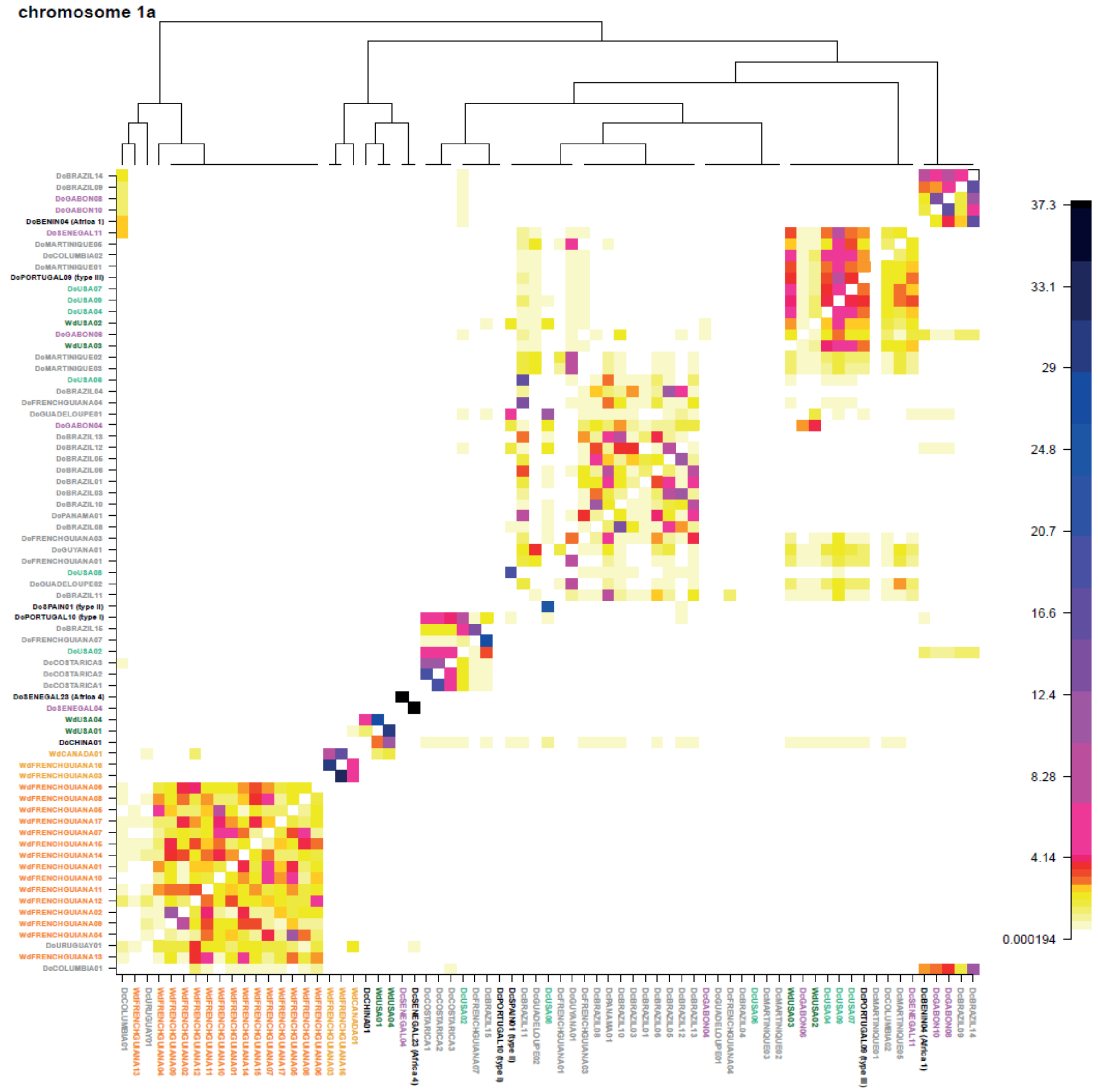

*Supplementary Fig. 2b.*

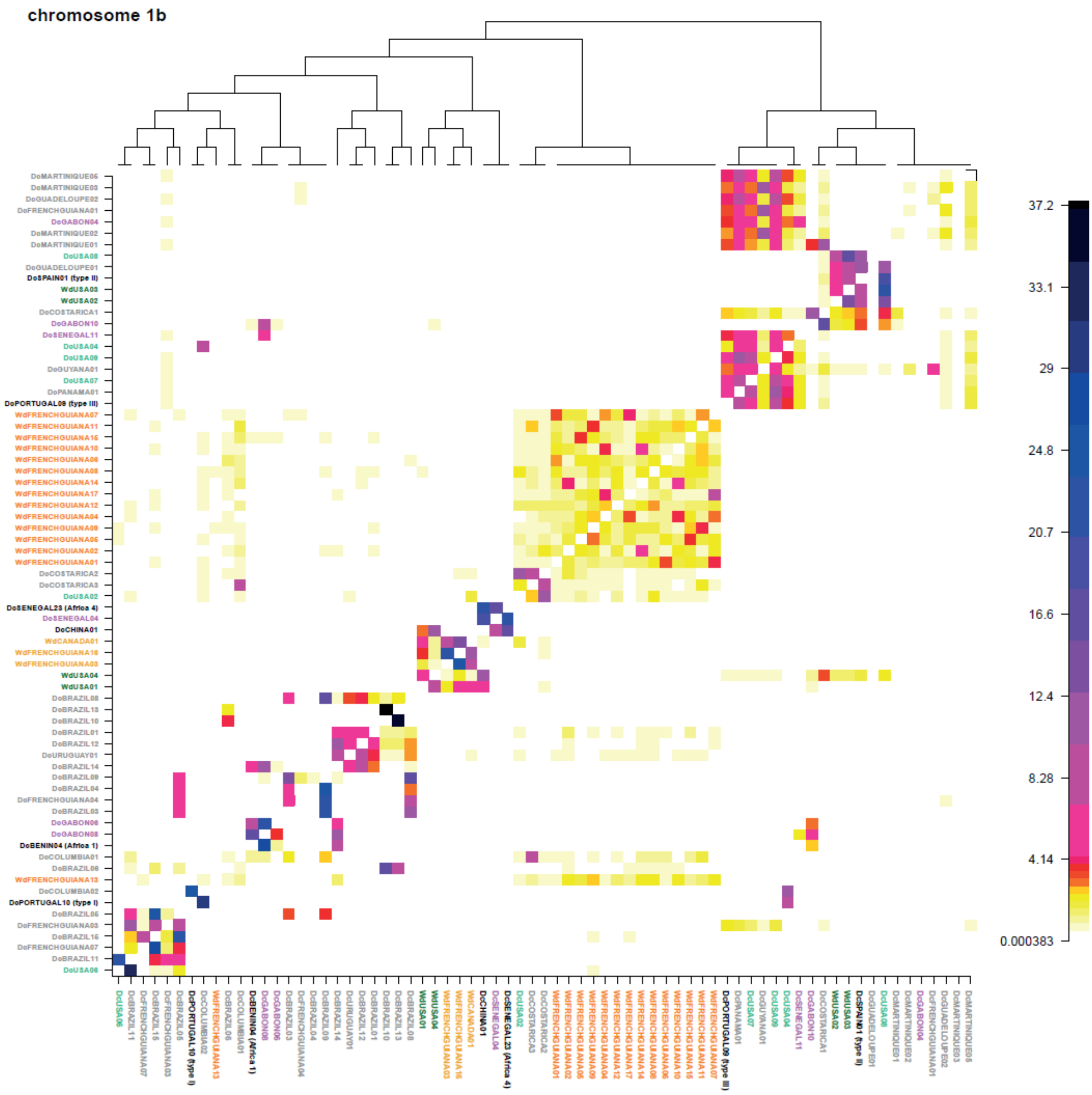

***Supplementary Fig. 2c.***

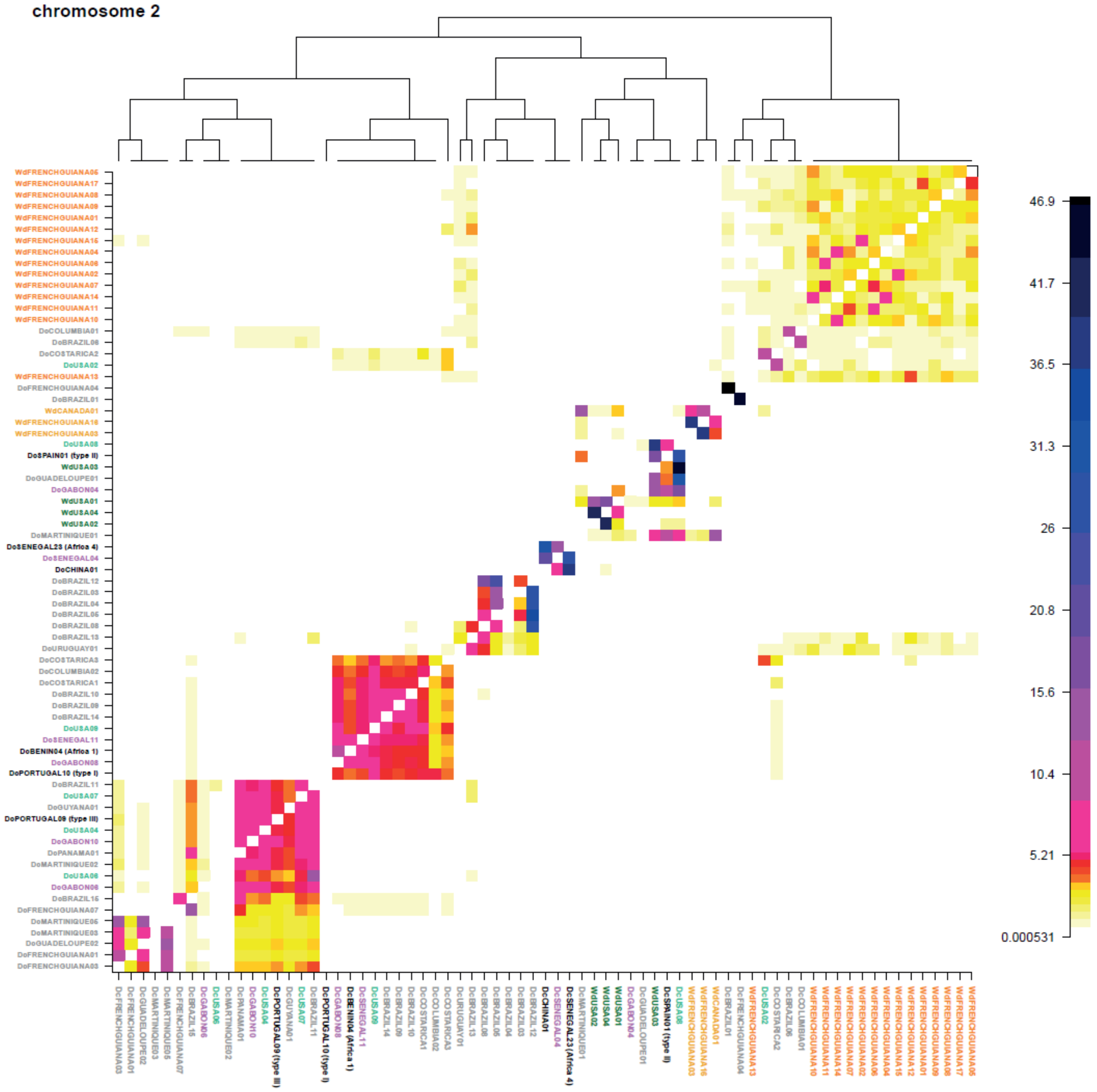

*Supplementary Fig. 2d.*

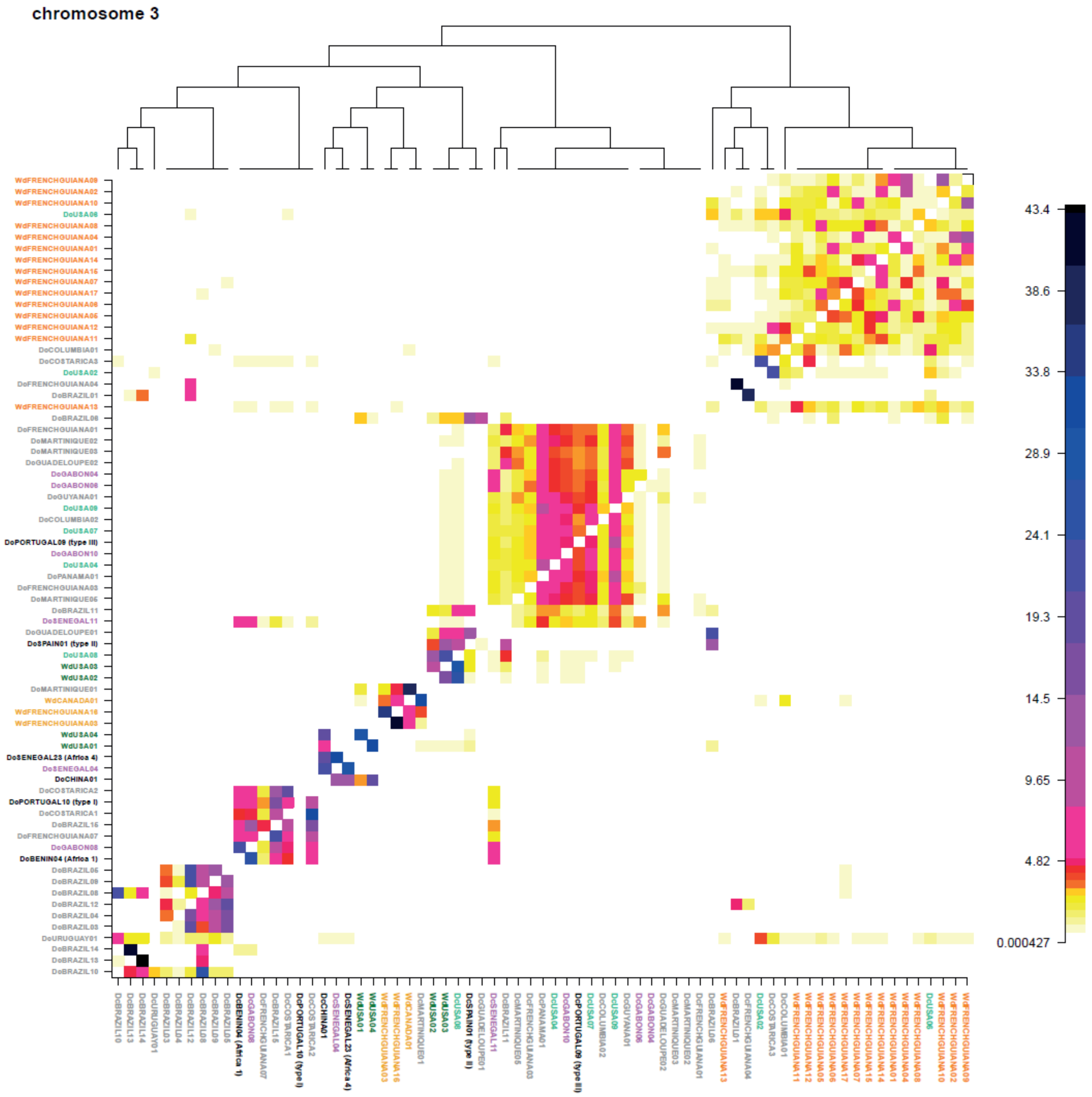

***Supplementary Fig. 2e.***

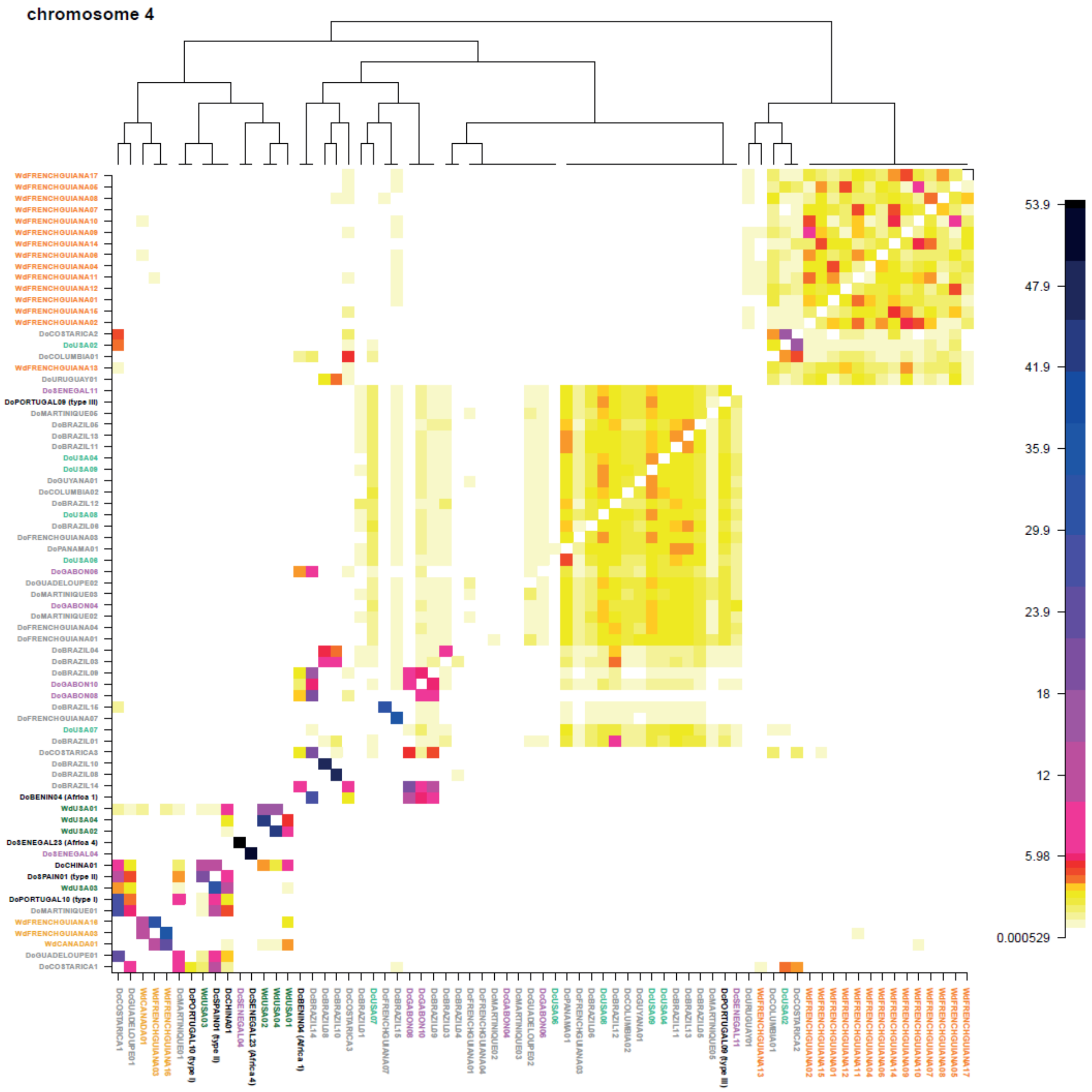

*Supplementary Fig. 2f.*

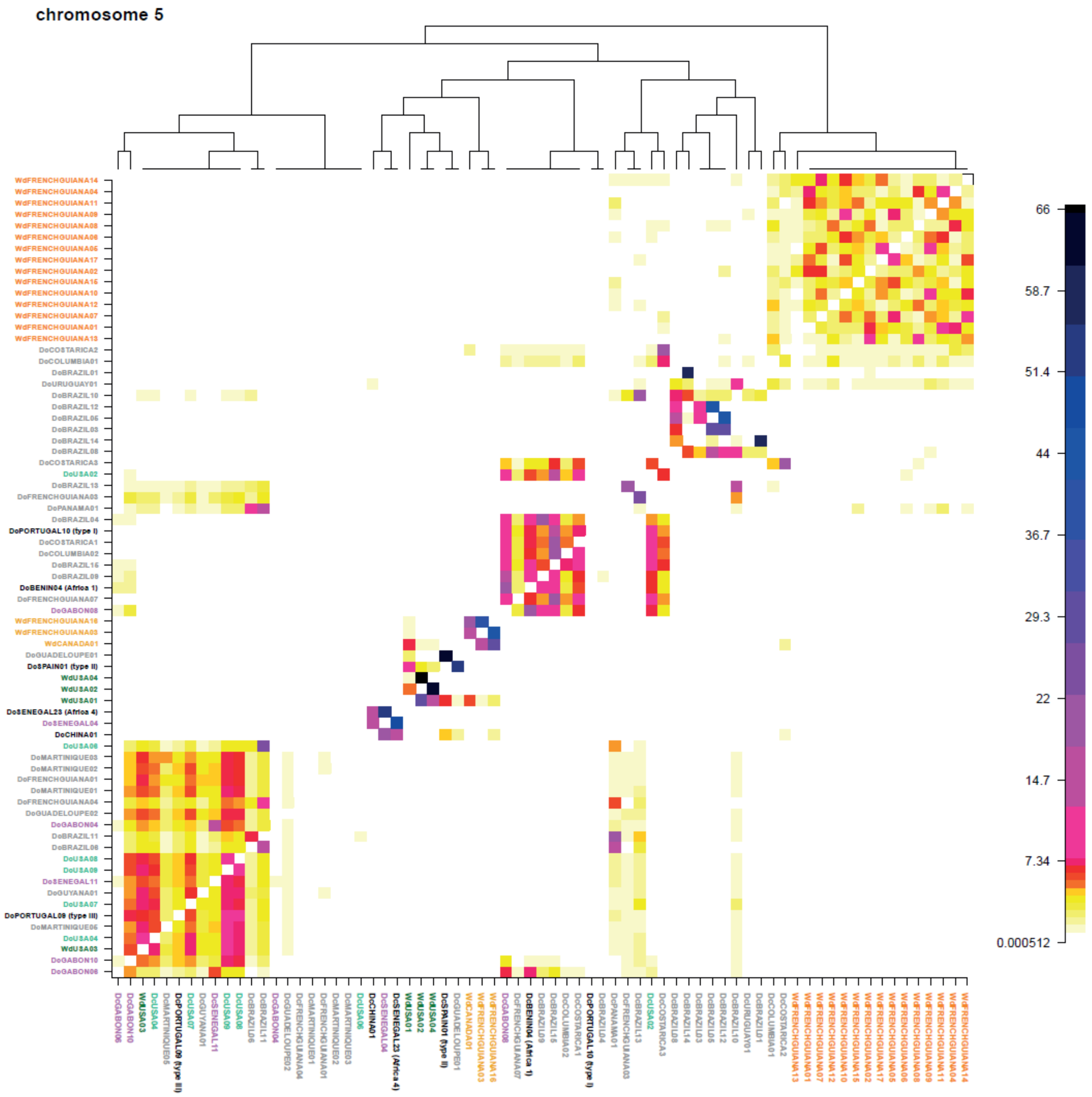

*Supplementary Fig. 2g.*

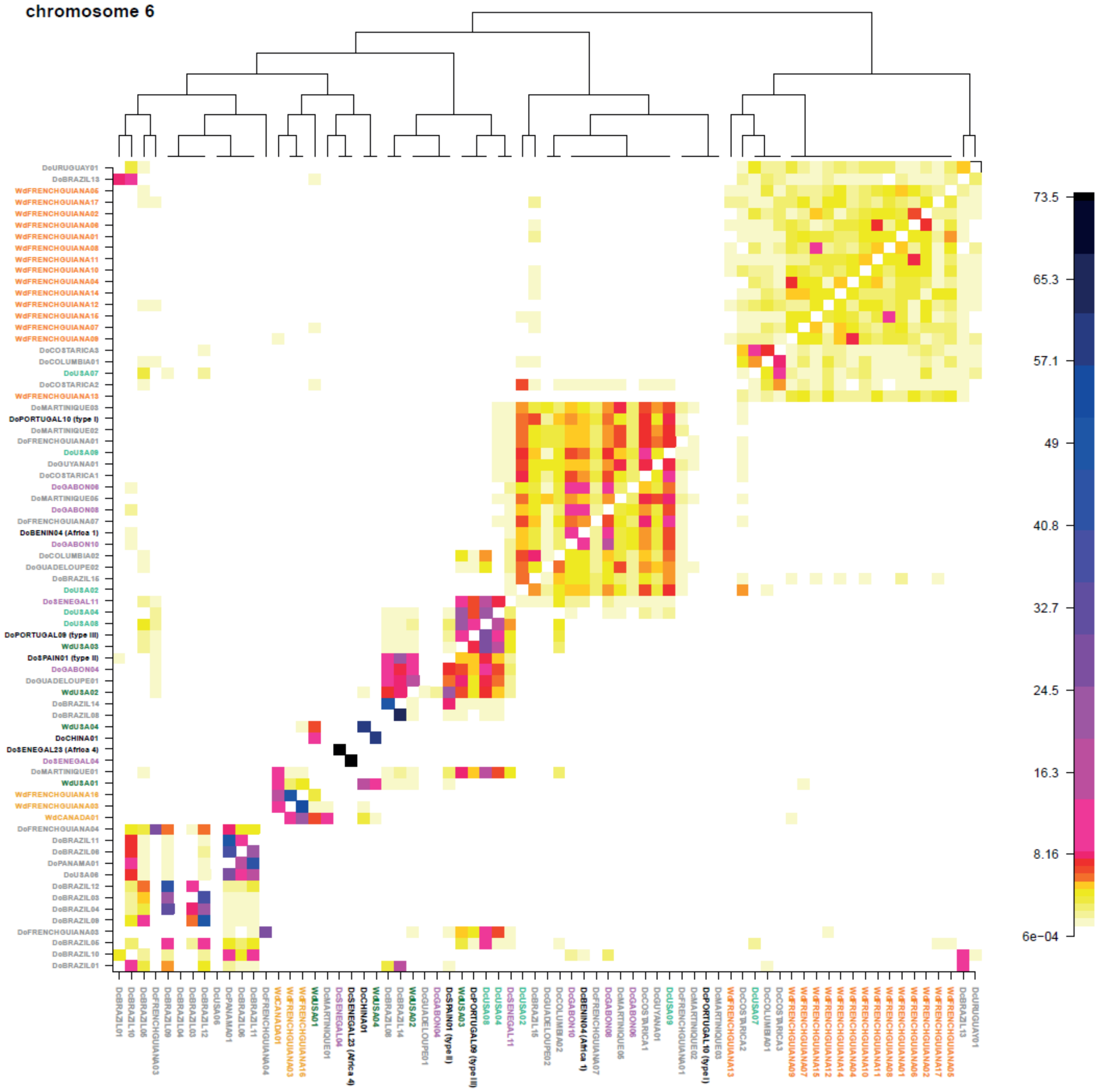

*Supplementary Fig. 2h.*

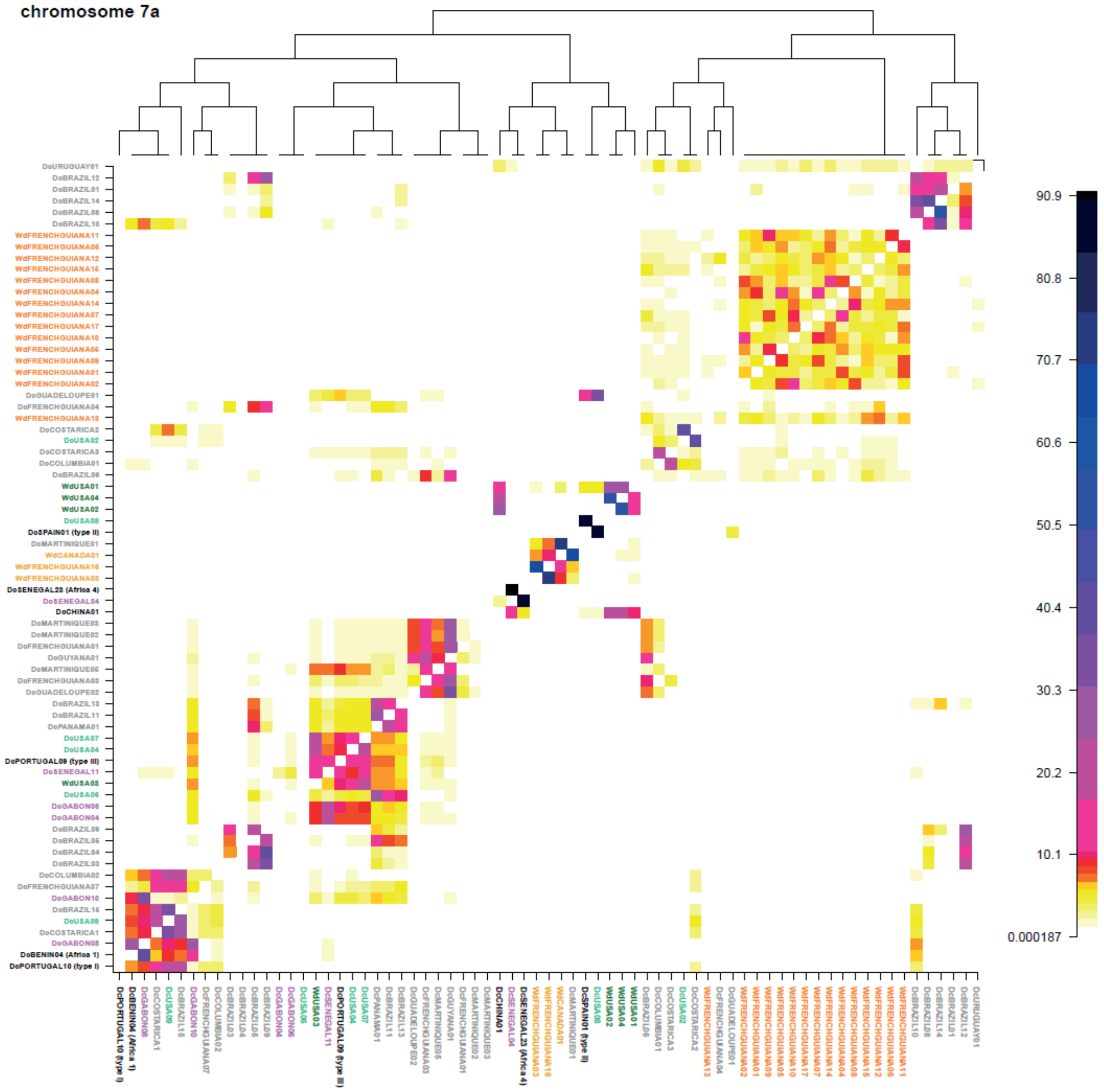

***Supplementary Fig. 2i.***

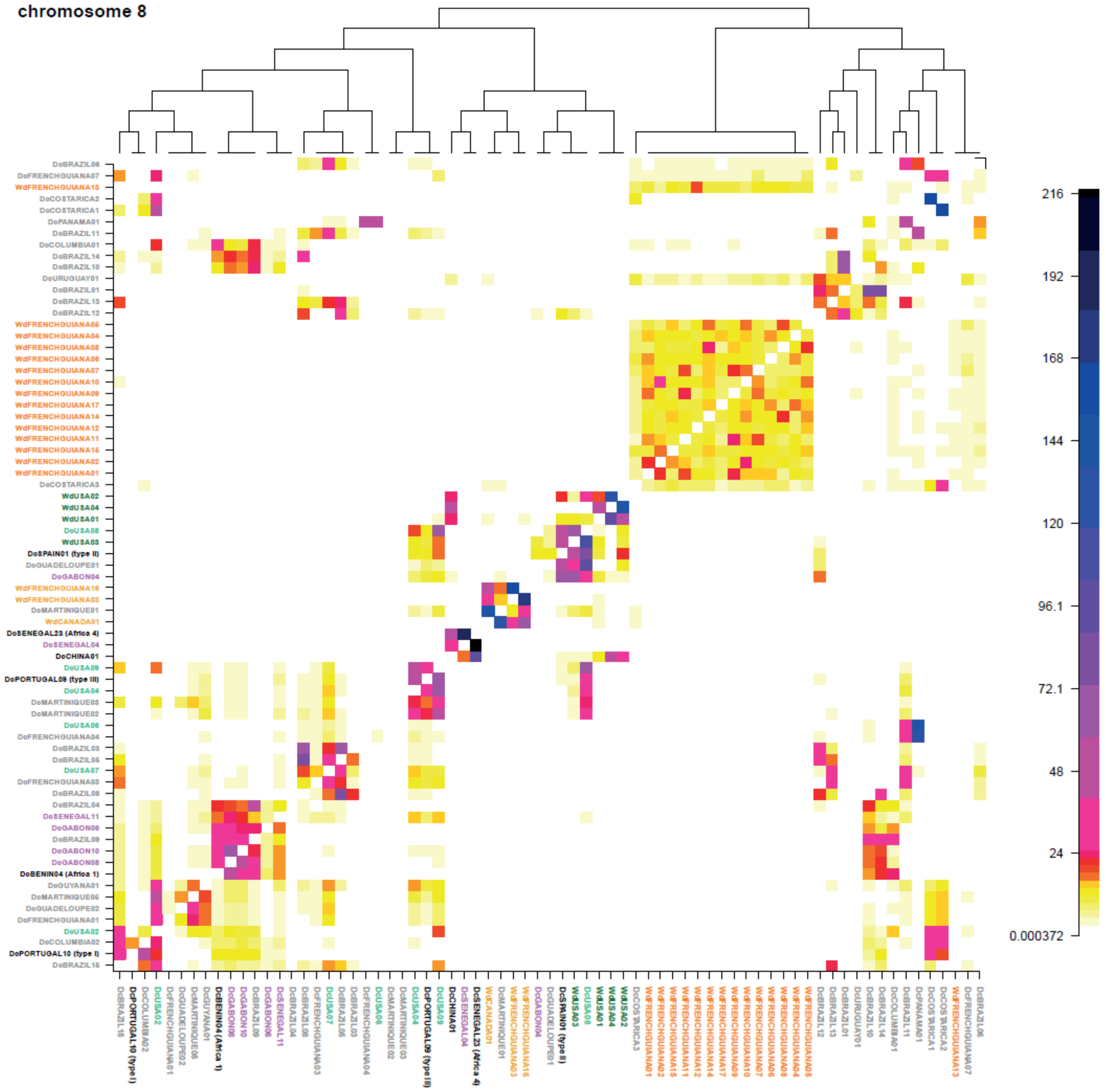

***Supplementary Fig. 2j.***

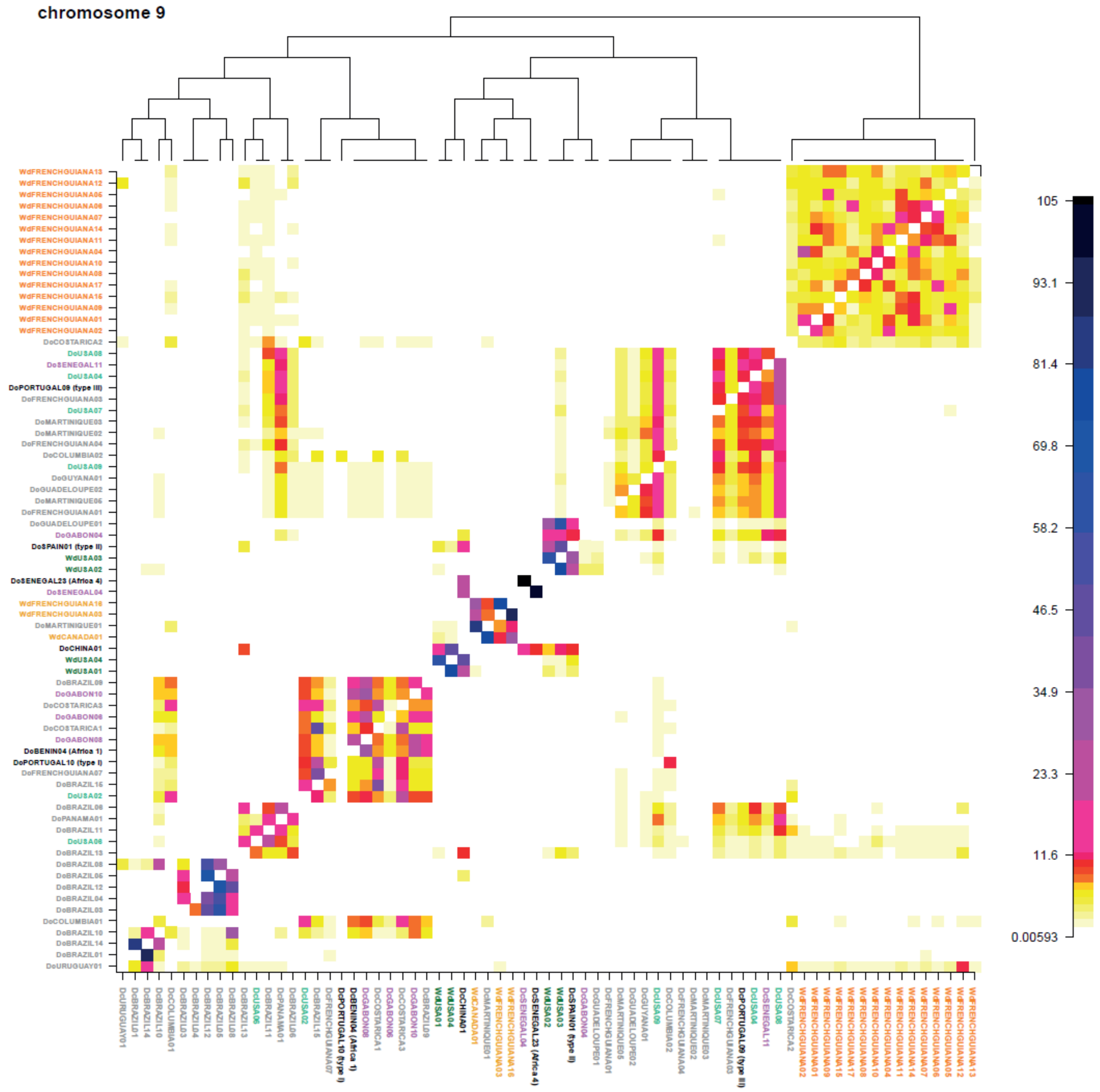

*Supplementary Fig. 2k.*

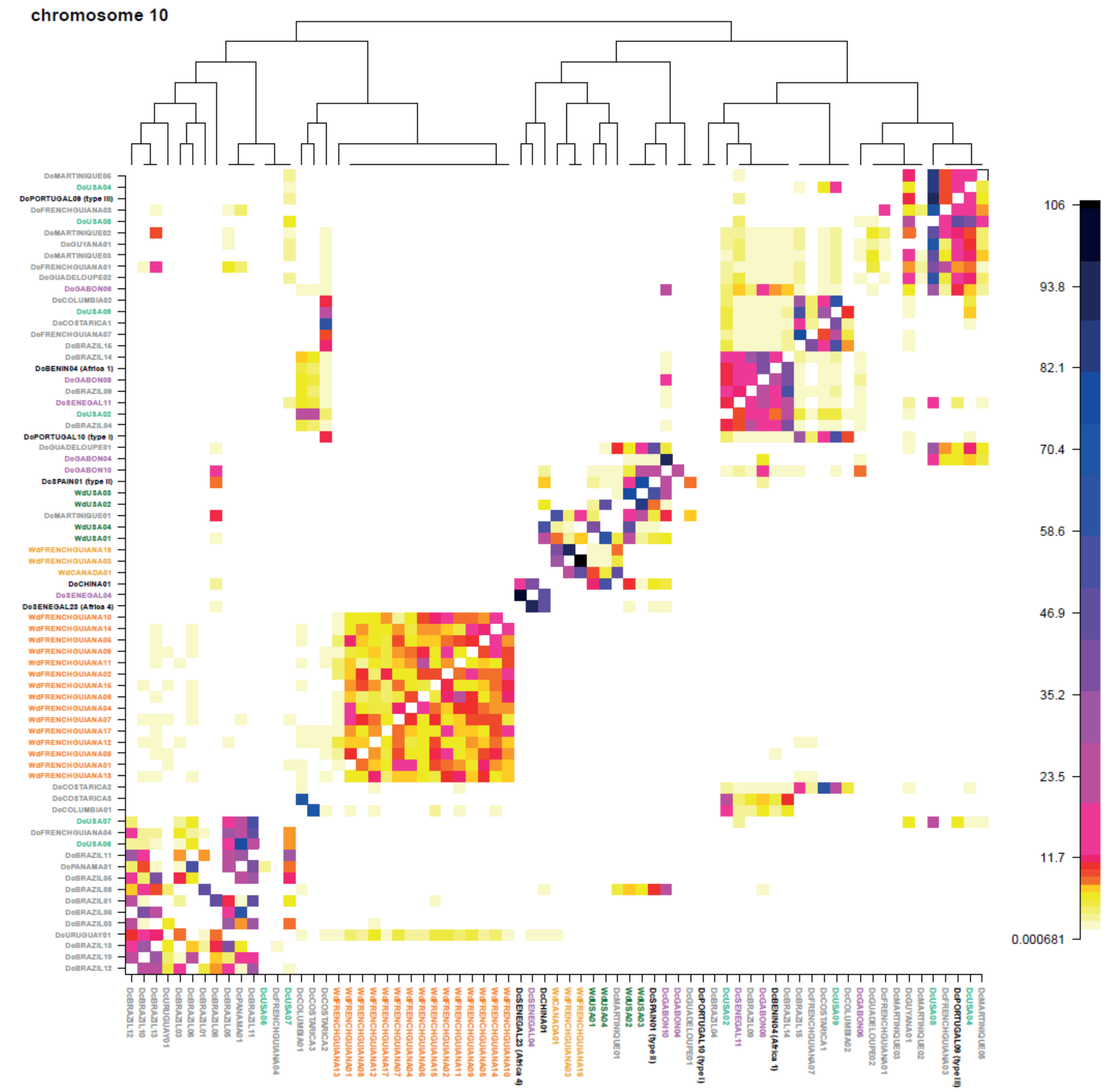

*Supplementary Fig. 2l.*

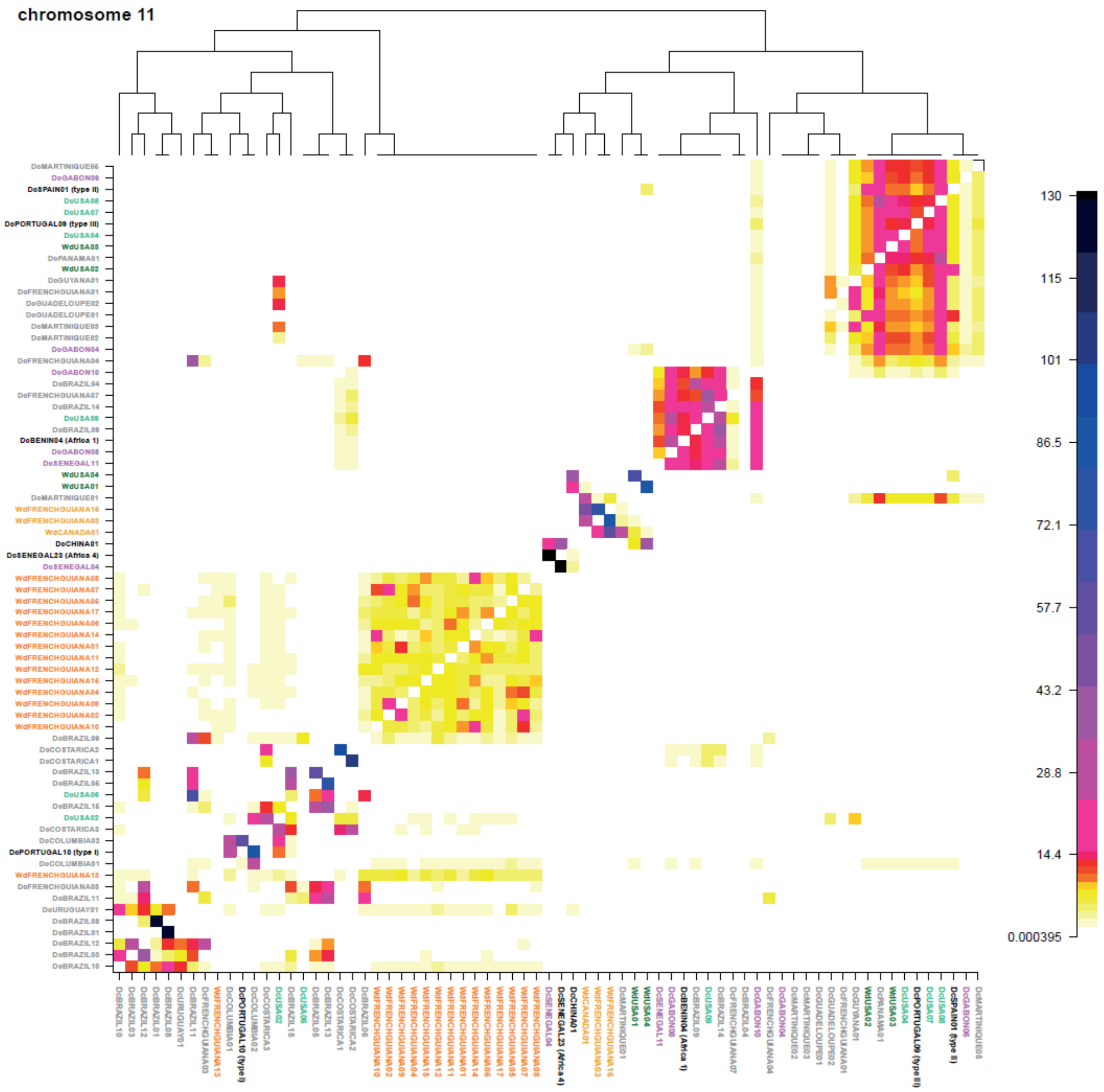

*Supplementary Fig. 2m.*

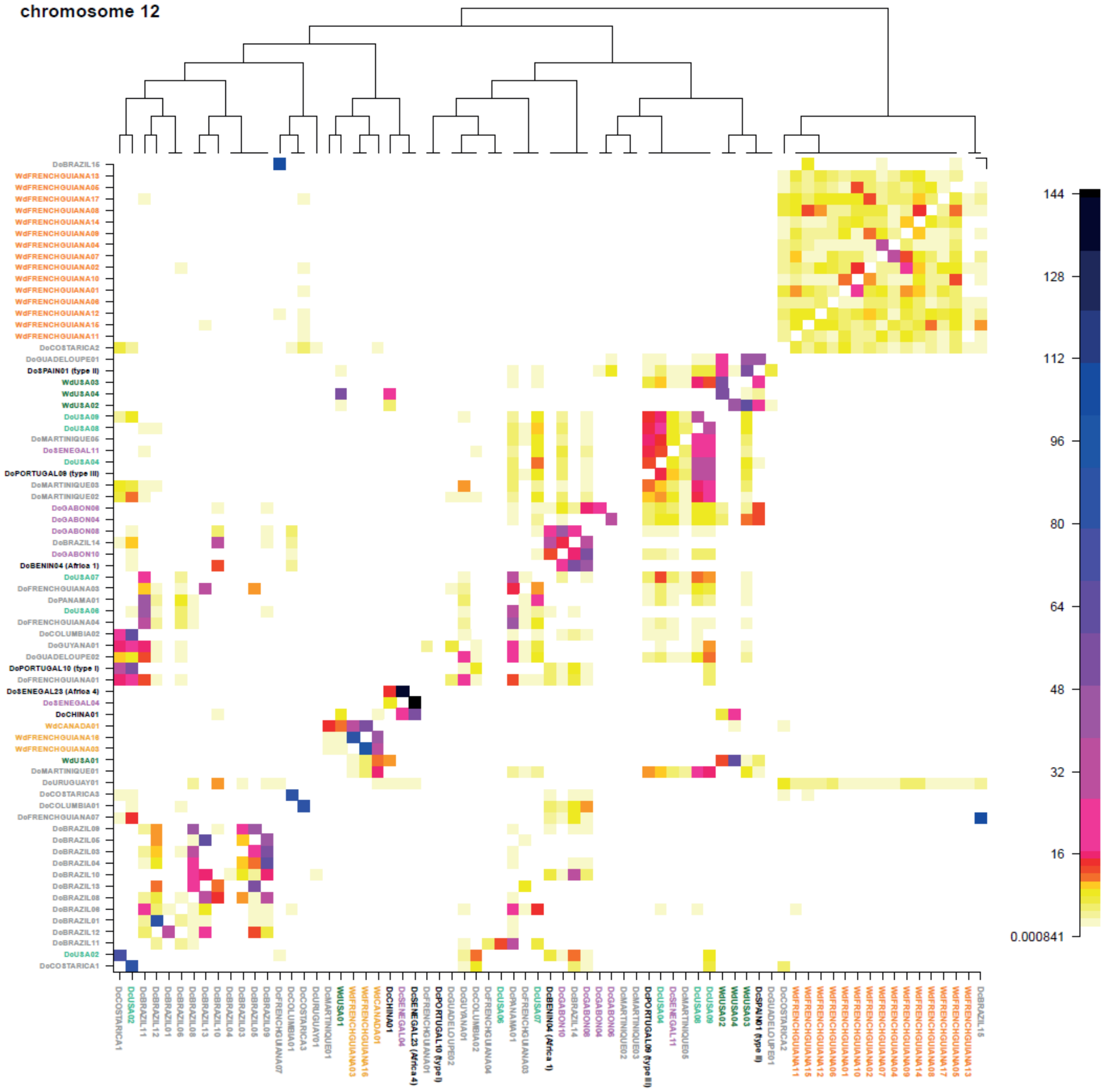

***Supplementary Fig. 2n.***

**Supplementary Fig. 2. ChromoPainter co-ancestry matrices for *Toxoplasma gondii* genomes with population structure assignment based on fineSTRUCTURE analysis.** Co-ancestry matrices were generated for the genome-wide dataset of 71 strains and 588,777 SNPs (a), then for each chromosome independantly (b-n).The colour of each cell of the matrix indicates the expected number of genetic material (chunks) copied from a donor *T. gondii* genome (x-axis) to a recipient genome (y-axis). On the top is the maximum a posteriori (MAP) tree generated by fineSTRUCTURE which shows the groupings of the different populations. The colour legend of IDs is identical to that of ADMIXTURE plots.

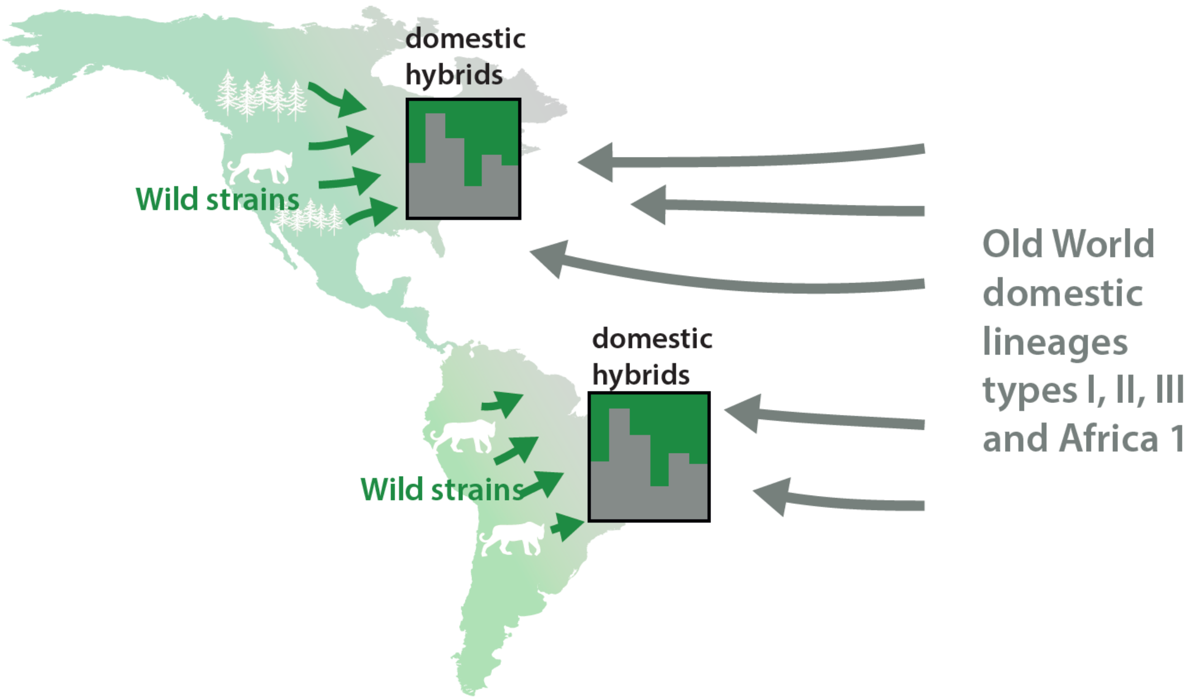

**Supplementary Fig. 3. Cartoon figure representing the presumed hybridization process behind emergence of domestic populations of *T. gondii* in the New World.**

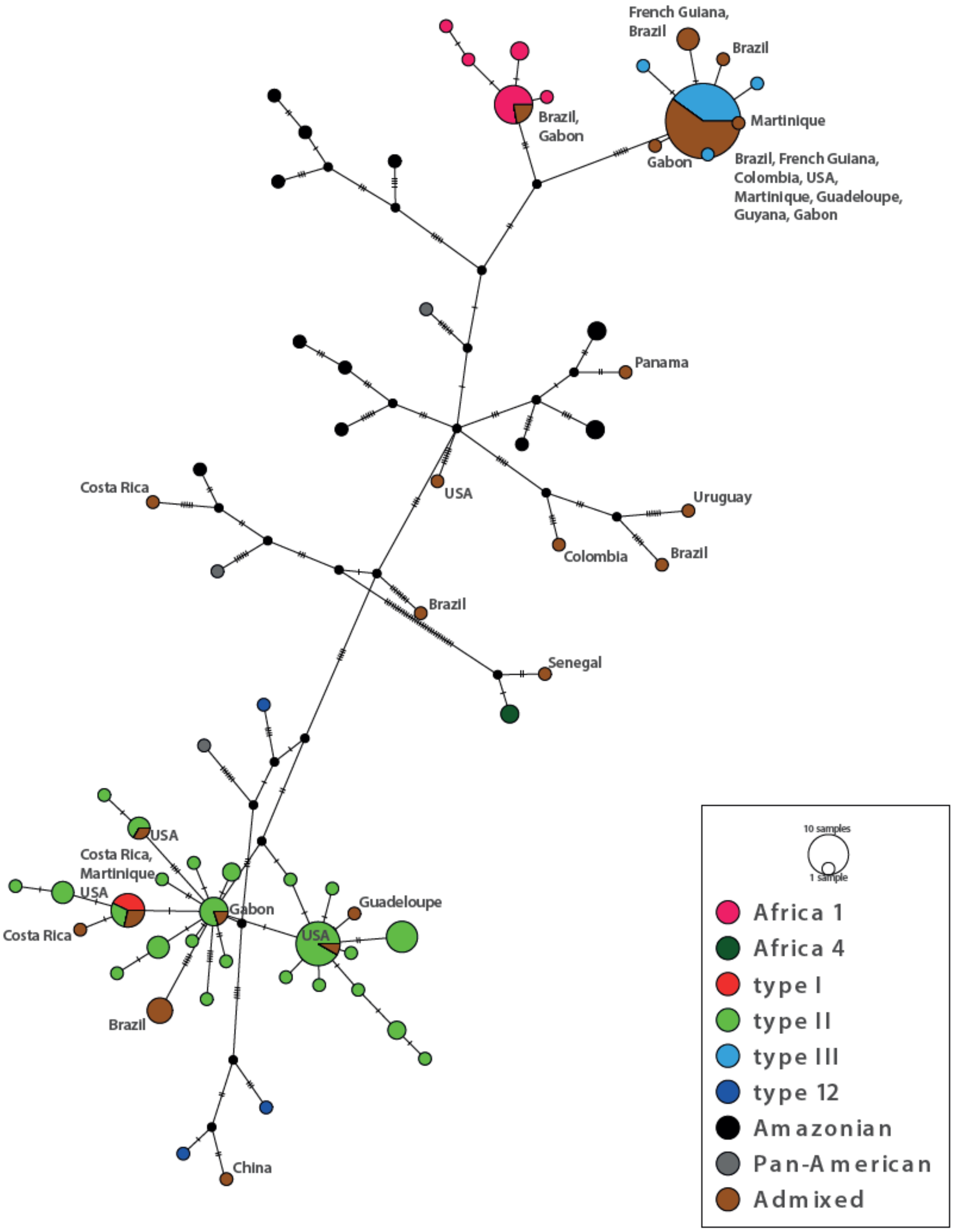

**Supplementary Fig. 4. TCS network of apicoplast sequences.** Only countries of origin of putative hybrids (in brown) are shown.

#### Supplementary Table 1. *Toxoplasma gondii* strains description: hosts, geographical origins, ecotypes and whole-genome sequencing depth.

| Strain name | Alternative names | BRC identifiers | WGS sequencing | Mean sequencing depth | Continent of origin | Country of origin | City or region of origin | Ecotype | Host | Year |
|---|---|---|---|---|---|---|---|---|---|---|
| WdUSA01 | ARI | | Lorenzi et al., 2016 | 29,5 | North America | USA | NA | wild[d] | Human (*Homo sapiens*) | 1992 |
| WdUSA02 | B41 | | Lorenzi et al., 2016 | 28,3 | North America | USA | NA | wild | Bear (*Ursus arctos*) | 1994 |
| WdUSA03 | B73 | | Lorenzi et al., 2016 | 93,3 | North America | USA | NA | wild | Bear (*Ursus arctos*) | 1994 |
| DcBENIN01 | P637S1AJ14 | TgA119001 | this study | 11,6 | Africa | Benin | Natitingou | domestic | Chicken (*Gallus domesticus*) | 2018 |
| DcBENIN02 | P19S1AJ6 | TgA119002 | this study | 15,2 | Africa | Benin | Cotonou | domestic | Chicken (*Gallus domesticus*) | 2018 |
| DcBENIN03 | P193S1CJ28 | TgA119003 | this study | 25,6 | Africa | Benin | Ouidah | domestic | Chicken (*Gallus domesticus*) | 2018 |
| DcBENIN04 | P82S1AJ6 | TgA119004 | this study | 31,0 | Africa | Benin | Cotonou | domestic | Chicken (*Gallus domesticus*) | 2018 |
| DcBENIN05 | P63S1AJ6 | TgA119005 | this study | 7,9 | Africa | Benin | Cotonou | domestic | Chicken (*Gallus domesticus*) | 2018 |
| BOF | BOF ; BE-BOF | | Lorenzi et al., 2016 | 28,6 | Africa | Unknown | NA | Unknown | Human (*Homo sapiens*) | 1993 |
| DcBRAZIL01 | BRA-FEL-CAT001 | TgA115001 | this study | 19,6 | South America (include Central America) | Brazil | Rio de Janeiro | domestic | Cat (*Felis catis*) | 2014 |
| DcUSA02 | CAST ; US-CAST | TgH00008 | Lorenzi et al., 2016 | 59,7 | North America | USA | NA | domestic | Human (*Homo sapiens*) | 1988 |
| DcURUGUAY01 | CASTELLS | | Lorenzi et al., 2016 | 34,1 | South America (include Central America) | Uruguay | NA | domestic | Sheep (*Ovis aries*) | 1993 |
| DcCAMEROON01 | BCB007-MOU | TgH42007A | this study | 17,7 | Africa | Cameroon | NA | domestic | Human (*Homo sapiens*) | 2017 |
| DcDROC01 | PSP026-2005-MUP | TgH29026A | this study | 24,6 | Africa | Democratic Republic of Congo | NA | domestic | Human (*Homo sapiens*) | 2005 |
| WdCANADA01 | COUG ; TgCgCa01 | | Lorenzi et al., 2016 | 32,7 | North America | Canada | NA | wild | Cougar (*Puma concolor*) | 1996 |
| DcSPAIN01 | 170329Lmic01ES | TgA21067 | this study | 28,2 | Europe | Spain | Ebro Delta | domestic | Yellow-legged gull (*Larus michahellis*) | 2017 |
| FOU | FOU ; BRE-FOU | TgH00007 | Lorenzi et al., 2016 | 15,6 | Africa | Unknown | NA | Unknown | Human (*Homo sapiens*) | 1992 |
| DcFRANCE01 | FR-Gal dom-006 | TgA21017 | this study | 22,8 | Europe | France | Le Havre | domestic | Chicken (*Gallus domesticus*) | 2006 |
| DcFRANCE02 | FR-Gal dom-007 | TgA21018 | this study | 15,8 | Europe | France | Le Havre | domestic | Chicken (*Gallus domesticus*) | 2006 |
| DcFRANCE03 | FR-Vul vul-017 | TgA21043 | this study | 18,5 | Europe | France | Haute-Vienne | domestic[a] | Fox (*Vulpes vulpes*) | 2012 |
| DcFRANCE04 | FR-Sus scr-063 | TgA21053 | this study | 12,8 | Europe | France | Camargue | domestic | Pig (*Sus scrofa domesticus*) | 2016 |
| DcFRANCE05 | 190201Bbut002 | TgA21071 | this study | 24,0 | Europe | France | Bordeaux | domestic[a] | Common buzzard *(Buteo buteo)* | 2019 |
| DcFRANCE06 | 190312Rnor01 | TgA21072 | this study | 18,8 | Europe | France | Bordeaux | domestic | Brown rat (*Rattus norvegicus*) | 2019 |
| DcFRANCE07 | 190419Bbut001 | TgA21075 | this study | 15,6 | Europe | France | Bordeaux | domestic[a] | Common buzzard *(Buteo buteo)* | 2019 |
| DcFRANCE08 | FR-Cap cap-009 | TgA32011 | this study | 14,0 | Europe | France | Marne | domestic | Goat (Capra aegagrus hircus) | 2007 |
| DcFRANCE09 | FR-Cap cap-012 | TgA32014 | this study | 17,8 | Europe | France | Seine-et-Marne | domestic[a] | European roe deer (*Capreolus capreolus*) | 2008 |
| DcFRANCE10 | FR-Ond zib002 | TgA32032 | this study | 14,5 | Europe | France | Marne | domestic | Sheep (*Ovis aries*) | 2004 |
| DcFRANCE11 | FR-Sus scr-015 | TgA32067 | this study | 21,1 | Europe | France | Corse | domestic[a] | Boar (*Sus scrofa*) | 2007 |
| DcFRANCE12 | FR-Ovi ari-011 | TgA32078 | this study | 19,5 | Europe | France | Reims | domestic | Sheep (*Ovis aries*) | 2006 |
| DcFRANCE13 | FR-Bos tau-001 | TgA32127 | this study | 24,3 | Europe | France | NA | domestic | Cattle (*Bos taurus*) | 2009 |
| DcFRANCE14 | FR-Vul vul-031 | TgA32154 | this study | 19,3 | Europe | France | Nancy | domestic[a] | Fox (*Vulpes vulpes*) | 2012 |
| DcFRANCE15 | FR-Sus scr-027 | TgA32175 | this study | 18,6 | Europe | France | Rennes | domestic | Pig (*Sus scrofa domesticus*) | 2013 |
| DcFRANCE16 | FR-Sus scr-034 | TgA32182 | this study | 24,1 | Europe | France | NA | domestic | Pig (*Sus scrofa domesticus*) | NA |
| DcFRANCE17 | FR-Sus scr-035 | TgA32183 | this study | 19,9 | Europe | France | NA | domestic | Pig (*Sus scrofa domesticus*) | NA |
| DcFRANCE18 | FR-Sus scr-039 | TgA32187 | this study | 8,8 | Europe | France | NA | domestic | Pig (*Sus scrofa domesticus*) | NA |
| DcFRANCE19 | FR-Sus scr-047 | TgA32197 | this study | 17,8 | Europe | France | Rennes | domestic | Pig (*Sus scrofa domesticus*) | 2013 |
| DcFRANCE20 | FR-Sus-scr-062 | TgA32215 | this study | 24,7 | Europe | France | NA | domestic | Pig (*Sus scrofa domesticus*) | NA |
| DcFRANCE21 | FR-Vul vul-056 | TgA32248 | this study | 17,9 | Europe | France | Nancy | domestic[a] | Fox (*Vulpes vulpes*) | 2015 |
| DcFRANCE22 | FR-Vul vul-068 | TgA32260 | this study | 18,3 | Europe | France | Nancy | domestic[a] | Fox (*Vulpes vulpes*) | 2015 |
| DcFRANCE23 | CAE013-PIR | TgH12013A | this study | 25,3 | Europe | France | NA | domestic | Human (*Homo sapiens*) | 2007 |
| DcFRANCE24 | CCH058-2009-GRO | TgH13058A | this study | 19,1 | Europe | France | NA | domestic | Human (*Homo sapiens*) | 2009 |
| DcPANAMA01 | G662M | | Lorenzi et al., 2016 | 42,9 | South America (include Central America) | Panama | NA | domestic | Ruddy ground dove (*Columbina talpacoti*) | 1992 |
| DcGABON08 | GAB3-2007-GAL-DOM2 | TgA105001 | Lorenzi et al., 2016 | 64,0 | Africa | Gabon | Libreville | domestic | Chicken (*Gallus domesticus*) | 2007 |
| DcGABON09 | GAB5-2007-GAL-DOM1 | TgA105002 | Lorenzi et al., 2016 | 61,9 | Africa | Gabon | Franceville | domestic | Chicken (*Gallus domesticus*) | 2007 |
| DcGABON10 | GAB1-2007-GAL-DOM10 | TgA105003 | Lorenzi et al., 2016 | 67,0 | Africa | Gabon | Dienga | domestic | Chicken (*Gallus domesticus*) | 2007 |
| DcGABON11 | GAB2-2007-GAL-DOM2 | TgA105004 | Lorenzi et al., 2016 | 83,7 | Africa | Gabon | Makokou | domestic | Chicken (*Gallus domesticus*) | 2007 |
| DcGABON12 | GAB3-2007-GAL-DOM9 | TgA105005 | Lorenzi et al., 2016 | 49,4 | Africa | Gabon | Libreville | domestic | Chicken (*Gallus domesticus*) | 2007 |
| DcGABON13 | GAB5-2007-GAL-DOM6 | TgA105006 | Lorenzi et al., 2016 | 38,0 | Africa | Gabon | Franceville | domestic | Chicken (*Gallus domesticus*) | 2007 |
| DcGABON01 | GAB1-CAP-AEG007 | TgA105012 | this study | 11,3 | Africa | Gabon | Dienga | domestic | Goat (Capra aegagrus hircus) | 2007 |
| DcGABON02 | GAB1-2007-CAP-AEG004 | TgA105033 | this study | 15,4 | Africa | Gabon | Dienga | domestic | Goat (Capra aegagrus hircus) | 2007 |
| DcGABON03 | GAB1-FEL-CAT001 | TgA105034 | this study | 14,8 | Africa | Gabon | Dienga | domestic | Cat (*Felis catis*) | 2007 |
| DcGABON04 | GAB2-2007-GAL-DOM006 | TgA105040 | this study | 16,6 | Africa | Gabon | Makokou | domestic | Chicken (*Gallus domesticus*) | 2007 |
| DcGABON05 | GAB3-GAL-DOM013 | TgA105045 | this study | 20,2 | Africa | Gabon | Libreville | domestic | Chicken (*Gallus domesticus*) | 2007 |
| DcGABON06 | GAB4-2007-GAL-DOM001 | TgA105053 | this study | 20,4 | Africa | Gabon | La Lopé | domestic | Chicken (*Gallus domesticus*) | 2007 |
| DcGABON07 | GAB7-GAL-DOM007 | TgA105057 | this study | 12,9 | Africa | Gabon | Leconi | domestic | Chicken (*Gallus domesticus*) | 2007 |
| DcGUADELOUPE01 | PAP001-BS | TgH40001A | this study | 14,6 | South America (include Central America) | Guadeloupe (French Overseas department) | NA | domestic[b] | Human (*Homo sapiens*) | 2009 |
| DcGUADELOUPE02 | PAP002-2010-GOM | TgH40002A | this study | 20,6 | South America (include Central America) | Guadeloupe (French Overseas department) | NA | domestic | Human (*Homo sapiens*) | 2010 |
| DcUK01 | NAN033-2005-PLA | TgH26033A | this study | 24,8 | Europe | United Kingdom | NA | domestic | Human (*Homo sapiens*) | 2005 |
| DcUSA03 | GT1 | TgA00004 | Lorenzi et al., 2016 | 67,9 | North America | USA | NA | domestic | Goat (*Capra aegagrus hircus*) | 1980 |
| WdFRENCHGUIANA01 | GUY-GAL-VIT-001 | TgA18005 | this study | 20,8 | South America (include Central America) | French Guiana (French Overseas department) | Cayenne | wild | Greater grison (*Galatis vitata*) | 2009 |
| WdFRENCHGUIANA02 | GUY-CAN-FAM-007 | TgA18006 | this study | 22,9 | South America (include Central America) | French Guiana (French Overseas department) | Roura | wild[c] | Dog (*Canis familiaris*) | 2009 |
| DcFRENCHGUIANA01 | GUY-CAN-FAM-009 | TgA18009 | this study | 16,7 | South America (include Central America) | French Guiana (French Overseas department) | Cayenne | domestic | Dog (*Canis familiaris*) | 2009 |
| DcFRENCHGUIANA02 | GUY-CAN-FAM-018 | TgA18020 | this study | 12,8 | South America (include Central America) | French Guiana (French Overseas department) | Matoury | domestic | Dog (*Canis familiaris*) | 2009 |
| DcFRENCHGUIANA03 | GUY-CAN-FAM-019 | TgA18030 | this study | 25,0 | South America (include Central America) | French Guiana (French Overseas department) | Matoury | domestic | Dog (*Canis familiaris*) | 2009 |
| DcFRENCHGUIANA04 | GUY-CAN-FAM-003 | TgA18031 | this study | 17,5 | South America (include Central America) | French Guiana (French Overseas department) | Macouria | domestic | Dog (*Canis familiaris*) | 2009 |
| WdFRENCHGUIANA03 | GUY-MAZ-OUA-001 | TgA18032 | this study | 14,4 | South America (include Central America) | French Guiana (French Overseas department) | Regina | wild | Gray brocket (*Mazama gouazoubira*) | 2009 |
| DcFRENCHGUIANA05 | GUY-CAN-FAM-008 | TgA18033 | this study | 13,9 | South America (include Central America) | French Guiana (French Overseas department) | Montjoly | domestic | Dog (*Canis familiaris*) | 2009 |
| DcFRENCHGUIANA06 | GUY-FEL-CAT-009 | TgA18034 | this study | 11,2 | South America (include Central America) | French Guiana (French Overseas department) | Matoury | domestic | Cat (*Felis catis*) | 2009 |
| WdFRENCHGUIANA09 | GUY-DOS | TgH18001 | Lorenzi et al., 2016 | 28,9 | South America (include Central America) | French Guiana (French Overseas department) | NA | wild | Human (*Homo sapiens*) | 2001 |
| WdFRENCHGUIANA10 | GUY-KOE | TgH18002 | Lorenzi et al., 2016 | 25,3 | South America (include Central America) | French Guiana (French Overseas department) | NA | wild | Human (*Homo sapiens*) | 2002 |
| WdFRENCHGUIANA11 | GUY-MAT | TgH18003 | Lorenzi et al., 2016 | 31,2 | South America (include Central America) | French Guiana (French Overseas department) | NA | wild | Human (*Homo sapiens*) | 2002 |
| WdFRENCHGUIANA12 | GUY-2003-MEL | TgH18007 | Lorenzi et al., 2016 | 40,3 | South America (include Central America) | French Guiana (French Overseas department) | NA | wild | Human (*Homo sapiens*) | 2003 |
| WdFRENCHGUIANA13 | GUY-2004-ABE | TgH18008 | Lorenzi et al., 2016 | 59,7 | South America (include Central America) | French Guiana (French Overseas department) | NA | wild | Human (*Homo sapiens*) | 2004 |
| WdFRENCHGUIANA14 | GUY009-AKO | TgH18009 | Lorenzi et al., 2016 | 79,5 | South America (include Central America) | French Guiana (French Overseas department) | NA | wild | Human (*Homo sapiens*) | 2004 |
| WdFRENCHGUIANA04 | GUY013-2004-LAB | TgH18013A | this study | 17,1 | South America (include Central America) | French Guiana (French Overseas department) | NA | wild | Human (*Homo sapiens*) | 2009 |
| WdFRENCHGUIANA15 | GUY 021 - TOJ | TgH18021 | Lorenzi et al., 2016 | 39,0 | South America (include Central America) | French Guiana (French Overseas department) | NA | wild | Human (*Homo sapiens*) | 2006 |
| DcFRENCHGUIANA07 | GUY057-GEN | TgH18057A | this study | 21,3 | South America (include Central America) | French Guiana (French Overseas department) | NA | domestic | Human (*Homo sapiens*) | 2009 |
| WdFRENCHGUIANA06 | GUY066-MON | TgH18066A | this study | 17,3 | South America (include Central America) | French Guiana (French Overseas department) | NA | wild | Human (*Homo sapiens*) | 2009 |
| WdFRENCHGUIANA07 | GUY071-GUY | TgH18071A | this study | 25,3 | South America (include Central America) | French Guiana (French Overseas department) | NA | wild | Human (*Homo sapiens*) | 2009 |
| WdFRENCHGUIANA08 | GUYS006-BAY | TgH19006A | this study | 17,5 | South America (include Central America) | French Guiana (French Overseas department) | NA | wild | Human (*Homo sapiens*) | 2009 |
| WdFRENCHGUIANA16 | GUY-JAG1 | TgA18001 | Lorenzi et al., 2016 | 26,7 | South America (include Central America) | French Guiana (French Overseas department) | Regina | wild | Jaguar*(Panthera onca)* | 2004 |
| LGE-CUV | LGE-CUV | TgH21016 | Lorenzi et al., 2016 | 76,4 | Unknown | Unknown | NA | Unknown | Human (*Homo sapiens*) | 2007 |
| DcUSA04 | M7741 | | Lorenzi et al., 2016 | 77,7 | North America | USA | NA | domestic | Sheep (*Ovis aries*) | 1958 |
| MAS | MAS ; LPN-MAS | TgH00006 | Lorenzi et al., 2016 | 32,7 | Unknown | Unknown | NA | Unknown | Human (*Homo sapiens*) | 1991 |
| DcUSA05 | ME49 | | Lorenzi et al., 2016 | 631,7 | North America | USA | NA | domestic | Sheep (*Ovis aries*) | 1965 |
| DcMARTINIQUE01 | FDF001-ANG | TgH16001A | this study | 15,4 | South America (include Central America) | Martinique (French Overseas department) | NA | domestic | Human (*Homo sapiens*) | 2007 |
| DcMARTINIQUE02 | FDF010-ANS | TgH16010A | this study | 14,7 | South America (include Central America) | Martinique (French Overseas department) | NA | domestic | Human (*Homo sapiens*) | 2013 |
| DcMARTINIQUE03 | FDF012-MAN | TgH16012A | this study | 13,1 | South America (include Central America) | Martinique (French Overseas department) | NA | domestic | Human (*Homo sapiens*) | 2014 |
| DcMARTINIQUE04 | FDF012-DEA | TgH16013B | this study | 15,7 | South America (include Central America) | Martinique (French Overseas department) | NA | domestic | Human (*Homo sapiens*) | 2017 |
| DcMARTINIQUE05 | FDF014-NEB | TgH16014A | this study | 23,1 | South America (include Central America) | Martinique (French Overseas department) | NA | domestic | Human (*Homo sapiens*) | 2018 |
| DcUSA06 | P89 ; TgPgUs15 | TgA00005 | Lorenzi et al., 2016 | 12,5 | North America | USA | NA | domestic | Pig (*Sus scrofa domesticus*) | 1991 |
| DcCHINA01 | PgCatPRC2 | | Lorenzi et al., 2016 | 22,3 | Asia | China | NA | domestic | Cat (*Felis catis*) | 2007 |
| DcPORTUGAL01 | PT-2004-SUS SCR001 | TgA103006 | this study | 27,6 | Europe | Portugal | Vinhais | domestic | Pig (*Sus scrofa domesticus*) | 2004 |
| DcPORTUGAL02 | PT-SUS SCR002 | TgA103007 | this study | 13,2 | Europe | Portugal | Vinhais | domestic | Pig (*Sus scrofa domesticus*) | 2004 |
| DcPORTUGAL03 | PT-2005-SUS SCR004 | TgA103009 | this study | 15,7 | Europe | Portugal | Vinhais | domestic | Pig (*Sus scrofa domesticus*) | 2004 |
| DcPORTUGAL04 | PT-SUS SCR005 | TgA103010 | this study | 26,3 | Europe | Portugal | Vinhais | domestic | Pig (*Sus scrofa domesticus*) | 2004 |
| DcPORTUGAL05 | PT-2005-SUS SCR006 | TgA103011 | this study | 22,7 | Europe | Portugal | Vinhais | domestic | Pig (*Sus scrofa domesticus*) | 2004 |
| DcPORTUGAL06 | PT-2005-SUS SCR007 | TgA103012 | this study | 17,8 | Europe | Portugal | Vinhais | domestic | Pig (*Sus scrofa domesticus*) | 2004 |
| DcPORTUGAL07 | PT-2005-SUS SCR 11 | TgA103016 | this study | 21,9 | Europe | Portugal | Vinhais | domestic | Pig (*Sus scrofa domesticus*) | 2004 |
| DcPORTUGAL08 | PT-2005-SUS SCR013 | TgA103018 | this study | 19,5 | Europe | Portugal | Vinhais | domestic | Pig (*Sus scrofa domesticus*) | 2004 |
| DcPORTUGAL09 | PT-SUS SCR014 | TgA103019 | this study | 27,4 | Europe | Portugal | Vinhais | domestic | Pig (*Sus scrofa domesticus*) | 2004 |
| DcPORTUGAL10 | PT-B1 | TgH103001 | this study | 24,2 | Europe | Portugal (Acores) | NA | domestic | Cattle (*Bos taurus*) | 2002 |
| WdUSA04 | RAY | | Lorenzi et al., 2016 | 14,8 | North America | USA | NA | wild[d] | Human (*Homo sapiens*) | 1993 |
| DcUSA07 | ROD ; ROD-US | | Lorenzi et al., 2016 | 78,8 | North America | USA | NA | domestic | Human (*Homo sapiens*) | 1993 |
| WdFRENCHGUIANA17 | RUB ; GUY-RUB | TgH00002 | Lorenzi et al., 2016 | 49,6 | South America (include Central America) | French Guiana (French Overseas department) | NA | wild | Human (*Homo sapiens*) | 1991 |
| DcSENEGAL01 | 160817Gdom21 | TgA117003 | this study | 31,5 | Africa | Senegal | Kedougou | domestic | Chicken (*Gallus domesticus*) | 2016 |
| DcSENEGAL02 | 160823Gdom12 | TgA117004 | this study | 38,1 | Africa | Senegal | Goree island | domestic | Chicken (*Gallus domesticus*) | 2016 |
| DcSENEGAL03 | 180218Gdom96 | TgA117005 | this study | 16,9 | Africa | Senegal | Saint-Louis | domestic | Chicken (*Gallus domesticus*) | 2018 |
| DcSENEGAL04 | 160504Fcat01 | TgA117007 | this study | 13,1 | Africa | Senegal | Dakar | domestic | Cat (*Felis catis*) | 2016 |
| DcSENEGAL05 | 160817Gdom23 | TgA117008 | this study | 17,1 | Africa | Senegal | Kedougou | domestic | Guinea Fowl (*Numida meleagris*) | 2016 |
| DcSENEGAL06 | 160730Gdom13 | TgA117012 | this study | 26,6 | Africa | Senegal | Saint-Louis | domestic | Chicken (*Gallus domesticus*) | 2016 |
| DcSENEGAL07 | 160421Gdom10 | TgA117015 | this study | 57,1 | Africa | Senegal | Dakar | domestic | Chicken (*Gallus domesticus*) | 2016 |
| DcSENEGAL08 | 160416Gdom13 | TgA117016 | this study | 20,1 | Africa | Senegal | Dakar | domestic | Chicken (*Gallus domesticus*) | 2016 |
| DcSENEGAL09 | 160421Gdom18 | TgA117019 | this study | 18,5 | Africa | Senegal | Dakar | domestic | Chicken (*Gallus domesticus*) | 2016 |
| DcSENEGAL10 | 160428Gdom06 | TgA117020 | this study | 11,2 | Africa | Senegal | Dakar | domestic | Chicken (*Gallus domesticus*) | 2016 |
| DcSENEGAL11 | 160510Gdom02 | TgA117022 | this study | 38,7 | Africa | Senegal | Dakar | domestic | Chicken (*Gallus domesticus*) | 2016 |
| DcSENEGAL12 | 160518Gdom05 | TgA117027 | this study | 42,6 | Africa | Senegal | Dakar | domestic | Chicken (*Gallus domesticus*) | 2016 |
| DcSENEGAL13 | 160529Gdom26 | TgA117028 | this study | 32,7 | Africa | Senegal | Saint-Louis | domestic | Chicken (*Gallus domesticus*) | 2016 |
| DcSENEGAL14 | 160530Cmos12 | TgA117032 | this study | 11,9 | Africa | Senegal | Saint-Louis | domestic | Duck (*Cairina moschata*) | 2016 |
| DcSENEGAL15 | 160614Gdom21 | TgA117036 | this study | 34,3 | Africa | Senegal | Dakar | domestic | Chicken (*Gallus domesticus*) | 2016 |
| DcSENEGAL16 | 160615Gdom02 | TgA117037 | this study | 39,1 | Africa | Senegal | Dakar | domestic | Chicken (*Gallus domesticus*) | 2016 |
| DcSENEGAL17 | 160622Gdom02 | TgA117041 | this study | 19,3 | Africa | Senegal | Saint-Louis | domestic | Chicken (*Gallus domesticus*) | 2016 |
| DcSENEGAL18 | 160622Gdom38 | TgA117043 | this study | 37,3 | Africa | Senegal | Saint-Louis | domestic | Chicken (*Gallus domesticus*) | 2016 |
| DcSENEGAL19 | 160623Gdom08 | TgA117045 | this study | 34,6 | Africa | Senegal | Saint-Louis | domestic | Chicken (*Gallus domesticus*) | 2016 |
| DcSENEGAL20 | 160623Gdom19 | TgA117046 | this study | 12,0 | Africa | Senegal | Saint-Louis | domestic | Chicken (*Gallus domesticus*) | 2016 |
| DcSENEGAL21 | 160625Gdom11 | TgA117048 | this study | 54,0 | Africa | Senegal | Saint-Louis | domestic | Chicken (*Gallus domesticus*) | 2016 |
| DcSENEGAL22 | 160628Gdom17 | TgA117052 | this study | 31,8 | Africa | Senegal | Saint-Louis | domestic | Chicken (*Gallus domesticus*) | 2016 |
| DcSENEGAL23 | 160628Gdom28 | TgA117053 | this study | 24,0 | Africa | Senegal | Saint-Louis | domestic | Chicken (*Gallus domesticus*) | 2016 |
| DcSENEGAL24 | 160628Gdom35 | TgA117054 | this study | 37,6 | Africa | Senegal | Saint-Louis | domestic | Chicken (*Gallus domesticus*) | 2016 |
| DcSENEGAL25 | 160816Gdom37 | TgA117059 | this study | 21,2 | Africa | Senegal | Kedougou | domestic | Chicken (*Gallus domesticus*) | 2016 |
| DcSENEGAL26 | 160823Gdom04 | TgA117062 | this study | 21,5 | Africa | Senegal | Goree island | domestic | Chicken (*Gallus domesticus*) | 2016 |
| DcUSA08 | SOU | | Lorenzi et al., 2016 | 76,6 | North America | USA | NA | domestic | Human (*Homo sapiens*) | 1985 |
| DcSERBIA01 | RS-CTN | TgH106002 | this study | 21,4 | Europe | Serbia | NA | domestic | Human (*Homo sapiens*) | 2008 |
| DcBRAZIL02 | TgCatBr01 | | Lorenzi et al., 2016 | 28,8 | South America (include Central America) | Brazil | Sao Paulo | domestic | Cat (*Felis catis*) | 2006 |
| DcBRAZIL03 | TgCatBr05 | TgA00007 | Lorenzi et al., 2016 | 20,1 | South America (include Central America) | Brazil | Sao Paulo | domestic | Cat (*Felis catis*) | 2006 |
| DcBRAZIL04 | TgCatBr09 | | Lorenzi et al., 2016 | 12,0 | South America (include Central America) | Brazil | Sao Paulo | domestic | Cat (*Felis catis*) | 2006 |
| DcBRAZIL05 | TgCatBr10 | | Lorenzi et al., 2016 | 28,1 | South America (include Central America) | Brazil | Sao Paulo | domestic | Cat (*Felis catis*) | 2006 |
| DcBRAZIL06 | TgCatBr15 | | Lorenzi et al., 2016 | 36,5 | South America (include Central America) | Brazil | Sao Paulo | domestic | Cat (*Felis catis*) | 2006 |
| DcBRAZIL07 | TgCatBr18 | | Lorenzi et al., 2016 | 26,3 | South America (include Central America) | Brazil | Sao Paulo | domestic | Cat (*Felis catis*) | 2006 |
| DcBRAZIL08 | TgCatBr25 | | Lorenzi et al., 2016 | 41,7 | South America (include Central America) | Brazil | Sao Paulo | domestic | Cat (*Felis catis*) | 2006 |
| DcBRAZIL09 | TgCatBr26 | | Lorenzi et al., 2016 | 31,9 | South America (include Central America) | Brazil | Sao Paulo | domestic | Cat (*Felis catis*) | 2006 |
| DcBRAZIL10 | TgCatBr3 | | Lorenzi et al., 2016 | 40,5 | South America (include Central America) | Brazil | Parana | domestic | Cat (*Felis catis*) | 2003 |

| | | | | | | | | | | |
|---|---|---|---|---|---|---|---|---|---|---|
| DcBRAZIL11 | TgCatBr34 | | Lorenzi et al., 2016 | 47,6 | South America (include Central America) | Brazil | Sao Paulo | domestic | Cat (*Felis catis* ) | 2006 |
| DcBRAZIL12 | TgCatBr44 | | Lorenzi et al., 2016 | 43,8 | South America (include Central America) | Brazil | Sao Paulo | domestic | Cat (*Felis catis* ) | 2008 |
| DcBRAZIL13 | TgCatBr64 | | Lorenzi et al., 2016 | 56,6 | South America (include Central America) | Brazil | Sao Paulo | domestic | Cat (*Felis catis* ) | 2003 |
| DcBRAZIL14 | TgCatBr72 | | Lorenzi et al., 2016 | 61,3 | South America (include Central America) | Brazil | Sao Paulo | domestic | Cat (*Felis catis* ) | 2003 |
| DcBRAZIL15 | TgCkBr141 | | Lorenzi et al., 2016 | 34,3 | South America (include Central America) | Brazil | Para | domestic | Chicken (*Gallus domesticus* ) | 2006 |
| DcCOSTARICA1 | TgCkCr01 | | Lorenzi et al., 2016 | 42,8 | South America (include Central America) | Costa Rica | NA | domestic | Chicken (*Gallus domesticus* ) | 2005 |
| DcCOSTARICA2 | TgCkCr10 | | Lorenzi et al., 2016 | 131,4 | South America (include Central America) | Costa Rica | NA | domestic | Chicken (*Gallus domesticus* ) | 2006 |
| DcGUYANA01 | TgCkGy2 | | Lorenzi et al., 2016 | 26,3 | South America (include Central America) | Guyana | NA | domestic | Chicken (*Gallus domesticus* ) | 2007 |
| DcCOLUMBIA01 | TgCtCo5 | | Lorenzi et al., 2016 | 43,8 | South America (include Central America) | Columbia | Bogota | domestic | Cat (*Felis catis* ) | 2006 |
| DcCOLUMBIA02 | TgDgCo17 | | Lorenzi et al., 2016 | 33,0 | South America (include Central America) | Columbia | NA | domestic | Dog (*Canis familiaris* ) | 2006 |
| TgH20005 | IPP-URB | TgH20005 | Lorenzi et al., 2016 | 64,0 | Unknown | Unknown | NA | Unknown | Human (*Homo sapiens* ) | 2006 |
| TgH26044 | | TgH26044 | Lorenzi et al., 2016 | 46,1 | Unknown | Unknown | NA | Unknown | Human (*Homo sapiens* ) | 2007 |
| DcCOSTARICA3 | TgRsCr01 | | Lorenzi et al., 2016 | 36,4 | South America (include Central America) | Costa Rica | NA | domestic[e] | Toucan (*Ramphastos sulfuratus* ) | 2006 |
| DcUSA09 | TgShUs28 | | Lorenzi et al., 2016 | 47,1 | North America | USA | NA | domestic | Sheep (*Ovis aries* ) | 2008 |
| DcTUNISIA01 | TUN-Oviari-066 | TgA104005 | this study | 10,4 | Africa | Tunisia | Monastir | domestic | Sheep (*Ovis aries* ) | 2016 |
| DcTUNISIA02 | TUN-Oviari-071 | TgA104010 | this study | 16,8 | Africa | Tunisia | Monastir | domestic | Sheep (*Ovis aries* ) | 2016 |
| DcTUNISIA03 | TUN-Oviari-074 | TgA104013 | this study | 14,4 | Africa | Tunisia | Monastir | domestic | Sheep (*Ovis aries* ) | 2016 |
| DcTURKEY01 | Ankara LS-1 | TgH102001A | this study | 13,4 | Asia | Turkey | NA | domestic | Human (*Homo sapiens* ) | 1972 |
| DcTURKEY02 | TR-EGE1 LS-2 | TgH102002A | this study | 13,2 | Asia | Turkey | NA | domestic | Human (*Homo sapiens* ) | 2007 |
| DcUSA01 | US-CT1 | TgA00006 | this study | 16,5 | North America | USA | NA | domestic | Cattle (*Bos taurus* ) | 1989 |
| WdFRENCHGUIANA05 | VAND ; GUY-VAND | TgH00009 | Lorenzi et al., 2016 | 21,4 | South America (include Central America) | French Guiana (French Overseas department) | NA | wild | Human (*Homo sapiens* ) | 1997 |
| DcUSA10 | VEG | TgH00005 | Lorenzi et al., 2016 | 70,1 | North America | USA | NA | domestic | Sheep (*Ovis aries* ) | 1988 |

NA: Not available

[a] Domestic T. gondii strains in wild animals: in western Europe, wild felids are nearly extinct, and domestic cats are virtually the only shedders of the oocysts infecting wild birds and mammals. Given the large populations of domestic cats in Europe, billions of oocysts spread over long distances via waterways before reaching wildlife (VanWormer et al., 2013; Gotteland et al., 2014). This is verified by the similarity of the genotypes infecting domestic and wild animals in Europe, all belonging to few domestic T. gondii lineages (mostly type II and type III) (Aubert et al., 2010; Shwab et al., 2018).

[b] The information about the ecotype of origin of each strain was provided by the French Biological Resource Centre (BRC) for Toxoplasma. In contrast to animal strains for which determining the ecotype of origin is often straightforward as it is based on the host species ecology, humans are facing different exposure situations, and tracing back the origin of the infection is sometimes difficult. The BRC works through a network of correspondents who send human isolates to the French National Reference Center for Toxoplasmosis (parasitology departments of Reims and Limoges University hospitals, France), with associated metadata about patient history (patients reporting forest-related activities such as ingestion of surface water, consumption of undercooked game meat, and hunting…).

[c] This strain, genetically characterized as belonging to the Amazonian wild group, was isolated from a stray dog from Roura, an area surrounded by the Amazon rainforest. It is likely that this dog was infected by this wild strain after consumption of wild game or running water in the forest or after coming into contact with contaminated forest soil (Mercier et al., 2011).

[d] There is no available data on the ecotype of origin (domestic or wild) for this strain. However, it belongs to type 12, a North American lineage found mainly in wildlife (Khan et al., 2011).

[e] Although being a wild bird, this Toucan got infected in a zoo, in the heart of Santa José, the capital of Costa Rica (Dubey et al., 2009).

#### Supplementary Table 2. *Toxoplasma gondii* multilocus genotypes

| Strain name | Multilocus Microsatellite classification | TUB2 | W35 | TgM-A | B18 | B17 | M33 | MIV.1 | MXI.1 | M48 | M102 | N60 | N82 | AA | N61 | N83 | RFLP genotype |
|---|---|---|---|---|---|---|---|---|---|---|---|---|---|---|---|---|---|
| WdUSA01 | Type 12 | 289 | 242 | 209 | 158 | 336 | 169 | 274 | 362 | 215 | 170 | 147 | 131 | 295 | 89 | 316 | 5 |
| WdUSA02 | Type 12 | 289 | 242 | 207 | 162 | 336 | 169 | 274 | 356 | 213 | 170 | 142 | 111 | 287 | 107 | 316 | 4 |
| WdUSA03 | unclassified genotype | 289 | 242 | 207 | 160 | 336 | 169 | 274 | 356 | 213 | 190 | 142 | 111 | 267 | 101 | 310 | 127 |
| DcBENIN01 | Africa 1 | 291 | 248 | 205 | 160 | 342 | 165 | 274 | 354 | 229 | 166 | 147 | 111 | 271 | 89 | 308 | 6 |
| DcBENIN02 | Africa 1 | 291 | 248 | 205 | 160 | 342 | 165 | 274 | 354 | 227 | 166 | 147 | 111 | 267 | 89 | 306 | 6 |
| DcBENIN03 | Type III | 289 | 242 | 205 | 160 | 336 | 165 | 278 | 356 | 213 | 190 | 151 | 111 | 265 | 089 | 312 | 2 |
| DcBENIN04 | Africa 1 | 291 | 248 | 205 | 160 | 342 | 165 | 274 | 354 | 233 | 166 | 147 | 109 | 267 | 89 | 306 | 6 |
| DcBENIN05 | Africa 1 | 291 | 248 | 205 | 160 | 342 | 165 | 274 | 354 | 239 | 166 | 149 | 111 | 269 | 91 | 306 | 6 |
| BOF | Africa 1 | 291 | 248 | 205 | 160 | 342 | 165 | 274 | 354 | 227 | 166 | 147 | 111 | 273 | 89 | 306 | 6 |
| DcBRAZIL01 | unclassified genotype | 289 | 242 | 205 | 160 | 342 | 165 | 278 | 358 | 231 | 164 | 145 | 111 | 316 | 89 | 308 | NA |
| DcUSA02 | unclassified genotype | 291 | 242 | 205 | 158 | 342 | 167 | 276 | 356 | 211 | 168 | 147 | 119 | 279 | 87 | 306 | 28 |
| DcURUGUAY01 | unclassified genotype | 287 | 242 | 207 | 158 | 358 | 169 | 274 | 356 | 239 | 164 | 138 | 109 | 283 | 87 | 324 | 15 |
| DcCAMEROON01 | Africa 1 | 291 | 248 | 205 | 160 | 342 | 165 | 274 | 354 | 225 | 166 | 151 | 111 | 273 | 89 | 306 | 6 |
| DcDROC01 | Type III | 289 | 242 | 205 | 160 | 336 | 165 | 278 | 356 | 211 | 190 | 145 | 111 | 267 | 87 | 312 | 2 |
| WdCANADA01 | unclassified genotype | 289 | 242 | 205 | 158 | 336 | 169 | 274 | 354 | 219 | 174 | 151 | 119 | 259 | 79 | 332 | 66 |
| DcSPAIN01 | Type II | 289 | 242 | 207 | 158 | 336 | 169 | 274 | 356 | 221 | 178 | 145 | 115 | 263 | 119 | 308 | 1 or 3 |
| FOU | Africa 1 | 291 | 248 | 205 | 160 | 342 | 165 | 274 | 354 | 227 | 166 | 147 | 111 | 281 | 89 | 306 | 6 |
| DcFRANCE01 | Type II | 289 | 242 | 207 | 158 | 336 | 169 | 274 | 356 | 233 | 176 | 140 | 111 | 261 | 93 | 310 | 1 or 3 |
| DcFRANCE02 | Type II | 289 | 242 | 207 | 158 | 336 | 169 | 274 | 356 | 213 | 172 | 142 | 111 | 265 | 107 | 310 | 1 or 3 |
| DcFRANCE03 | Type II | 289 | 242 | 207 | 158 | 336 | 169 | 274 | 356 | 215 | 178 | 140 | 111 | 267 | 95 | 312 | 1 or 3 |
| DcFRANCE04 | Type II | 289 | 242 | 207 | 158 | 336 | 169 | 274 | 356 | 215 | 174 | 140 | 113 | 261 | 101 | 310 | 1 or 3 |
| DcFRANCE05 | Type II | 289 | 242 | 207 | 158 | 336 | 169 | 274 | 356 | 243 | 178 | 140 | 111 | 263 | 85 | 310 | 1 or 3 |
| DcFRANCE06 | Type II | 289 | 242 | 207 | 158 | 336 | 169 | 274 | 356 | 221 | 174 | 140 | 115 | 263 | 93 | 312 | 1 or 3 |
| DcFRANCE07 | Type II | 289 | 242 | 207 | 158 | 336 | 169 | 274 | 356 | 213 | 174 | 140 | 109 | 281 | 83 | 310 | 1 or 3 |
| DcFRANCE08 | Type II | 289 | 242 | 207 | 158 | 336 | 169 | 274 | 358 | 213 | 176 | 142 | 115 | 275 | 87 | 312 | 1 or 3 |
| DcFRANCE09 | Type II | 289 | 244 | 207 | 158 | 336 | 169 | 274 | 356 | 213 | 176 | 140 | 113 | 259 | 107 | 310 | 1 or 3 |
| DcFRANCE10 | Type II | 289 | 242 | 207 | 158 | 336 | 169 | 274 | 356 | 221 | 174 | 138 | 111 | 277 | 91 | 312 | 1 or 3 |
| DcFRANCE11 | Type II | 289 | 242 | 207 | 158 | 336 | 169 | 274 | 356 | 221 | 178 | 145 | 115 | 263 | 119 | 308 | 1 or 3 |
| DcFRANCE12 | Type II | 289 | 242 | 207 | 158 | 336 | 169 | 274 | 356 | 219 | 172 | 151 | 117 | 267 | 85 | 310 | 1 or 3 |
| DcFRANCE13 | Type II | 289 | 242 | 207 | 158 | 336 | 169 | 274 | 356 | 227 | 174 | 145 | 109 | 277 | 083 | 310 | 1 or 3 |
| DcFRANCE14 | Type II | 289 | 242 | 207 | 158 | 336 | 169 | 274 | 358 | 215 | 174 | 140 | 111 | 261 | 95 | 310 | 1 or 3 |
| DcFRANCE15 | Type II | 289 | 242 | 207 | 158 | 336 | 169 | 274 | 356 | 227 | 176 | 140 | 111 | 263 | 99 | 310 | 1 or 3 |
| DcFRANCE16 | Type II | 289 | 244 | 207 | 158 | 336 | 169 | 274 | 356 | 213 | 176 | 140 | 113 | 259 | 97 | 310 | 1 or 3 |
| DcFRANCE17 | Type II | 289 | 242 | 207 | 158 | 336 | 169 | 274 | 356 | 215 | 178 | 140 | 109 | 287 | 105 | 312 | 1 or 3 |
| DcFRANCE18 | Type II | 289 | 242 | 207 | 158 | 336 | 169 | 274 | 356 | 243 | 174 | 140 | 111 | 263 | 89 | 312 | 1 or 3 |
| DcFRANCE19 | Type II | 289 | 242 | 207 | 158 | 336 | 169 | 274 | 356 | 213 | 176 | 145 | 111 | 265 | 111 | 320 | 1 or 3 |
| DcFRANCE20 | Type III | 289 | 242 | 205 | 160 | 336 | 165 | 278 | 356 | 211 | 190 | 147 | 111 | 267 | 89 | 312 | 2 |
| DcFRANCE21 | Type II | 289 | 242 | 207 | 158 | 336 | 169 | 274 | 356 | 213 | 184 | 140 | 109 | 285 | 101 | 310 | 1 or 3 |
| DcFRANCE22 | Type II | 289 | 242 | 207 | 158 | 336 | 169 | 274 | 356 | 221 | 174 | 155 | 121 | 259 | 85 | 310 | 1 or 3 |
| DcFRANCE23 | Type II | 289 | 242 | 207 | 158 | 336 | 169 | 274 | 356 | 213 | 188 | 142 | 123 | 259 | 97 | 310 | 1 or 3 |
| DcFRANCE24 | Type II | 289 | 242 | 207 | 158 | 336 | 169 | 274 | 356 | 213 | 174 | 142 | 123 | 257 | 109 | 318 | 1 or 3 |
| DcPANAMA01 | unclassified genotype | 289 | 242 | 205 | 160 | 336 | 165 | 278 | 356 | 221 | 190 | 145 | 111 | 279 | 87 | 314 | 7 |
| DcGABON08 | Africa 1 | 291 | 248 | 205 | 160 | 342 | 165 | 274 | 354 | 223 | 166 | 147 | 111 | 269 | 89 | 306 | 6 |
| DcGABON09 | Africa 1 | 291 | 248 | 205 | 160 | 342 | 165 | 274 | 354 | 231 | 166 | 149 | 111 | 277 | 87 | 306 | 6 |
| DcGABON10 | Africa 3 | 291 | 242 | 207 | 160 | 342 | 165 | 278 | 354 | 225 | 166 | 142 | 111 | 275 | 97 | 310 | 203 |
| DcGABON11 | Africa 3 | 291 | 242 | 207 | 160 | 342 | 165 | 278 | 354 | 223 | 166 | 142 | 111 | 277 | 97 | 310 | 203 |
| DcGABON12 | Africa 3 | 291 | 242 | 207 | 160 | 342 | 165 | 278 | 354 | 229 | 166 | 142 | 111 | 273 | 95 | 310 | 203 |
| DcGABON13 | Africa 3 | 291 | 242 | 207 | 160 | 342 | 165 | 278 | 354 | 225 | 166 | 145 | 111 | 275 | 101 | 310 | 203 |
| DcGABON01 | Type III | 289 | 242 | 205 | 160 | 336 | 165 | 278 | 356 | 211 | 190 | 147 | 111 | 267 | 85 | 312 | 2 |
| DcGABON02 | unclassified genotype | 291 | 242 | 205 | 160 | 336 | 165 | 278 | 356 | 211 | 190 | 147 | 111 | 267 | 85 | 312 | NA |
| DcGABON03 | Type III | 289 | 242 | 205 | 160 | 336 | 165 | 278 | 356 | 213 | 190 | 149 | 111 | 267 | 89 | 312 | 2 |
| DcGABON04 | unclassified genotype | 289 | 242 | 207 | 160 | 336 | 165 | 278 | 356 | 223 | 190 | 147 | 111 | 261 | 103 | 316 | NA |
| DcGABON05 | Type III | 289 | 242 | 205 | 160 | 336 | 165 | 278 | 356 | 213 | 190 | 147 | 111 | 265 | 87 | 314 | 2 |
| DcGABON06 | unclassified genotype | 291 | 242 | 205 | 160 | 336 | 165 | 278 | 356 | 213 | 190 | 145 | 111 | 269 | 89 | 306 | NA |
| DcGABON07 | Type III | 289 | 242 | 205 | 160 | 336 | 165 | 278 | 356 | 213 | 190 | 147 | 111 | 275 | 89 | 312 | 2 |
| DcGUADELOUPE01 | unclassified genotype | 289 | 242 | 207 | 162 | 336 | 169 | 274 | 356 | 237 | 166 | 142 | 111 | 281 | 93 | 312 | NA |
| DcGUADELOUPE02 | Caribbean 2 | 291 | 242 | 205 | 162 | 336 | 165 | 278 | 356 | 211 | 164 | 147 | 109 | 277 | 87 | 312 | 12 or 31 |
| DcUK01 | Type II | 289 | 242 | 207 | 158 | 336 | 169 | 274 | 356 | 227 | 174 | 142 | 121 | 263 | 85 | 310 | 1 or 3 |
| DcUSA03 | Type I | 291 | 248 | 209 | 160 | 342 | 169 | 274 | 358 | 209 | 168 | 145 | 119 | 265 | 87 | 306 | 10 |
| WdFRENCHGUIANA01 | Amazonian | 289 | 242 | 203 | 160 | 336 | 165 | 272 | 356 | 213 | 176 | 140 | 105 | 279 | 87 | 312 | NA |
| WdFRENCHGUIANA02 | Amazonian | 289 | 246 | 209 | 160 | 342 | 173 | 272 | 356 | 239 | 168 | 138 | 113 | 261 | 87 | 312 | NA |
| DcFRENCHGUIANA01 | Caribbean 1 | 291 | 242 | 205 | 162 | 342 | 165 | 278 | 356 | 213 | 164 | 142 | 109 | 277 | 89 | 312 | 13 or 34 |
| DcFRENCHGUIANA02 | Caribbean 2 | 291 | 242 | 205 | 162 | 336 | 165 | 278 | 356 | 213 | 164 | 142 | 109 | 265 | 89 | 312 | 12 or 31 |
| DcFRENCHGUIANA03 | unclassified genotype | 289 | 242 | 205 | 162 | 336 | 165 | 278 | 356 | 219 | 164 | 145 | 105 | 273 | 93 | 312 | NA |
| DcFRENCHGUIANA04 | unclassified genotype | 289 | 242 | 205 | 160 | 336 | 165 | 278 | 356 | 213 | 174 | 140 | 111 | 267 | 87 | 316 | NA |
| WdFRENCHGUIANA03 | unclassified genotype | 289 | 242 | 205 | 160 | 336 | 169 | 274 | 358 | 211 | 172 | 132 | 111 | 257 | 83 | 318 | NA |
| DcFRENCHGUIANA05 | Type III | 289 | 242 | 205 | 160 | 336 | 165 | 278 | 356 | 213 | 190 | 145 | 111 | 269 | 89 | 312 | 2 |
| DcFRENCHGUIANA06 | Type III | 289 | 242 | 205 | 160 | 336 | 165 | 278 | 356 | 213 | 190 | 145 | 111 | 269 | 89 | 312 | 2 |
| WdFRENCHGUIANA09 | Amazonian | 289 | 246 | 203 | 160 | 344 | 167 | 272 | 356 | 229 | 176 | 142 | 113 | 263 | 85 | 312 | 97 |
| WdFRENCHGUIANA10 | Amazonian | 289 | 246 | 203 | 160 | 337 | 165 | 274 | 356 | 209 | 172 | 136 | 111 | 251 | 109 | 310 | 60 |
| WdFRENCHGUIANA11 | Amazonian | 291 | 242 | 203 | 160 | 339 | 165 | 272 | 358 | 221 | 174 | 138 | 107 | 277 | 95 | 312 | 95 |
| WdFRENCHGUIANA12 | Amazonian | 289 | 242 | 203 | 158 | 344 | 171 | 272 | 356 | 209 | 182 | 149 | 121 | 265 | 89 | 317 | 193 |
| WdFRENCHGUIANA13 | Amazonian | 289 | 246 | 203 | 158 | 338 | 167 | 276 | 354 | 213 | 168 | 138 | 111 | 281 | 93 | 318 | NA |
| WdFRENCHGUIANA14 | Amazonian | 291 | 242 | 203 | 160 | 338 | 167 | 274 | 356 | 209 | 188 | 138 | 115 | 263 | 91 | 312 | 60 |
| WdFRENCHGUIANA04 | Amazonian | 291 | 242 | 203 | 160 | 346 | 167 | 272 | 356 | 217 | 170 | 147 | 127 | 257 | 85 | 310 | NA |
| WdFRENCHGUIANA15 | Amazonian | 291 | 246 | 205 | 166 | 334 | 167 | 272 | 356 | 213 | 174 | 151 | 107 | 267 | 87 | 325 | 194 |
| DcFRENCHGUIANA07 | unclassified genotype | 291 | 248 | 209 | 160 | 342 | 165 | 278 | 354 | 209 | 166 | 140 | 113 | 277 | 95 | 304 | NA |
| WdFRENCHGUIANA06 | Amazonian | 291 | 244 | 203 | 162 | 344 | 167 | 272 | 356 | 219 | 174 | 147 | 105 | 277 | 85 | 312 | NA |
| WdFRENCHGUIANA07 | Amazonian | 291 | 242 | 207 | 162 | 336 | 167 | 278 | 358 | 215 | 174 | 138 | 127 | 201 | 85 | 302 | NA |
| WdFRENCHGUIANA08 | Amazonian | 293 | 246 | 203 | 162 | 344 | 171 | 272 | 356 | 211 | 182 | 138 | 105 | 267 | 91 | 302 | NA |
| WdFRENCHGUIANA16 | unclassified genotype | 289 | 242 | 203 | 160 | 336 | 169 | 274 | 358 | 209 | 176 | 142 | 111 | 257 | 107 | 341 | 197 |
| LGE-CUV | unclassified genotype | 287 | 242 | 207 | 158 | 362 | 169 | 274 | 356 | 217 | 164 | 138 | 109 | 273 | 97 | 326 | 17 |
| DcUSA04 | Type III | 289 | 242 | 205 | 160 | 336 | 165 | 278 | 356 | 215 | 190 | 147 | 111 | 267 | 91 | 312 | 133 |
| MAS | unclassified genotype | 291 | 242 | 205 | 162 | 362 | 169 | 272 | 358 | 221 | 166 | 142 | 111 | 332 | 95 | 338 | 17 |

| | | | | | | | | | | | | | | | | | |
|---|---|---|---|---|---|---|---|---|---|---|---|---|---|---|---|---|---|
| **DcUSA05** | **Type II** | **289** | **242** | **207** | **158** | **336** | **169** | **274** | **356** | **215** | **174** | **142** | **111** | **265** | **91** | **310** | **1** |
| **DcMARTINIQUE01** | **Type II** | **289** | **242** | **207** | **158** | **336** | **169** | **274** | **356** | **213** | **174** | **147** | **119** | **271** | **89** | **335** | **1 or 3** |
| **DcMARTINIQUE02** | **Caribbean 2** | **291** | **242** | **205** | **162** | **336** | **165** | **278** | **356** | **213** | **164** | **142** | **111** | **283** | **85** | **312** | **12 or 31** |
| **DcMARTINIQUE03** | **Caribbean 3** | **289** | **242** | **205** | **162** | **336** | **165** | **278** | **356** | **211** | **164** | **142** | **109** | **277** | **85** | **312** | **25 or 141** |
| **DcMARTINIQUE04** | **Caribbean 1** | **291** | **242** | **205** | **162** | **342** | **165** | **278** | **356** | **213** | **164** | **145** | **109** | **279** | **89** | **312** | **13 or 34** |
| **DcMARTINIQUE05** | **unclassified genotype** | **291** | **242** | **205** | **160** | **336** | **165** | **278** | **356** | **211** | **190** | **147** | **111** | **277** | **89** | **312** | **NA** |
| **DcUSA06** | **unclassified genotype** | **291** | **242** | **205** | **160** | **348** | **165** | **278** | **356** | **213** | **190** | **142** | **111** | **261** | **87** | **314** | **8** |
| **DcCHINA01** | **NA** | **-** | **-** | **-** | **-** | **-** | **-** | **-** | **-** | **-** | **-** | **-** | **-** | **-** | **-** | **-** | **9** |
| **DcPORTUGAL01** | **Type II** | **289** | **242** | **207** | **158** | **336** | **169** | **274** | **356** | **219** | **174** | **140** | **127** | **269** | **91** | **308** | **1 or 3** |
| **DcPORTUGAL02** | **Type III** | **289** | **242** | **205** | **160** | **336** | **165** | **278** | **356** | **209** | **192** | **145** | **111** | **269** | **89** | **312** | **2** |
| **DcPORTUGAL03** | **Type II** | **289** | **242** | **207** | **158** | **336** | **169** | **274** | **356** | **223** | **178** | **140** | **111** | **261** | **87** | **312** | **1 or 3** |
| **DcPORTUGAL04** | **Type III** | **289** | **242** | **205** | **160** | **336** | **165** | **278** | **356** | **213** | **190** | **147** | **111** | **263** | **89** | **312** | **2** |
| **DcPORTUGAL05** | **Type II** | **289** | **242** | **207** | **158** | **336** | **169** | **274** | **356** | **213** | **176** | **145** | **113** | **259** | **93** | **310** | **1 or 3** |
| **DcPORTUGAL06** | **Type II** | **289** | **242** | **207** | **158** | **336** | **169** | **274** | **356** | **213** | **176** | **140** | **115** | **273** | **93** | **316** | **1 or 3** |
| **DcPORTUGAL07** | **Type II** | **289** | **244** | **207** | **158** | **336** | **169** | **274** | **356** | **213** | **176** | **140** | **113** | **259** | **99** | **310** | **1 or 3** |
| **DcPORTUGAL08** | **Type II** | **289** | **242** | **207** | **158** | **336** | **169** | **274** | **356** | **233** | **174** | **140** | **123** | **269** | **97** | **308** | **1 or 3** |
| **DcPORTUGAL09** | **Type III** | **289** | **242** | **205** | **160** | **336** | **165** | **278** | **356** | **213** | **190** | **149** | **111** | **267** | **89** | **312** | **2** |
| **DcPORTUGAL10** | **Type I** | **291** | **248** | **209** | **160** | **342** | **169** | **274** | **358** | **209** | **166** | **147** | **119** | **273** | **87** | **306** | **10** |
| **WdUSA04** | **Type 12** | **289** | **242** | **211** | **160** | **336** | **169** | **274** | **362** | **233** | **176** | **151** | **113** | **283** | **99** | **320** | **5** |
| **DcUSA07** | **unclassified genotype** | **289** | **242** | **205** | **160** | **336** | **165** | **278** | **356** | **213** | **190** | **147** | **111** | **267** | **89** | **314** | **72** |
| **WdFRENCHGUIANA17** | **Amazonian** | **289** | **242** | **205** | **170** | **360** | **167** | **274** | **356** | **223** | **190** | **142** | **109** | **259** | **85** | **312** | **98** |
| **DcSENEGAL01** | **Type II** | **289** | **242** | **207** | **158** | **336** | **169** | **274** | **356** | **227** | **176** | **140** | **119** | **261** | **101** | **308** | **1 or 3** |
| **DcSENEGAL02** | **Type II** | **289** | **242** | **207** | **158** | **336** | **169** | **274** | **356** | **227** | **174** | **140** | **111** | **269** | **113** | **308** | **1 or 3** |
| **DcSENEGAL03** | **Type II** | **289** | **242** | **207** | **158** | **336** | **169** | **274** | **356** | **223** | **176** | **142** | **113** | **279** | **85** | **310** | **1 or 3** |
| **DcSENEGAL04** | **Africa 4** | **293** | **242** | **203** | **156** | **336** | **165** | **274** | **354** | **217** | **174** | **130** | **109** | **281** | **101** | **306** | **137** |
| **DcSENEGAL05** | **Africa 1** | **291** | **248** | **205** | **160** | **342** | **165** | **274** | **354** | **231** | **166** | **144** | **111** | **279** | **89** | **308** | **6** |
| **DcSENEGAL06** | **Type II** | **289** | **242** | **207** | **158** | **336** | **169** | **274** | **356** | **227** | **174** | **140** | **111** | **273** | **101** | **308** | **1 or 3** |
| **DcSENEGAL07** | **Type II** | **289** | **242** | **207** | **158** | **336** | **169** | **274** | **356** | **227** | **174** | **140** | **111** | **283** | **127** | **308** | **1 or 3** |
| **DcSENEGAL08** | **Type II** | **289** | **242** | **207** | **158** | **336** | **169** | **274** | **356** | **227** | **174** | **140** | **111** | **269** | **107** | **308** | **1 or 3** |
| **DcSENEGAL09** | **Type III** | **289** | **242** | **205** | **160** | **336** | **165** | **278** | **356** | **209** | **188** | **151** | **111** | **275** | **91** | **312** | **2** |
| **DcSENEGAL10** | **Africa 1** | **291** | **248** | **205** | **160** | **342** | **165** | **274** | **354** | **235** | **166** | **145** | **111** | **269** | **89** | **306** | **6** |
| **DcSENEGAL11** | **unclassified genotype** | **289** | **248** | **205** | **160** | **336** | **165** | **278** | **354** | **213** | **190** | **149** | **111** | **265** | **89** | **306** | **NA** |
| **DcSENEGAL12** | **Type II** | **289** | **242** | **207** | **158** | **336** | **169** | **274** | **356** | **229** | **174** | **140** | **119** | **273** | **121** | **308** | **1 or 3** |
| **DcSENEGAL13** | **Type II** | **289** | **242** | **207** | **158** | **336** | **169** | **274** | **356** | **231** | **174** | **140** | **111** | **275** | **105** | **308** | **1 or 3** |
| **DcSENEGAL14** | **Africa 4** | **291** | **242** | **203** | **156** | **336** | **165** | **274** | **354** | **223** | **174** | **130** | **109** | **312** | **99** | **310** | **20** |
| **DcSENEGAL15** | **Type III** | **289** | **242** | **205** | **160** | **336** | **165** | **278** | **356** | **213** | **190** | **149** | **111** | **267** | **91** | **312** | **2** |
| **DcSENEGAL16** | **Type II** | **289** | **242** | **207** | **158** | **336** | **169** | **274** | **356** | **217** | **174** | **140** | **113** | **259** | **89** | **310** | **1 or 3** |
| **DcSENEGAL17** | **Type II** | **289** | **244** | **207** | **158** | **336** | **169** | **274** | **356** | **215** | **176** | **140** | **113** | **259** | **97** | **312** | **1 or 3** |
| **DcSENEGAL18** | **Type II** | **289** | **244** | **207** | **158** | **336** | **169** | **274** | **356** | **213** | **176** | **140** | **113** | **275** | **99** | **310** | **1 or 3** |
| **DcSENEGAL19** | **Type II** | **289** | **242** | **207** | **158** | **336** | **169** | **274** | **356** | **223** | **174** | **140** | **119** | **261** | **103** | **308** | **1 or 3** |
| **DcSENEGAL20** | **Type III** | **289** | **242** | **205** | **160** | **336** | **165** | **278** | **356** | **209** | **192** | **153** | **111** | **269** | **89** | **312** | **2** |
| **DcSENEGAL21** | **Type II** | **289** | **244** | **207** | **158** | **336** | **169** | **274** | **356** | **213** | **176** | **140** | **113** | **259** | **99** | **310** | **1 or 3** |
| **DcSENEGAL22** | **Africa 1** | **291** | **248** | **205** | **160** | **342** | **165** | **274** | **354** | **225** | **166** | **147** | **111** | **269** | **89** | **304** | **6** |
| **DcSENEGAL23** | **Africa 4** | **291** | **242** | **203** | **156** | **336** | **165** | **274** | **354** | **223** | **174** | **130** | **109** | **303** | **101** | **310** | **20** |
| **DcSENEGAL24** | **Type II** | **289** | **242** | **207** | **158** | **336** | **169** | **274** | **356** | **229** | **174** | **140** | **111** | **275** | **113** | **308** | **1 or 3** |
| **DcSENEGAL25** | **Africa 1** | **291** | **248** | **205** | **160** | **342** | **165** | **274** | **354** | **231** | **166** | **147** | **111** | **271** | **89** | **306** | **6** |
| **DcSENEGAL26** | **Type III** | **289** | **242** | **205** | **160** | **336** | **165** | **278** | **356** | **209** | **190** | **149** | **111** | **273** | **89** | **312** | **2** |
| **DcUSA08** | **unclassified genotype** | **289** | **242** | **205** | **158** | **336** | **165** | **278** | **356** | **225** | **174** | **142** | **111** | **259** | **89** | **312** | **139** |
| **DcSERBIA01** | **Type II** | **289** | **242** | **207** | **158** | **336** | **169** | **274** | **356** | **211** | **176** | **142** | **109** | **293** | **89** | **310** | **1 or 3** |
| **DcBRAZIL02** | **unclassified genotype** | **289** | **242** | **205** | **160** | **342** | **165** | **278** | **358** | **233** | **164** | **147** | **111** | **316** | **89** | **308** | **11** |
| **DcBRAZIL03** | **unclassified genotype** | **291** | **242** | **205** | **160** | **362** | **165** | **278** | **356** | **237** | **174** | **140** | **111** | **265** | **89** | **314** | **19** |
| **DcBRAZIL04** | **unclassified genotype** | **291** | **242** | **205** | **160** | **362** | **165** | **278** | **354** | **227** | **174** | **140** | **111** | **269** | **89** | **308** | **42** |
| **DcBRAZIL05** | **unclassified genotype** | **291** | **242** | **207** | **160** | **360** | **165** | **278** | **356** | **229** | **174** | **140** | **105** | **263** | **91** | **314** | **21** |
| **DcBRAZIL06** | **unclassified genotype** | **289** | **242** | **205** | **162** | **344** | **165** | **278** | **358** | **225** | **164** | **142** | **111** | **263** | **105** | **312** | **14** |
| **DcBRAZIL07** | **unclassified genotype** | **291** | **242** | **207** | **160** | **338** | **169** | **272** | **358** | **229** | **164** | **142** | **111** | **263** | **89** | **308** | **119** |
| **DcBRAZIL08** | **unclassified genotype** | **291** | **242** | **207** | **160** | **338** | **169** | **272** | **358** | **229** | **164** | **142** | **111** | **263** | **89** | **308** | **47** |
| **DcBRAZIL09** | **unclassified genotype** | **291** | **248** | **205** | **160** | **362** | **165** | **278** | **354** | **229** | **174** | **140** | **111** | **271** | **89** | **308** | **80** |
| **DcBRAZIL10** | **unclassified genotype** | **289** | **242** | **205** | **160** | **348** | **165** | **278** | **356** | **213** | **190** | **142** | **111** | **263** | **113** | **312** | **8** |
| **DcBRAZIL11** | **unclassified genotype** | **291** | **248** | **205** | **160** | **338** | **169** | **272** | **356** | **245** | **164** | **136** | **111** | **316** | **87** | **314** | **104** |
| **DcBRAZIL12** | **unclassified genotype** | **291** | **242** | **205** | **162** | **342** | **165** | **278** | **356** | **231** | **166** | **153** | **111** | **265** | **91** | **344** | **34** |
| **DcBRAZIL13** | **unclassified genotype** | **289** | **242** | **207** | **160** | **338** | **165** | **278** | **356** | **225** | **190** | **136** | **105** | **263** | **97** | **310** | **111** |
| **DcBRAZIL14** | **unclassified genotype** | **289** | **248** | **205** | **160** | **342** | **165** | **274** | **354** | **231** | **164** | **147** | **111** | **271** | **89** | **308** | **85** |
| **DcBRAZIL15** | **unclassified genotype** | **291** | **248** | **209** | **160** | **344** | **165** | **278** | **356** | **209** | **166** | **142** | **113** | **281** | **95** | **306** | **77** |
| **DcCOSTARICA1** | **unclassified genotype** | **291** | **248** | **209** | **160** | **348** | **NA** | **276** | **358** | **209** | **166** | **142** | **125** | **259** | **89** | **306** | **91** |
| **DcCOSTARICA2** | **unclassified genotype** | **-** | **-** | **-** | **-** | **-** | **-** | **-** | **-** | **-** | **-** | **-** | **-** | **-** | **-** | **-** | **43** |
| **DcGUYANA01** | **Caribbean 2** | **291** | **242** | **205** | **162** | **336** | **165** | **278** | **NA** | **213** | **164** | **142** | **109** | **265** | **103** | **312** | **12** |
| **DcCOLUMBIA01** | **unclassified genotype** | **291** | **242** | **205** | **160** | **336** | **165** | **276** | **356** | **223** | **166** | **142** | **121** | **279** | **87** | **304** | **61** |
| **DcCOLUMBIA02** | **unclassified genotype** | **291** | **244** | **209** | **160** | **336** | **165** | **278** | **358** | **213** | **166** | **147** | **119** | **263** | **91** | **306** | **38** |
| **TgH20005** | **unclassified genotype** | **287** | **242** | **207** | **160** | **354** | **169** | **274** | **356** | **221** | **168** | **147** | **109** | **289** | **87** | **324** | **17** |
| **TgH26044** | **unclassified genotype** | **287** | **242** | **207** | **158** | **354** | **169** | **274** | **356** | **221** | **168** | **155** | **109** | **281** | **103** | **323** | **NA** |
| **DcCOSTARICA3** | **unclassified genotype** | **291** | **248** | **205** | **160** | **364** | **165** | **274** | **356** | **209** | **192** | **140** | **115** | **263** | **97** | **304** | **52** |
| **DcUSA09** | **unclassified genotype** | **291** | **248** | **209** | **160** | **336** | **165** | **278** | **354** | **213** | **166** | **147** | **111** | **277** | **87** | **304** | **73** |
| **DcTUNISIA01** | **Type II** | **289** | **242** | **207** | **158** | **336** | **169** | **274** | **356** | **225** | **176** | **140** | **123** | **259** | **95** | **310** | **1 or 3** |
| **DcTUNISIA02** | **Type III** | **289** | **242** | **205** | **160** | **336** | **165** | **278** | **356** | **213** | **190** | **147** | **111** | **265** | **89** | **312** | **2** |
| **DcTUNISIA03** | **Type II** | **289** | **242** | **207** | **158** | **336** | **169** | **274** | **356** | **215** | **182** | **140** | **109** | **279** | **111** | **310** | **1 or 3** |
| **DcTURKEY01** | **Africa 1** | **291** | **248** | **205** | **160** | **342** | **165** | **274** | **354** | **227** | **166** | **147** | **111** | **297** | **91** | **310** | **6** |
| **DcTURKEY02** | **Africa 1** | **291** | **248** | **205** | **160** | **342** | **165** | **274** | **354** | **227** | **166** | **149** | **111** | **287291** | **91** | **310** | **6** |
| **DcUSA01** | **Type I** | **291** | **248** | **209** | **160** | **342** | **169** | **274** | **358** | **209** | **168** | **145** | **119** | **265** | **87** | **306** | **10** |
| **WdFRENCHGUIANA05** | **Amazonian** | **291** | **242** | **203** | **162** | **344** | **167** | **276** | **356** | **217** | **170** | **142** | **113** | **277** | **91** | **308** | **60** |
| **DcUSA10** | **Type III** | **289** | **242** | **205** | **160** | **336** | **165** | **278** | **356** | **213** | **188** | **153** | **111** | **267** | **89** | **312** | **2** |

NA: Not available

**Supplementary Table 3. Estimates of the time (in years) to the most recent common ancestor (TMRCA) between New World strains and their closest relative from the Old World belonging to the same clonal lineage**

| | New World strain | Old World strain | Lineage | Lower estimate | Upper estimate |
|---|---|---|---|---|---|
| 1 | DcUSA03 | DcPORTUGAL10 | typeI | 651 | 2457 |
| 2 | DcUSA05 | DcFRANCE14 | typeII | 4725 | 17832 |
| 3 | DcUSA10 | DcTUNISIA02 | typeIII | 286 | 1080 |
| 4 | DcFRENCHGUIANA05 | DcPORTUGAL02 | typeIII | 408 | 1539 |

#### Supplementary Table 4. Estimates of the time (in years) to the most recent common ancestor (TMRCA) between full chromosomes of New World hybrid strains and their closest relative from the Old World having the same respective chromosomal ancestry

| ID | chr01a | chr01b | chr02 | chr03 | chr04 | chr05 | chr06 |
|---|---|---|---|---|---|---|---|
| DcBRAZIL01 | | | | | | | |
| DcBRAZIL02 | | | | | | | |
| DcBRAZIL03 | | | | | | | |
| DcBRAZIL04 | | | | | | 1 DcSENEGAL22: 361-1,364 | |
| DcBRAZIL05 | | | | | 1 DcPORTUGAL02*: 317-1,197 | | |
| DcBRAZIL06 | | | | | 2 DcSENEGAL11*: 634-2,394 | | |
| DcBRAZIL07 | | | | | 3 DcPORTUGAL04*: 396-1,496 | | |
| DcBRAZIL08 | | | | | 4 | | |
| DcBRAZIL09 | | | 1 DcSENEGAL05: 322-1,214 | | 5 | 2 DcSENEGAL22: 361-1,364 | |
| DcBRAZIL10 | | | | | 6 | | |
| DcBRAZIL11 | | | | | 4 DcPORTUGAL02*: 396-1,496 | | |
| DcBRAZIL12 | 1 DcFRANCE01: 2,014-7,601 | | | | 5 DcGABON07*: 793-2,992 | | |
| DcBRAZIL13 | | | | | 6 DcSENEGAL11*: 634-2,394 | | |
| DcBRAZIL14 | | | 2 DcSENEGAL25: 643-2,428 | | 7 DcBENIN01: 79-299 | | |
| DcBRAZIL15 | 2 DcPORTUGAL10: 3,491-13,175 | | | 1 DcPORTUGAL10*: 997-3,761 | 11 | 3 DcCAMEROON01: 663-2,501 | |
| DcCOSTARICA1 | | 1 DcPORTUGAL07: 5,362-20,238 | 3 DcSENEGAL22: 161-607 | 2 DcPORTUGAL10*: 613-2,315 | 12 | 4 DcCAMEROON01: 663-2,501 | 1 DcPORTUGAL10: 610-2,302 |
| DcCOSTARICA2 | 3 DcPORTUGAL02*: 268-1,013 | | | 3 DcPORTUGAL10*: 843-3,183 | 13 | | |
| DcCOLUMBIA01 | | | | | 14 | | |
| DcCOLUMBIA02 | | | | 4 DcPORTUGAL09: 307-1,157 | 8 DcPORTUGAL02: 159-598 | 5 DcCAMEROON01: 603-2,274 | |
| DcCOSTARICA3 | | | | | 16 | | |
| DcFRENCHGUIANA01 | | | | 5 DcPORTUGAL09: 1,686-6,365 | 9 DcSENEGAL11*: 317-1,197 | 6 DcPORTUGAL09: 542-2,047 | 2 DcPORTUGAL10: 793-2,992 |
| DcFRENCHGUIANA02 | | | | | 18 | | |
| DcFRENCHGUIANA03 | | | | 6 DcPORTUGAL09: 383-1,447 | 10 DcGABON07*: 317-1,197 | | |
| DcFRENCHGUIANA04 | | | | | 11 DcPORTUGAL02: 79-299 | | |
| DcFRENCHGUIANA | 4 DcPORTUGAL10: 3,088-11,655 | | | 7 SENEGAL22*: 1,150-4,340 | 21 | 7 DcSENEGAL10: 783-2,956 | 3 DcBENIN03: 549-2,071 |
| DcGUADELOUPE01 | 5 DcFRANCE13: 940-3,547 | 2 DcFRANCE02: 699-2,640 | 4 DcFRANCE14: 402-1,517 | 8 DcSPAIN01: 6,593-24,882 | 22 | 8 DcSPAIN01: 3,133-11,825 | 4 DcUK01: 793-2,992 |
| DcGUADELOUPE02 | | | | 9 DcPORTUGAL09: 1,610-6,076 | 12 DcGABON07*: 555-2,094 | 9 DcPORTUGAL09: 241-910 | |
| DcGUYANA01 | | | 5 DcPORTUGAL04*: 161-607 | 10 DcPORTUGAL09: 307-1,157 | 13 DcPORTUGAL02*: 159-598 | 10 DcPORTUGAL09: 542-2,047 | |
| DcMARTINIQUE01 | 6 DcGABON07: 268-1,013 | | | | 14 DcPORTUGAL10: 1,348-5,087 | 11 DcPORTUGAL02: 301-1,137 | |
| DcMARTINIQUE02 | | | 6 DcFRANCE20: 241-910 | 11 DcSENEGAL15: 307-1,157 | 15 DcGABON07*: 238-898 | 12 DcPORTUGAL02: 301-1,137 | 5 DcGABON10: 549-2,071 |
| DcMARTINIQUE03 | | | | 12 DcPORTUGAL09: 1,763-6,654 | 16 DcGABON07*: 396-1,496 | 13 DcPORTUGAL02: 301-1,137 | 6 DcPORTUGAL10: 731-2,762 |
| DcMARTINIQUE04 | | | | | | | |
| DcMARTINIQUE05 | 7 DcBENIN03*: 2,685-10,134 | 3 DcFRANCE20: 117-440 | | 13 DcPORTUGAL09: 383-1,447 | 17 DcPORTUGAL04*: 475-1,795 | 14 DcPORTUGAL09: 361-1,364 | 7 DcGABON10: 549-2,071 |
| DcPANAMA01 | | 4 DcPORTUGAL09: 350-1,320 | 7 DcPORTUGAL04*: 322-1,214 | 14 DcPORTUGAL09: 153-579 | 18 DcSENEGAL11*: 317-1,197 | | |
| DcURUGUAY01 | | | | | | | |
| DcUSA02 | 8 DcPORTUGAL10: 1,343-5,067 | | | | | | |
| DcUSA04 | 9 DcPORTUGAL02*: 134-507 | | 8 DcSENEGAL15*: 322-1,214 | 15 DcPORTUGAL09: 77-289 | 19 DcPORTUGAL02*: 159-598 | 15 DcPORTUGAL02: 120-455 | |
| DcUSA06 | | | | | 20 DcGABON07*: 555-2,094 | | |
| DcUSA07 | 10 DcSENEGAL20: 268-1,013 | 5 DcPORTUGAL09: 117-440 | | 16 DcPORTUGAL09: 77-289 | | 16 DcPORTUGAL09: 241-910 | |
| DcUSA08 | 11 DcUK01: 1,074-4,054 | 6 DcFRANCE02: 699-2,640 | 9 DcFRANCE14: 723-2,731 | 17 DcFRANCE14: 383-1,447 | 21 DcSENEGAL11*: 396-1,496 | 17 DcPORTUGAL02: 181-682 | 8 DcTUNISIA: 183-691 |
| DcUSA09 | 12 DcTUNISIA02: 402-1,520 | 7 DcPORTUGAL09: 350-1,320 | 10 DcCAMEROON01: 322-1,214 | 18 DcPORTUGAL09: 77-289 | 22 DcGABON07*: 238-898 | 18 DcPORTUGAL09: 361-1,364 | 9 DcGABON12: 731-2,762 |
| WdUSA01 | | | | 19 DcTUNISIA01: 47,068-177,645 | | | |
| WdUSA02 | | 8 DcFRANCE02: 233-880 | 11 DcTUNISIA01: 29,512-111,386 | 20 DcFRANCE14: 613-2,315 | | | 10 DcFRANCE21: 2,805-10,588 |
| WdUSA03 | 13 DcPORTUGAL02*: 268-1,013 | 9 DcFRANCE02: 233-880 | 12 DcFRANCE14: 643-2,428 | 21 DcFRANCE14: 920-3,472 | 23 DcFRANCE14: 2,933-11,071 | 19 DcPORTUGAL09: 361-1,364 | |
| WdUSA04 | | | 13 DcTUNISIA02: 39,082-147,503 | 22 DcTUNISIA01: 36,556-138,008 | | | |

| ID | chr07a | chr08 | chr09 | chr10 | chr11 | chr12 |
|---|---|---|---|---|---|---|
| DcBRAZIL01 | | | | | | |
| DcBRAZIL02 | | | | | | |
| DcBRAZIL03 | | | | | | |
| DcBRAZIL04 | | | | | | |
| DcBRAZIL05 | | | | | | |
| DcBRAZIL06 | | | | | | |
| DcBRAZIL07 | | | | | | |
| DcBRAZIL08 | | | | | | |
| DcBRAZIL09 | | 1 DcCAMEROON01: 344-1,300 | 1 DcBENIN02: 399-1,505 | | | |
| DcBRAZIL10 | | | | | | |
| DcBRAZIL11 | | | | | | |
| DcBRAZIL12 | | | | | | |
| DcBRAZIL13 | | | | | | |
| DcBRAZIL14 | | | | | | 1 DcSENEGAL05: 408-1,541 |
| DcBRAZIL15 | | | | | | |
| DcCOSTARICA1 | | | | | | |
| DcCOSTARICA2 | | | | | | |
| DcCOLUMBIA01 | | | | | | |
| DcCOLUMBIA02 | | | | | | |
| DcCOSTARICA3 | | | | | | |
| DcFRENCHGUIANA01 | | | | | | |
| DcFRENCHGUIANA02 | | | | | | |
| DcFRENCHGUIANA03 | | | 2 DcGABON07: 632-2,384 | | | |
| DcFRENCHGUIANA04 | | | | | | |
| DcFRENCHGUIANA07 | | | | | | |
| DcGUADELOUPE01 | | 2 DcFRANCE14: 1,580-5,963 | | | 1 DcFRANCE14: 570-2150 | 2 DcFRANCE02: 534-2,015 |
| DcGUADELOUPE02 | | | | | | |
| DcGUYANA01 | | | | | | |
| DcMARTINIQUE01 | | | | | | |
| DcMARTINIQUE02 | | | | | | |
| DcMARTINIQUE03 | | | | | | |
| DcMARTINIQUE04 | | | | | | |
| DcMARTINIQUE05 | | | | 1 DcSENEGAL15: 328-1,238 | 2 DcPORTUGAL04*: 427-1612 | 3 DcGABON01:188-711 |
| DcPANAMA01 | | | | | 3 DcPORTUGAL04*: 284-1075 | |
| DcURUGUAY01 | | | | | | |
| DcUSA02 | | | | | | |
| DcUSA04 | 1 DcGABON3: 358-1,351 | 3 DcPORTUGAL04: 344-1,300 | 3 DcGABON01: 598-2,258 | | 4 DcPORTUGAL04: 356-1344 | 4 DcGABON01: 157-593 |
| DcUSA06 | | | | | | |
| DcUSA07 | 2 DcGABON3: 409-1,543 | | | | 5 DcPORTUGAL04*: 427-1612 | |
| DcUSA08 | | | 4 DcGABON01: 166-627 | 2 DcSENEGAL15: 209-788 | 6 DcSENEGAL15*: 249-940 | 5 DcGABON01: 283-1,067 |
| DcUSA09 | | | | | | |
| WdUSA01 | | | | | | |
| WdUSA02 | | | 5 DcUK01: 3,623-13,674 | | 7 DcFRANCE14: 714-2687 | |
| WdUSA03 | 3 DcGABON03: 153-579 | 4 DcFRANCE14: 6,441-24,311 | | 3 DcPORTUGAL08: 6,443-24,317 | 8 DcSENEGAL15*: 392-1478 | |
| WdUSA04 | | | | | 9 DcSENEGAL20: 47985-181106 | 6 DcTUNISIA01: 37,780-142,590 |

Each cell indicates the Old World strain that is the most closely related to the New World hybrid strain at this chromosome (lowest TMRCA). The colour indicates the lineage to which belong  to this Old World strain: type I in red, type II in green, type III in blue, Africa 1 in purple and Africa 3 in grey. The stars indicate that another Old World strain from another continent has an equal TMRCA at this chromosome. Empty cells indicate that the New World hybrid strain has a mixed ancestry at this chromosome.

**Supplementary Table 5. Candidate missense variants involved in the process of adaptation to the domestic environment**

| chromosome | position | REF | ALT | gene | gene ID | gene alternative ID | start | end |
|---|---|---|---|---|---|---|---|---|
| chromosome 1a | 1373109 | G | A | exonuclease | TGRH88_020120 | TGME49_295060 | 1373043 | 1376325 |
| chromosome 1a | 1391757 | T | G | hypothetical protein | TGRH88_020140 | TGME49_295080 | 1391288 | 1393202 |
| chromosome 1a | 1392832 | A | G | | | | | |
| chromosome 1a | 1396686 | G | A | hypothetical protein | TGRH88_020150 | TGME49_295090 | 1396511 | 1400356 |
| chromosome 1a | 1397103 | C | T | | | | | |
| chromosome 1a | 1398069 | A | C | | | | | |
| chromosome 1a | 1398289 | C | T | | | | | |
| chromosome 1a | 1398396 | G | A | | | | | |
| chromosome 1a | 1400034 | G | C | | | | | |
| chromosome 1a | 1403822 | T | C | hypothetical protein | TGRH88_020160 | TGME49_295100 | 1403656 | 1409958 |
| chromosome 1a | 1404032 | C | A | | | | | |
| chromosome 1a | 1404037 | T | G | | | | | |
| chromosome 1a | 1405492 | C | T | | | | | |
| chromosome 1a | 1406320 | G | A | | | | | |
| chromosome 1a | 1406330 | T | C | | | | | |
| chromosome 1a | 1407316 | G | A | | | | | |
| chromosome 1a | 1407730 | G | A | | | | | |
| chromosome 1a | 1408511 | T | A | | | | | |
| chromosome 1a | 1408824 | A | G | | | | | |
| chromosome 1a | 1409751 | A | C | | | | | |
| chromosome 1a | 1423144 | TG | TC | rhoptry protein ROP4 | TGRH88_020200 | TGME49_295125 | 1422225 | 1423961 |
| chromosome 1a | 1423857 | CTGTC | CTGTA | | | | | |
| chromosome 1a | 1433501 | C | T | hypothetical protein | TGRH88_020230 | TGME49_296015 | 1433000 | 1434202 |
| chromosome 1a | 1438724 | G | A | phosphatidylinositol 3- and 4-kinase | TGRH88_020240 | TGME49_296010 | 1435694 | 1445711 |
| chromosome 1a | 1448598 | C | T | hypothetical protein | TGRH88_020250 | TGME49_296000 | 1448137 | 1448721 |
| chromosome 1a | 1459039 | A | C | putative ubiquitin conjugating enzyme E2 | TGRH88_020270 | TGME49_295990 | 1458798 | 1462319 |
| chromosome 1a | 1459408 | C | G | | | | | |
| chromosome 1a | 1459591 | A | C | | | | | |
| chromosome 1a | 1460091 | A | G | | | | | |
| chromosome 1a | 1466358 | G | C | hypothetical protein | TGRH88_020280 | TGME49_295970 | 1463187 | 1469120 |
| chromosome 1a | 1466687 | G | C | | | | | |
| chromosome 1a | 1466926 | G | C | | | | | |
| chromosome 1a | 1466973 | G | T | | | | | |
| chromosome 1a | 1468777 | G | A | | | | | |
| chromosome 1a | 1469051 | C | G | | | | | |
| chromosome 1a | 1473271 | A | G | KRUF family protein | TGRH88_020300 | TGME49_295950 | 1473240 | 1477974 |
| chromosome 1a | 1473310 | T | G | | | | | |
| chromosome 1a | 1473407 | C | T | | | | | |
| chromosome 1a | 1473499 | C | G | | | | | |
| chromosome 1a | 1473860 | T | C | | | | | |
| chromosome 1a | 1473991 | T | C | | | | | |
| chromosome 1a | 1474721 | T | C | | | | | |
| chromosome 1a | 1474762 | T | C | | | | | |
| chromosome 1a | 1477602 | C | G | | | | | |
| chromosome 1a | 1479692 | T | G | | | | | |
| chromosome 1a | 1481161 | G | C | KRUF family protein | TGRH88_020320 | TGME49_295935 | 1480505 | 1485327 |
| chromosome 1a | 1482760 | A | C | | | | | |
| chromosome 1a | 1484150 | A | T | | | | | |
| chromosome 1a | 1484406 | C | T | | | | | |
| chromosome 1a | 1484603 | T | C | | | | | |
| chromosome 1a | 1493248 | C | T | hypothetical protein | TGRH88_020330 | TGME49_295920 | 1491437 | 1496860 |
| chromosome 1a | 1493913 | C | T | | | | | |
| chromosome 1a | 1495970 | T | G | | | | | |
| chromosome 1a | 1496377 | G | T | | | | | |
| chromosome 1a | 1496409 | G | A | | | | | |
| chromosome 1a | 1500274 | A | C | ATP-dependent Clp protease adaptor protein ClpS protein | TGRH88_020340 | TGME49_295910 | 1499801 | 1503215 |
| chromosome 1a | 1500312 | C | G | | | | | |
| chromosome 1a | 1501768 | G | A | | | | | |

#### Supplementary Table 6. Candidate genes involved in the process of adaptation to the domestic environment

| chromosome | gene | gene ID | gene alternative ID | start | end | stage specificity |
|---|---|---|---|---|---|---|
| chromosome 1a | HEAT repeat-containing protein | TGRH88_020100 | TGME49_295040 | 1350985 | 1362413 | none |
| chromosome 1a | tRNA ligase class II core domain (G, H, P, S and T) domain-containing protein | TGRH88_020110 | TGME49_295050 | 1366210 | 1370652 | none |
| chromosome 1a | exonuclease | TGRH88_020120 | TGME49_295060 | 1373043 | 1376325 | none |
| chromosome 1a | helicase associated domain (ha2) protein | TGRH88_020130 | TGME49_295070 | 1377505 | 1389595 | none |
| chromosome 1a | hypothetical protein | TGRH88_020140 | TGME49_295080 | 1391288 | 1393202 | none |
| chromosome 1a | hypothetical protein | TGRH88_020150 | TGME49_295090 | 1396511 | 1400356 | none |
| chromosome 1a | hypothetical protein | TGRH88_020160 | TGME49_295100 | 1403656 | 1409958 | bradyzoite |
| chromosome 1a | putative rhoptry protein | TGRH88_020170 | TGME49_295105 | 1411822 | 1412130 | none |
| chromosome 1a | rhoptry protein ROP7 | TGRH88_020180 | TGME49_295110 | 1414348 | 1415373 | none |
| chromosome 1a | rhoptry protein ROP7 | TGRH88_020190 | TGME49_295125 | 1418290 | 1420017 | none |
| chromosome 1a | rhoptry protein ROP4 | TGRH88_020200 | TGME49_295130 | 1422225 | 1423961 | none |
| chromosome 1a | 1-phosphatidylinositol 4-kinase | TGRH88_020210 | TGME49_295140 | 1427938 | 1428662 | bradyzoite |
| chromosome 1a | hypothetical protein | TGRH88_020220 | TGME49_296020 | 1430324 | 1430797 | sporozoite |
| chromosome 1a | hypothetical protein | TGRH88_020230 | TGME49_296015 | 1433000 | 1434202 | none |
| chromosome 1a | phosphatidylinositol 3- and 4-kinase | TGRH88_020240 | TGME49_296010 | 1435694 | 1445711 | none |
| chromosome 1a | hypothetical protein | TGRH88_020250 | TGME49_296000 | 1448137 | 1448721 | none |
| chromosome 1a | hypothetical protein | TGRH88_020260 | TGME49_295995 | 1452548 | 1453991 | enteroepithelial stages (EES) |
| chromosome 1a | putative ubiquitin conjugating enzyme E2 | TGRH88_020270 | TGME49_295990 | 1458798 | 1462319 | none |
| chromosome 1a | hypothetical protein | TGRH88_020280 | TGME49_295980 | 1463187 | 1469120 | none |
| chromosome 1a | hypothetical protein | TGRH88_020290 | TGME49_295970 | 1469141 | 1469990 | none |
| chromosome 1a | KRUF family protein | TGRH88_020300 | TGME49_295950 | 1473240 | 1477974 | none |
| chromosome 1a | hypothetical protein | TGRH88_020310 | TGME49_295945 | 1478954 | 1479757 | bradyzoite |
| chromosome 1a | KRUF family protein | TGRH88_020320 | TGME49_295935 | 1480505 | 1485327 | none |
| chromosome 1a | hypothetical protein | TGRH88_020330 | TGME49_295920 | 1491437 | 1496860 | enteroepithelial stages (EES) |
| chromosome 1a | ATP-dependent Clp protease adaptor protein ClpS protein | TGRH88_020340 | TGME49_295910 | 1499801 | 1503215 | none |

**Supplementary Table 7. Amino acid subtitutions on the top candidate gene for cat-adaptation TGRH88_020330 (TGME49_295920)**

| ID | M | S | K | N | Y | D | Q | S | N | L |
|---|---|---|---|---|---|---|---|---|---|---|
| WdUSA02 | | | | G | H | G | | R | H | S |
| WdUSA03 | | | | G | H | G | | R | H | S |
| DcBENIN04 | | | | G | H | G | | R | H | S |
| DcBRAZIL01 | | | | G | H | G | | R | H | S |
| DcUSA02 | | | | G | H | G | | R | H | S |
| DcSPAIN01 | | | | G | H | G | | R | H | S |
| DcPANAMA01 | | | | G | H | G | | R | H | S |
| DcGABON08 | | | | G | H | G | | R | H | S |
| DcGABON10 | | | | G | H | G | | R | H | S |
| DcGABON04 | | | | G | H | G | | R | H | S |
| DcGABON06 | | | | G | H | G | | R | H | S |
| DcGUADELOUPE01 | | | | G | H | G | | R | H | S |
| DcGUADELOUPE02 | | | | G | H | G | | R | H | S |
| DcFRENCHGUIANA01 | | | | G | H | G | | R | H | S |
| DcFRENCHGUIANA03 | | | | G | H | G | | R | H | S |
| DcFRENCHGUIANA04 | | | | G | H | G | | R | H | S |
| DcFRENCHGUIANA07 | | | | G | H | G | | R | H | S |
| DcUSA04 | | | | G | H | G | | R | H | S |
| DcMARTINIQUE01 | | | | G | H | G | | R | H | S |
| DcMARTINIQUE02 | | | | G | H | G | | R | H | S |
| DcMARTINIQUE03 | | | | G | H | G | | R | H | S |
| DcMARTINIQUE05 | | | | G | H | G | | R | H | S |
| DcUSA06 | | | | G | H | G | | R | H | S |
| DcCHINA01 | | | | G | H | G | | R | H | S |
| DcPORTUGAL09 | | | | G | H | G | | R | H | S |
| DcPORTUGAL10 | | | | G | H | G | | R | H | S |
| DcUSA07 | | | | G | H | G | | R | H | S |
| DcSENEGAL04 | | | Q | G | | G | | R | H | S |
| DcSENEGAL11 | | | | G | H | G | | R | H | S |
| DcSENEGAL23 | | | Q | G | | G | | R | H | S |
| DcUSA08 | | | | G | H | G | | R | H | S |
| DcBRAZIL03 | | | | G | H | G | | R | H | S |
| DcBRAZIL04 | | | | G | H | G | | R | H | S |
| DcBRAZIL05 | | | | G | H | G | | R | H | S |
| DcBRAZIL06 | | | | G | H | G | | R | H | S |
| DcBRAZIL08 | | | | G | H | G | | R | H | S |
| DcBRAZIL09 | | | | G | H | G | | R | H | S |
| DcBRAZIL10 | | | | G | H | G | | R | H | S |
| DcBRAZIL11 | | | | G | H | G | | R | H | S |
| DcBRAZIL12 | | | | G | H | G | | R | H | S |
| DcBRAZIL13 | | | | G | H | G | | R | H | S |
| DcBRAZIL14 | | | | G | H | G | | R | H | S |
| DcBRAZIL15 | | | | G | H | G | | R | H | S |
| DcCOSTARICA1 | | | | G | H | G | | R | H | S |
| DcCOSTARICA2 | | | | G | H | G | | R | H | S |
| DcGUYANA01 | | | | G | H | G | | R | H | S |
| DcCOLUMBIA01 | | | | G | H | G | | R | H | S |
| DcCOLUMBIA02 | | | | G | H | G | | R | H | S |
| DcCOSTARICA3 | | | | G | H | G | | R | H | S |
| DcUSA09 | | | | G | H | G | | R | H | S |
| WdUSA01 | | | | | | | | | | |
| DcURUGUAY01 | | | | | | | | | | |
| WdCANADA01 | | Y | | | | | | | | |
| WdFRENCHGUIANA01 | | | | | | | | | | |
| WdFRENCHGUIANA02 | | | | | | | | | | |
| WdFRENCHGUIANA03 | | Y | | | | | A | | | |
| WdFRENCHGUIANA09 | I | | | | | | | | | |
| WdFRENCHGUIANA10 | | | | | | | | | | |
| WdFRENCHGUIANA11 | | | | | | | | | | |
| WdFRENCHGUIANA12 | | | | | | | | | | |
| WdFRENCHGUIANA13 | | | | | | | | | | |
| WdFRENCHGUIANA14 | I | | | | | | | | | |
| WdFRENCHGUIANA04 | | | | | | | | | | |
| WdFRENCHGUIANA15 | | | | | | | | | | |
| WdFRENCHGUIANA06 | | | | | | | | | | |
| WdFRENCHGUIANA07 | | | | | | | | | | |
| WdFRENCHGUIANA08 | | | | | | | | | | |
| WdFRENCHGUIANA16 | | Y | | | | | A | | | |
| WdUSA04 | | | | | | | | | | |
| WdFRENCHGUIANA17 | | | | | | | | | | |
| WdFRENCHGUIANA05 | | | | | | | | | | |

### SUPPLEMENTARY METHODS

#### Parasitic culture

Each cryopreserved strain from the BRC *Toxoplasma* was intraperitoneally inoculated into two out-bred female Swiss Webster (SW) mice (1 mL/mice). Animal experimentation conducted in Limoges was approved and accepted by the Ethics Committee for Animal Experimentation n°033 validated by the French Ministry of National Education, Higher Education and Research (Registration numbers: APAFIS#14582-2018041010294175 v2). Experimental procedures were conducted according to European guidelines for animal care (‘‘Journal Officiel des Communautés Européennes”, L358, December 18, 1986). All inoculated mice were monitored daily for clinical signs of toxoplasmosis during four weeks. Ill mice developing ascites were aseptically punctured for peritoneal exudates to collect live tachyzoites before being euthanized. Peritoneal exudates were washed with sterile saline solution (0.9% NaCl) at 1500 rpm for 10 minutes. After four weeks, surviving mice were tested for *T. gondii* antibodies by modified agglutination test (cut-off at 1:20 serum dilution). Seropositive mice were euthanized and brain samples were aseptically collected, rinsed in saline solution, placed in 1 ml of saline solution, and extruded through a 21-gauge needle several times, and then through a 23-gauge needle. Half of this suspension was treated by 1 ml of trypsin-EDTA solution (pre-heated at 37˚C), thoroughly shaken, and incubated at 37˚C for 3 minutes to disrupt tissue-cysts and liberate bradyzoites. The obtained suspension was then re-extruded through a 25-gauge needle several times, washed in 5 ml of Iscove’s Modified Dulbecco’s Medium (IMDM), and re-suspended in 1ml of IMDM. Each aseptically prepared mouse sample containing either tachyzoites or bradyzoites was inoculated in a Vero cell monolayer in a T175-flask. The culture medium was composed of IMDM treated with 1% of antibiotic saline solution (1000 U/ml penicillin and 100 µg streptomycin/ml in saline solution) and enriched with 2% of foetal bovine serum (FBS). Parasite growth was observed between one and five weeks post-initial inoculation but lasted between two weeks and 5 months. When parasitic growth was sufficient (> 40 tachyzoites at 40x magnification), the T-175 flask was vigorously shaken, the culture media collected and spun at 380 rpm for 5 min to pellet the large debris. The supernatant was then pelleted with a faster spin of 1500 rpm for 10 min. Then the supernatant was discarded, the tachyzoites pellet re-suspended with 2 ml of PBS free of Ca2+ and Mg2+, and the mix was homogenized through several passages in a 25-gauge needle, before adding 10 ml of PBS and homogenizing the mix with gentle shaking. The tachyzoite suspension was then filtrated through 3.0 micron polycarbonate filters, spun at 1500 rpm for 10 min, before reducing PBS volume to 1 ml and re-suspending the pellet in this volume. Lastly, the tachyzoite suspension was spun at 4500 rpm for 5 min and the volume reduced to 200 µl.

#### Mutation rate and generation time estimation

*Estimation of single nucleotide substitution per mitosis*

RH strain

The laboratory of Parasitology of the Limoges University Hospital Centre carried out *in vivo* culture of the RH strain (type I lineage) to produce *T. gondii* antigen for purposes of human diagnosis. This strain was maintained during 30 years (from January 1989 to march 11$^{th}$ 2019) through successive passages in outbred mice. This strain is virulent for the mouse as its growth cannot be controlled by its immune system, leading to the development of an ascites rich in tachyzoites in 48h post-inoculation. Three passages in mice were carried out each week; the ascites was punctured from infected mice and after being euthanized, the number of tachyzoites was counted, and 1,000,000 to 1,500,000 tachyzoites were again inoculated to two new mice.

In total, the RH strain was cultured *in vivo* during ~259,482 hours (~30 years). Assuming a division time of 5 hours (Radke et al., 2001), we calculated that ~ 51,896 mitotic divisions occurred during this period. In January 1989, tachyzoites were collected from an infected mouse, filtrated and stored at -80°C. The same protocol was followed for tachyzoites collected from an infected mouse in March 2019, with the addition of a step of cell culture to obtain more parasitic DNA. For these two samples, we applied the same protocol used for sequencing, variant calling and annotation of field isolates (with the exception of using per default mapping configuration in BWE). We determined the number of mutations that have accumulated over this period by comparing the two strains. Eighty-one SNPs were found across the 13 nuclear chromosomes (63,973,855 bases) following all filtration steps, and 67 SNPs were validated after manual curation on IGV 2.9.4.

*Estimation of the mutation rate of RH strain based on whole nuclear genome data:*

Mutation rate per mitosis: $\frac{67}{51{,}896} = 1.3\text{x}10^{-3}$

Mutation rate per mitosis per site: $\frac{67}{51{,}896 * 63{,}973{,}855} = 2.0\text{x}10^{-11}$

There were 25 exonic (18 missense, 2 nonsense and 5 silent), 20 intronic and 22 intergenic SNPs. This higher proportion of non-synonymous *vs* synonymous mutations (Missense / Silent ratio of 3.6) is consistent with selective constraints acting on RH strain, probably due to the unnatural conditions of maintenance of this strain for many generations. In order to mitigate bias in mutation rate associated to selection, we choses to calculate the mutation rate of RH strain from the SNPs found in the intergenic regions of the genome (23,896,470 nucleotides).

Estimation of the mutation rate of RH strain based on intergenic sequence data:

Mutation rate per mitosis: $\frac{22}{51{,}896} = 4.2\text{x}10^{-4}$

Mutation rate per mitosis per site: $\frac{22}{51{,}896 * 23{,}896{,}470} = 1.8\text{x}10^{-11}$

PRU strain

For comparison purposes, we also calculated the mutation rate of PRU strain (type II lineage). This strain was also maintained during 30 years, with only 75 passages in outbred mice (2.5 passages per year in average). This strain is not virulent to mice; after an acute phase characterized by rapid parasite multiplication, the parasites form dormant intracellular tissue cysts to evade the host's immune system (Djurković-Djaković et al., 2012). The acute phase of infection lasts ~21 days and can be divided into two phases (Jerome et al., 1998). First, tachyzoites carry out rapid multiplication during ten days. Tachyzoites of type II strains have a division time of 9 hours —an estimation made on ME49 strain (Radke et al., 2001)— and hence this stage involves ~27 mitoses. Then, tachyzoites differentiate into bradyzoites, which continue to multiply for 11 days, albeit at a slower rate. Bradyzoite division time has not been estimated for type II strains and we based our calculations on the estimate obtained from VEG strain (type III), which is of 15 hours (Jerome et al., 1998). Hence, this second stage of multiplication involves ~18 mitoses. Assuming a total number of 45 mitoses per passage, we calculated that 3,375 mitoses can occur during the 75 passages. We followed the same protocol as that used for RH to PRU for culture, sequencing and variant calling. We first identified 8 SNPs differentiating PRU-1989 from PRU-2019, a number than drops to 4 SNPs after manual curation in IGV.

*Estimation of the mutation rate of PRU strain based on whole nuclear genome data:*

Mutation rate per mitosis: $\frac{4}{3,375} = 1.2\text{x}10^{-3}$

Mutation rate per mitosis per site: $\frac{4}{3,375 * 63,973,855} = 1.8\text{x}10^{-11}$

The mutation rates of RH and PRU showed a high degree of agreement, which reinforces the robustness of our estimates. The RH strain was subjected to a much higher number of passages than the PRU strain (51,896 versus 3,375), and we therefore chose to base all subsequent calculations on the RH estimate for better precision.

*Estimation of single nucleotide substitution per year*

*Toxoplasma gondii* has a life cycle composed of three stages: in (1) the final host (cats), (2) the environment and (3) the intermediate host (mainly rodents). Several full life cycles can occur within one year.

The total number of DNA replications per year was calculated based on different assumptions about number of mitoses per stage and time to complete different stages of the life cycle.

When infected with one to few (~10) bradyzoites, cats excrete in total ~ 50 to 100 million oocysts (Dubey, 2001). Reaching this number of oocysts involves ~26 mitoses. Infected cats can excrete oocysts between the 3$^{d}$ and the 21$^{th}$ days following infection (Dubey et al., 1970; Elmore et al., 2010; Lappin, 2010), although most oocysts are shed between the 6$^{th}$ and the 13$^{th}$ days (Dubey, 2001).

In the **environment**, the newly secreted oocyst contains a single diploid sporoblast, that undergoes a meiotic division (2N–N), followed by a mitotic division, giving rise to eight haploid nuclei. Two DNA replications occur during this stage (Freppel et al., 2019). Oocysts sporulate in the environment in 48-72 hours after their excretion; their survival depends mainly on climatic conditions, and is favoured by humidity. Although sporulated *T. gondii* oocysts can survive in the environment for 1.5 years, the median survival time varies between 27 days under dry conditions and 84 days under damp (ideal) conditions (Lélu et al., 2012).

Sporulated oocysts ingested by an **intermediate host** differentiate into rapidly replicating tachyzoites during the first 10 days of infection (Jerome et al., 1998). Tachyzoites division time varies according to the parasitic strains (Jerome et al., 1998; Radke et al., 2001): 5 hours for RH (type I), 9 hours for ME49 (type II) and 6 hours for VEG (type III). Hence, this first stage of multiplication in the intermediate host involves a minimum of 27 mitoses (assuming a division time of 9 hours) and a maximum of 48 mitoses (assuming a division time of 5 hours). The tachyzoites then differentiate into slow replicating bradyzoites which continue to actively replicate until the end of the third week (Djurković-Djaković et al., 2012). Bradyzoite division time is 15 hours (Jerome et al., 1998), an estimate obtained from VEG strain (type III). Hence, this second stage of multiplication in the intermediate host involves ~18 mitoses.

The length of this stage depends on the period between the infection time and time of death. Rodent lifespans vary by species and can reach several years for some species: voles and house mice, which are common cat prey, have a median lifespan of 60-90 days (Naughton, 2012) and 90-120 days (Phelan and Austad, 1989), respectively. Infection can occur at any time from weaning age (~21 days (König and Markl, 1987; Solomon, 1991)) until death. Thus, we assume on average that *T. gondii* is likely to infect rodent for:

Lower assumption: $\frac{60-21}{2} \approx 20days$ Higher assumption: $\frac{120-21}{2} \approx 50days$

*Summary table of the duration in days of the life cycle stages*

| stages | minimum | lower assumption | higher assumption | maximum |
|---|---|---|---|---|
| final host | 3 | 6 | 13 | 21 |
| environment | 2 | 27 | 87 | 547 |
| intermediate host | 1 | 20 | 50 | *Hundreds* |
| **full life cycle** | **6** | **53** | **150** | ***Hundreds*** |

By summing the estimates for the three stages, we get an estimate of 73 to 94 mitoses per full life cycle and of 2.4 to 6.9 life cycles per year. We used these estimates to calculate the mutation rate of *T. gondii* per life cycle and per year:

Mutation rate per full life cycle:

$(4.2x10^{-4}$ x 73) to $(4.2x10^{-4}$ x 94) => 0.03 to 0.04

Mutation rate per full life cycle per site:

$(1.8x10^{-11}$ x 73) to $(1.8x10^{-11}$ x 94) => $1.3x10^{-9}$ to $1.7x10^{-9}$

Mutation rate per year:

(0.03x 2.4) to (0.04x 6.9)  =>  0.07 to 0.28

Mutation rate per year per site:

$(1.3 \times 10^{-9} \times 2.4)$ to $(1.7 \times 10^{-9} \times 6.9)$  =>  $3.1 \times 10^{-9}$ to $11.7 \times 10^{-9}$

**Identifying clonal lineages.**

Strict clonal multiplication in a population is characterized by gradual accumulation of random mutations over generations (Milgroom, 2015). At the opposite, recombination with a distinct population introduces highly divergent genomic regions, leading to a sharp rise in genetic distances between individuals. The `mlg.filter` function of *poppr* R package exploits this property by enabling to define a threshold that delimits genetic distances resulting from the gradual accumulation of random mutations (small genetic distances) within a population from genetic distances resulting from recombination with a divergent population (large genetic distances). This threshold is expected to correspond to the maximal genetic distance resulting only from mutation accumulation (within a clonal lineage), and above which genetic distances are rather explained by recombination. The function `mlg.filter` gradually collapses genomes based on genetic distance (along the horizontal axis) using three different clustering algorithms in order to define clonal lineages boundaries. A sharp drop in the number of uncollapsed genomes is observed when collapsing genomes separated by a genetic distance < 0.01 (Supplementary Fig. 5). At this threshold value, all strains separated by small genetic distances are collapsed together; we consider that they belong to the same clonal lineage. This threshold was used to generate a minimum spanning network (MSN), in which strains of the same putative clonal lineage were collapsed in single circles. Overall, few mismatches between MS-defined and genome-defined lineages were noticed, as only two genomes did not cluster with the other strains of their respective MS-defined lineages: DcMARTINIQUE01 (type II) and DcGABON08 (Africa 1). To verify that *poppr*-defined lineages are true clonal lineages, we generated plots of SNPs density for each of the main lineages (type I, type II, type III, Africa 1 and Africa 4) by dividing the genome into 10kb windows (Supplementary Fig. 6). We expected to observe an even genome-wide density of SNPs in the case of strict clonality, and to observe sharp variations in the density of SNPs (designated as recombination break points) if one or more strains have inherited divergent genomic sequences following recombination with another lineage or population (Graham et al., 2005). We found repeated sharp variations in SNPs densities in plots produced for type III and Africa 4 lineages, a pattern usually observed when recombination break points are disrupting chromosomal ancestry along one or more genomes. We calculated pairwise SNP distances between the genomes of each of these two lineages. Type III genomes (n=19) showed a maximum of 350 SNPs differences, with the exception of DcUSA04 (M7741) that showed 10,077 to 10,459 SNP differences with other type III genomes. Among Africa 4 genomes (n=3), DcSENEGAL04 showed 9,820 SNP differences with DcSENEGAL14 and DcSENEGAL23, the latter two showing no SNP differences. We therefore excluded the divergent genomes from their respective *poppr*-defined lineages —which are likely the products of a recombination with a strain of a distinct population— and generated new SNPs density plots .The sharp variations in SNPs densities previously observed did not recur in the new plots (data not shown), indicating that the excluded genomes had divergent ancestry in certain chromosomal portions. The other lineages (type I, type II and Africa 1) exhibited an even distribution of SNPs across their genomes, albeit type II had a much higher SNPs density compared to the other lineages (Supplementary Table 8).

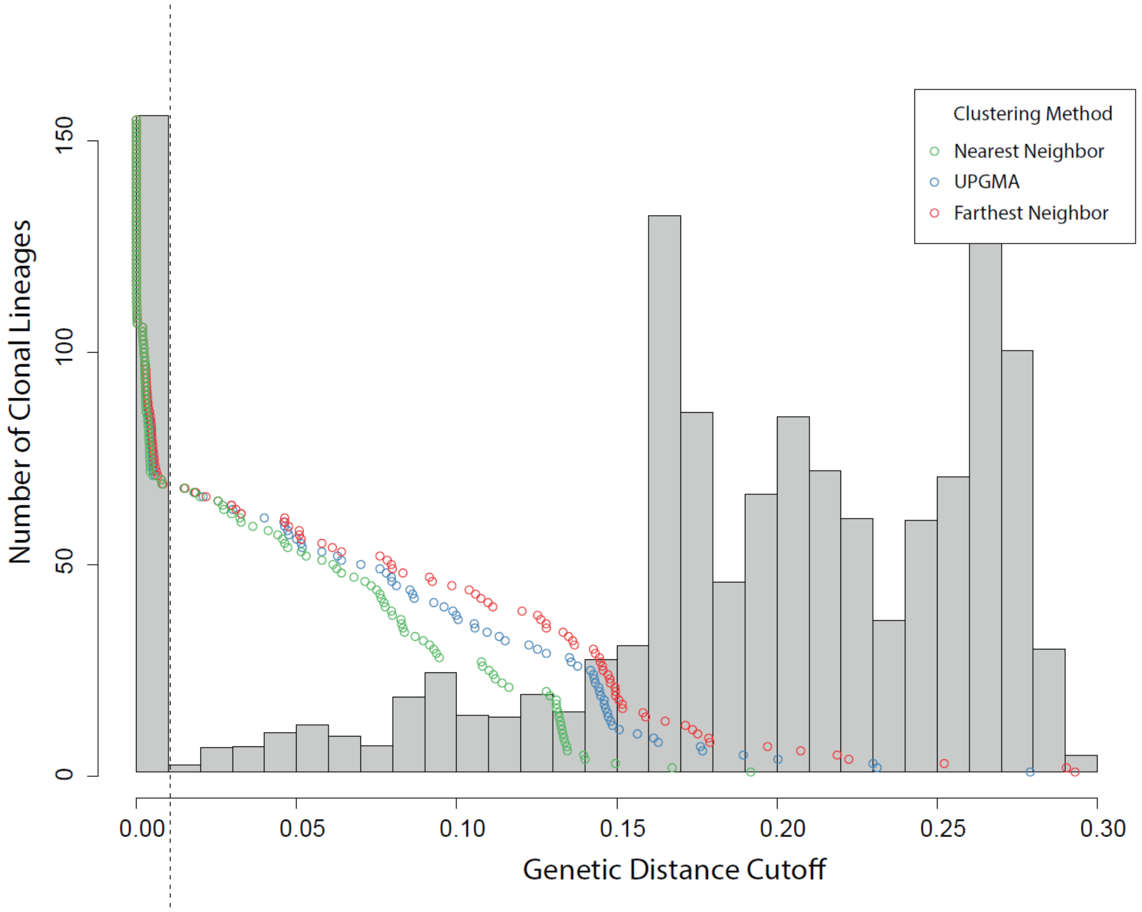

**Supplementary Fig. 5. Graphical representation of three different clustering algorithms collapsing the 156 *Toxoplasma gondii* genomes into clonal lineages.** The horizontal axis is the genetic distance based on a dissimilarity matrix as calculated in *poppr* R package. The vertical axis represents the number of lineages observed. Each point shows the threshold at which one would observe a given number of uncollapsed groups or individuals. The vertical dashed line marks the threshold used to collapse the 156 genomes into 10 clonal lineages and 59 non-clonal strains.

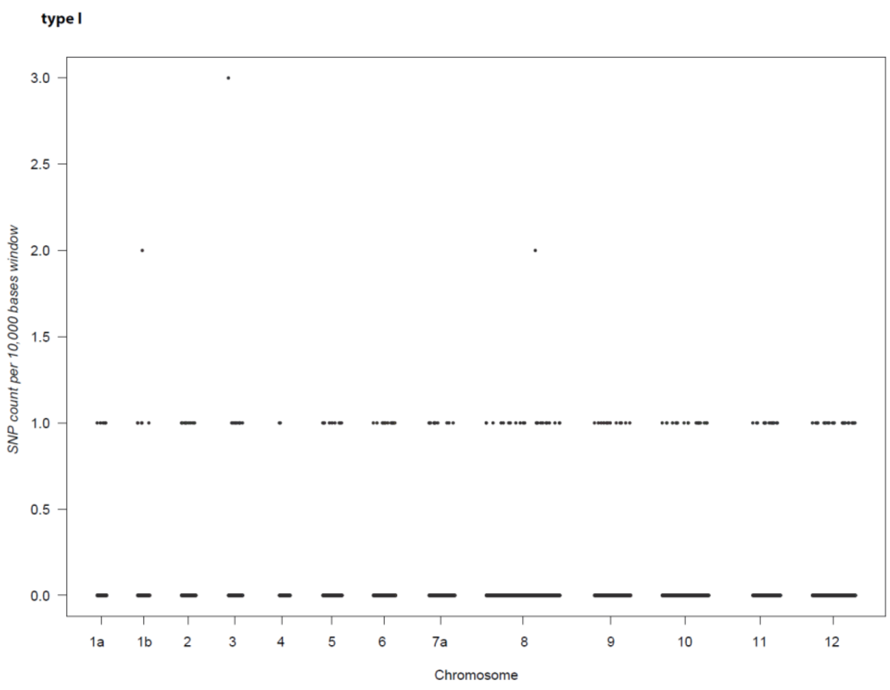

***Supplementary Fig. 6a.***

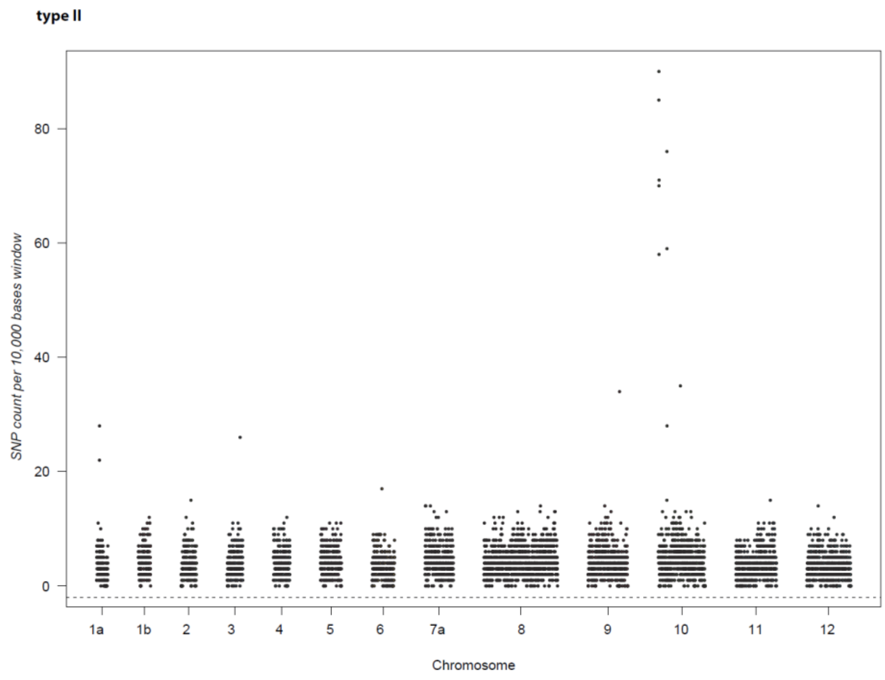

***Supplementary Fig. 6b.***

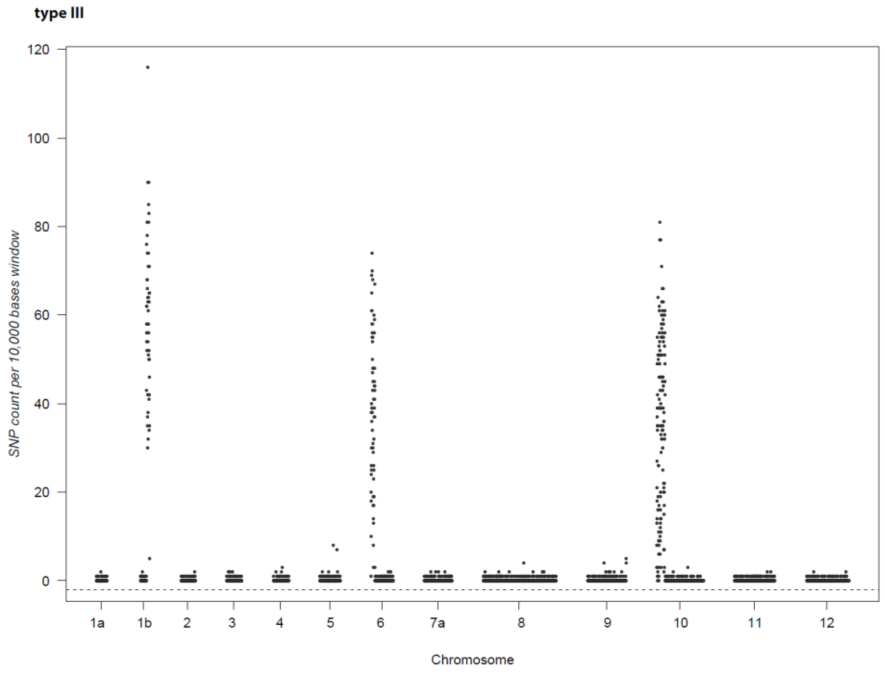

***Supplementary Fig. 6d.***

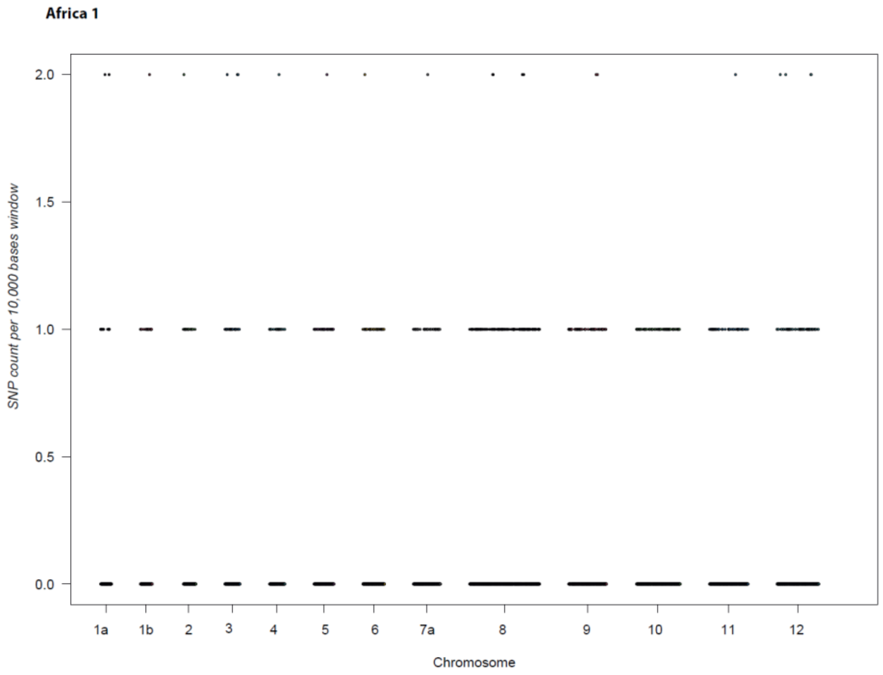

***Supplementary Fig. 6d.***

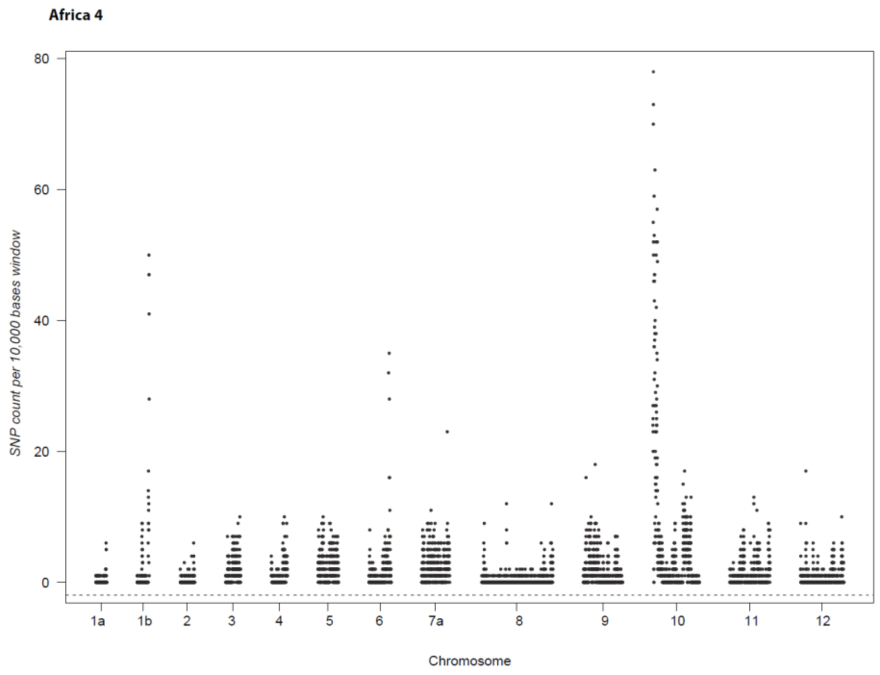

***Supplementary Fig. 6e.***

**Supplementary Fig. 6. SNPs density per window comparing strains of each intercontinental *poppr*-defined clonal lineages.** Plots were generated for each of the five intercontinental lineages: type I (a), type II (b), type III (c), Africa 1 (d) and Africa 4 (e).The total number of SNPs per 10 kb window for all strains within a lineage is plotted along the y-axis. Chromosome numbers are indicated along the x-axis.

**Supplementary Table 8. Genetic diversity of the most common intercontinental clonal lineages**

| | Number of strains | π diversity* | SNP distance between the two most divergent strains within the clonal lineage with filtering out singletons | SNP distance between the two most divergent strains within the clonal lineage without filtering out singletons |
|---|---|---|---|---|
| Type I | 3 | 5.6x10-5 | 160 | 502 |
| Type II | 48 | 1.1x10-4 | 8,408 | 9,116 |
| Type III | 18 | 1.7x10-5 | 350 | 457 |
| Africa 1 | 11 | 2.6x10-5 | 453 | 489 |

*Average nucleotide diversity calculated from the division of the genome into 10kb-windows (VCFtools)

The Variant Call Format and VCFtools, Petr Danecek, Adam Auton, Goncalo Abecasis, Cornelis A. Albers, Eric Banks, Mark A. DePristo, Robert Handsaker, Gerton Lunter, Gabor Marth, Stephen T. Sherry, Gilean McVean, Richard Durbin and 1000 Genomes Project Analysis Group, Bioinformatics, 2011
